# Speech perception under the tent: A domain-general predictive role for the cerebellum

**DOI:** 10.1101/2020.06.05.136804

**Authors:** Jeremy I Skipper, Daniel R Lametti

## Abstract

The role of the cerebellum in speech perception remains a mystery. Given its uniform architecture, we tested the hypothesis that it implements a domain-general mechanism whose role in speech is determined by connectivity. We collated all neuroimaging studies reporting cerebellar activity in the Neurosynth database (n = 8,206). From this set, we found all studies involving passive speech and sound perception (n = 72, 64% speech, 12.5% sounds, 12.5% music, and 11% tones) and speech production and articulation (n = 175). Standard and coactivation neuroimaging meta-analyses were used to compare cerebellar and associated cortical activations between passive perception and production. We found distinct regions of perception-and production-related activity in the cerebellum and regions of perception-production overlap. Each of these regions had distinct patterns of cortico-cerebellar connectivity. To test for domain generality versus specificity, we identified all psychological and task-related terms in the Neurosynth database that predicted activity in cerebellar regions associated with passive perception and production. Regions in the cerebellum activated by speech perception were associated with domain-general terms related to prediction. One hallmark of predictive processing is metabolic savings (i.e., decreases in neural activity when events are predicted). To test the hypothesis that the cerebellum plays a predictive role in speech perception, we examined cortical activation between studies reporting cerebellar activation and those without cerebellar activation during speech perception. When the cerebellum was active during speech perception there was far less cortical activation than when it was inactive. The results suggest that the cerebellum implements a domain-general mechanism related to prediction during speech perception.

## Introduction

The cerebellum is a remarkable structure, having about 80% of the neurons in the brain but only around 10% of its mass (Herculano-Houzel et al., 2015). Compared to other primates, it is significantly larger in humans relative to the size of the neocortex. This expansion may reflect the need for complex motor programs associated with tool use and speech, hallmarks of human evolution (Barton & Venditti, 2017; MacLeod et al., 2003). Indeed, the lateral cerebellum scales up with mammals that learn vocally, including elephants, humans, seals, dolphins and whales (Smaers et al., 2018). Consistent with more expansion in humans and its role in vocal learning, the cerebellum plays an important role in speech production (Ackermann et al., 2007). This is also consistent with the view since the early 1800s that the cerebellum is primarily a motor structure, considered the organ of sexuality by Gall, with Rolando and Flourens providing the first evidence for its more general role in motor function (Glickstein et al., 2009; Macklis & Macklis, 1992). Over the last 50 years, it has also become apparent that the cerebellum plays *some* role in ‘nonmotor’ language domains. Activity in the cerebellum is observed during lower-level auditory functions, like speech timing and phonology and higher-level tasks involving semantics, grammar and comprehension (Ackermann & Brendel, 2016; Mariën & Borgatti, 2018; Mariën & Manto, 2015). However, what the cerebellum contributes to these tasks remains, to quote one ‘consensus’ paper, ‘an ongoing enigma’ (Mariën et al., 2014). Here we address questions about the role of the cerebellum during speech perception.

### Domain-generality/Prediction

Though the evidence has remained elusive, some theories claim that the function of the cerebellum is domain-general (Diedrichsen et al., 2019). Thus, the contribution (or computation) contributed by the cerebellum to vocal learning and speech production would be similar to that contributed to speech perception. This idea is captured by the Universal Cerebellar Transform (UCT) theory (Schmahmann, 2019; Schmahmann et al., 2019), which maintains that, because the cerebellum has a relatively homogenous architecture, with repeating corticonuclear micro complexes (Eccles et al., 1967), it performs a ‘consistent’ computation. Any differences in what this computation contributes to would be determined by variations in cerebellar location and corresponding cortico-cerebellar connectivity.

There have been a number of proposals as to the nature of the domain-general computation in the cerebellum. One suggestion from the motor control literature is that it implements internal models specifically and prediction more generally (Ito, 2008; Popa & Ebner, 2018; Siman-Tov et al., 2019; Taylor & Ivry, 2014; Wolpert et al., 1998). Internal models are neural representations of an organism’s interactions with the world. These can be used to predict the sensory consequences of movements and maintain movement accuracy. When differences between predicted and actual sensory states occur, internal models are updated such that, on subsequent movements, the discrepancy between predicted and actual sensory feedback is reduced (Lametti et al., 2018; Shadmehr et al., 2010; Wolpert et al., 2011). In evidence that the cerebellum plays a key role in this process, participants with cerebellar disruption, due to stroke or brain stimulation, show slow or altered learning in a wide-range of tasks including movement adaptation in response to visual alterations of the limbs (Baizer & Glickstein, 1974; Martin et al., 1996; Morton & Bastian, 2004) and following physical perturbations of movement (Gibo et al., 2013; Rabe et al., 2009; Smith & Shadmehr, 2005).

Similarly, the cerebellum seems to play a key role in the predictive processing required for the maintenance of accurate speech production. Patients with cerebellar damage frequently present with a range of speech production deficits (Ackermann et al., 2007). They also exhibit impaired feedforward control of speech movements (i.e., an impaired ability to update internal models)(Parrell et al., 2017). In healthy participants, altering the cerebellum with noninvasive brain stimulation has been shown to alter sensorimotor adaptation during speech production (Lametti et al., 2017). A similar result was observed during sensorimotor adaptation associated with limb movements (Galea et al., 2011; Jayaram et al., 2011), although the impact of cerebellar tDCS on sensorimotor adaptation can be inconsistent (Jalali et al., 2017). Linking the aforementioned limb and speech adaptation literature, a recent meta-analysis found that sensory feedback manipulations resulting in adaptation were more likely to activate the cerebellum (Johnson et al., 2019).

### Speech perception

Thus, according to UCT-like theories, the contribution of the cerebellum to motor control should extend to non-motor domains. Does the cerebellum contribute predictions to speech perception (Moberget & Ivry, 2016)? Despite much theorising (Ackermann, 2008; Ackermann et al., 2007; Argyropoulos, 2016; Callan et al., 2007; Hertrich et al., 2016; Mariën et al., 2014; Mariën & Manto, 2015; Moberget & Ivry, 2016; Schwartze & Kotz, 2016), that question is difficult to answer because the associated neurobiological research is sparse. *Pubmed* (queried March 2021) lists *six* articles with ‘cerebellum’ or ‘cerebellar’ and ‘speech perception’ in the title, half of which are review articles (included in the references in the prior sentence). And yet, there are thousands of neuroimaging studies involving speech perception that report cerebellar activity (as we later show).

#### Domain-generality

Explicit evidence for domain-generality of the cerebellum for speech perception does not exist (because speech perception studies by definition only study speech). Nonetheless, a number of task battery and meta-analyses studies more or less address this topic. In two studies, participants did task batteries, some of which were language related (Guell, Gabrieli, et al., 2018; King et al., 2019). The conclusions from one of these is that, despite overlap between language and social cognition tasks, the cerebellum represents cognitive functions in a domain-specific manner because of the spatially modular appearance of these different functions (Guell, Gabrieli, et al., 2018). Though the other task-battery study does not explicitly make this claim, it also shows specific functions mapped to specific cerebellar regions, suggesting specificity (Diedrichsen et al., 2019; King et al., 2019). Five meta-analyses support the idea that the cerebellum plays specific roles in audition, speech perception and language comprehension (Balsters et al., 2014; E et al., 2014; Petacchi et al., 2005; Riedel et al., 2015; Stoodley & Schmahmann, 2009). Two of these use a large number of studies to profile clusters of cerebellar ROIs in terms of associated behavioural domains (Balsters et al., 2014; Riedel et al., 2015). Though some of these were associated with speech and language, they were also associated with a range of other domains and subdomains. Though domain-generality is not discussed, the breadth of tasks related to specific regions of cerebellar activity seems at odds with a domain-specific account suggested by task battery studies.

#### Prediction

Prediction likely plays an important role in speech perception. Due to differences in vocal-tract lengths, accents and speaking contexts, the acoustics of identical phonemes can vary considerably. To help solve this ‘lack of invariance’ problem, it has long been noted that the brain uses visual and linguistic information to predict forthcoming auditory information, constraining the interpretation of acoustic signals (Ganong, 1980; Holt & Lotto, 2002; Ladefoged & Broadbent, 1957; McGurk & MacDonald, 1976; Sjerps et al., 2011; Skipper, 2014; Skipper et al., 2007). This predictive process involves a wide array of ‘motor’ regions also involved in speech production, though most of this work has only examined cortical motor regions (Skipper, 2015; Skipper et al., 2005, 2017).

The majority of cerebellar studies pertaining to prediction have focused on ‘higher level’ linguistic prediction (e.g., involving word meaning)(Argyropoulos, 2016; D’Mello et al., 2017; Lesage et al., 2012, 2017; Moberget et al., 2014; Pleger & Timmann, 2018; Sheu et al., 2019). Only a small number of studies have specifically investigated the role of the cerebellum in speech perception, regardless of its role in prediction. Some of these studies show that the cerebellum seems to contribute timing signals to the unfolding process of speech perception. Patients with cerebellar damage show impairments in the perception of speech-sound contrasts distinguished by the purely durational cue occlusion time (e.g., the German words ‘boten’ vs ‘boden’)(Ackermann et al., 1997). These patients do not show impairments in the identification of speech sound contrasts that can be distinguished by both durational and non-durational cues such as voice onset time (Ivry & Gopal, 1993; Repp, 1979). These two findings were later explored with fMRI and the right cerebellum was linked to the durational characteristics of speech sounds (Guediche et al., 2015; Lametti et al., 2016; Mathiak et al., 2002). More generally, timing signals are clearly required for prediction (Kotz & Schwartze, 2010). Converging lesion, neuroimaging, and brain stimulation results link the cerebellum to both timing and predictive processing in speech perception (Guediche et al., 2015; Knolle et al., 2012; Kotz et al., 2014; Lametti et al., 2016; Moberget & Ivry, 2016; Schwartze et al., 2016; Schwartze & Kotz, 2013, 2016).

#### Tasks

Existing cerebellar speech perception studies tend to involve tasks that lack ecological validity and involve motor responses, limiting claims about domain-generality and prediction. In terms of validity, the tasks used in most studies are not particularly representative of natural speech perception. For example, in one task battery study, ‘language’ cerebellar regions are defined by activity associated with listening to and answering questions about Aesop’s Fables *subtracted* from activity associated with reading math problems and selecting the correct answer from two alternatives (Guell, Gabrieli, et al., 2018). In another task battery study, language processing seems to be defined using tasks like verbal working memory with letters, verb generation versus reading and/or a two alternative forced choice semantic task following sequential reading of five words. Similarly, the tasks used in the five previously mentioned meta-analyses mostly involved single word generation, repetition, reading or making semantic decisions (E et al., 2014; Petacchi et al., 2005; Stoodley & Schmahmann, 2009).

Perhaps more problematic, most tasks used in these studies required participants to either read, leading to subvocal speech production, or make a meta-linguistic judgement as indicated by a button response. Though studies often include task subtractions meant to control for motor engagement, it cannot be reasonably demonstrated that this was achieved (Friston et al., 1996; Poeppel, 1996). Similarly, studies having motor responses on discarded trials could still cause motor activation associated with participants’ expectations that they will need to make responses. This is an important oversight given the historical perspective that the cerebellum is predominantly a motor structure.

### Hypotheses

Theory and a small amount of empirical work suggests that the cerebellum might make a domain-general predictive contribution to speech perception similar to the contribution it makes to speech production. There are, however, no studies that directly address both domain-generality and predictive processing. Task-battery and meta-analyses studies suggest mixed results about domain-generality, while studies of prediction during speech perception can likely be counted on one hand and do not address domain-generality. Furthermore, there are few studies of speech perception and the cerebellum that examine natural speech perception i.e., speech perception in the complete absence of movement. To begin to address these gaps in the literature, we performed cerebellar meta-analyses, coactivation based meta-analyses and text-based functional profile analyses. Critically, we used a large number of studies that involve only ‘passive’ speech perception without an overt motor response on any trial. As a minority of studies, we included passive perception studies involving tone and nonspeech sound stimuli (e.g., instrumental music) because many languages are tonal (Yip, 2002) and nonspeech sounds activate cortical areas associated with speech perception (Peretz et al., 2015). We compared these passive perception studies to studies involving speech production and articulation (Figure 1 presents a schematic overview of this work).

**Figure 1.**
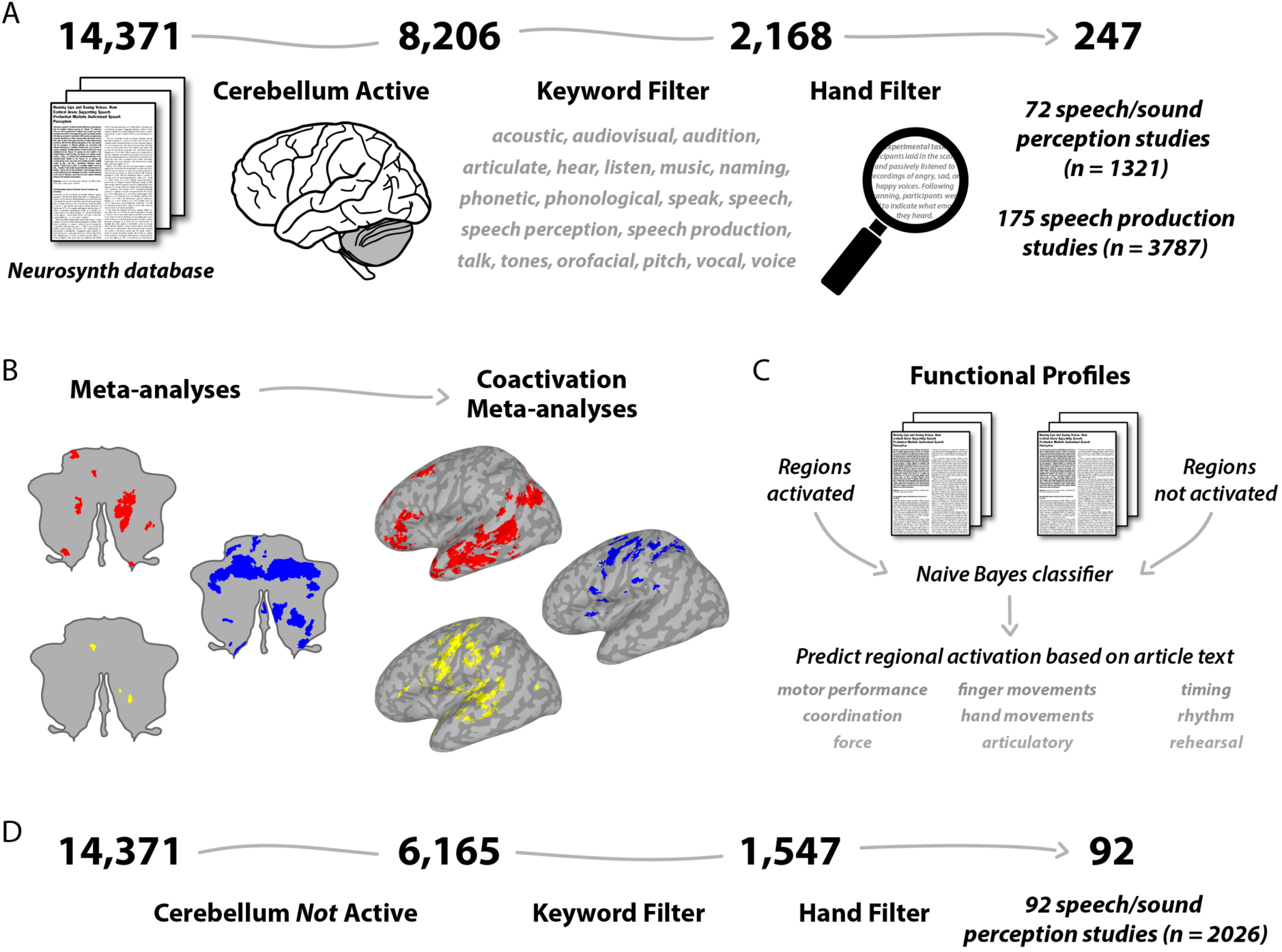
General procedure used for meta-analyses. A) All articles in the Neurosynth database reporting activity in a cerebellar mask were found. These were filtered by speech and language-related keywords and by hand to identify studies involving natural (i.e., ‘passive’) sound/speech perception and speech production. B) Meta-analyses were run using the identified studies to locate regions of activity in the cerebellum associated with speech perception (red), speech production (blue) and both tasks (yellow). Coactivation meta-analysis was then used to determine which other sub/cortical regions were functionally connected to these sets of regions. C) A classifier was trained to predict articles reporting cerebellar activity in the identified sets of speech perception, production and overlap regions based on keywords in the articles. D) In a second round of article selection, all articles in the Neurosynth database involving natural speech perception that did not report cerebellar activity were found.

Our first hypothesis was that the cerebellum plays a domain-general role in speech perception—that is, it makes a contribution to speech perception that is not inherently speech specific. Rather, any speech specificity partly derives from connectivity patterns that give the cerebellum a modular topological appearance. To test this, we examined regions of activity in the cerebellum related to speech perception and production. Speech perception and production are both sensorimotor processes that share subprocesses, but they are also distinct in important ways (e.g., production involves overt articulation). Thus, we anticipated a mix of overlapping and distinct activity patterns in the cerebellum reflecting the shared and unique components of perception and production. Using coactivation meta-analysis, we predicted that networks originating from speech perception, production and overlapping regions would have different cortical connectivity. To test for domain-generality, we analysed task-related terms mined from the abstracts of thousands of neuroimaging studies to see which of these predicted activity in cerebellar regions associated with speech perception, production or their overlap. We expected that these regions would also be associated with a wide range of other tasks that are not speech or domain-specific.

Our second hypothesis was that the domain-general role played by the cerebellum and its connections during speech perception is related to prediction. To assess this, we tested a fundamental tenet of predictive models, that prediction results in metabolic savings because the brain has to do less processing when predictions are accurate and, correspondingly, more processing for unexpected acoustic information (Moberget et al., 2014; Skipper, 2014). We compared speech perception-related whole-brain activity between studies reporting cerebellar activity to whole-brain activity in studies without reported cerebellar activity. We hypothesized that, if the cerebellum is involved in prediction during natural speech perception, there should be a greater amount of activity throughout the brain when the cerebellum is *not* active during this task.

## Methods

### Article selection

Figure 1A outlines the article selection steps. First, we created a maximum probability mask of the cerebellum using a probabilistic cerebellar atlas (Diedrichsen et al., 2009). The latter was created by averaging the cerebellar lobule masks from 20 participants, aligned to the MNI152 template by nonlinear registration (Diedrichsen et al., 2009). We then found all of the published articles that had activity somewhere in this mask and that appeared in the Neurosynth database (version 0.7, released July, 2018; https://github.com/neurosynth/neurosynth-data) (Yarkoni et al., 2011). This version contains 507,891 activation peaks or centres of mass from 14,371 studies with over 3,200 term-based features. The intersection of the cerebellum mask and database resulted in 8,206 articles (57% of the Neurosynth database).

Next, we searched Pubmed (October, 2018) for all articles that matched a set of 20 search terms (acoustic, audiovisual, audition, articulate, hear, listen, music, naming, phonetic, phonological, speak, speech, speech perception, speech production, talk, tones, orofacial, pitch, vocal, voice) and their variants (e.g., articulate, articulators, articulation, articulatory) and a set of eight Medical Subject Headings (MeSH) terms, a controlled vocabulary used by Pubmed for indexing life science articles (auditory perception, language, verbal behaviour, hearing, hearing tests, speech, speech acoustics, speech production measurements). This search returned 1,002,940 articles. We then found the intersection of these articles and the 8,206 articles in the Neurosynth database reporting cerebellar activity. This resulted in 2,168 articles with cerebellum activity that potentially involved speech, language and/or articulation (15% of the Neurosynth database).

We then went through the abstract and methods of these 2,168 articles by hand to determine if they included a 1) natural speech perception task (i.e., passive speech/sound/music perception that simply involved listening without another explicit task) or 2) a speech production task (i.e., speaking overtly/covertly or moving the articulators). We required that a number of criteria be met for studies to be included. In particular, studies that focused on reading, used patient populations, tested participants younger than 18 or focused on resting state analyses were excluded. Critically, perception studies that involved *any* motor response no matter how minor (e.g., a button press on 5% of trials to maintain alertness) were *not* included. Studies that involved the passive perception of tones and nonspeech sounds (e.g., instrumental music) were included in the analysis as a minority of studies. This decision was made for the following reasons: By some estimates, 60-70% of the world’s languages are tonal (Yip, 2002), tones can be produced by the human vocal tract and they are (arguably) similar to phonemes. Converging evidence from fMRI and direct neural recordings suggests that there’s overlap in cortical activity patterns associated with speech and music listening (Peretz et al., 2015). There’s also behavioural evidence that music and language processing draw on a shared resource (Kunert & Slevc, 2015). More generally, the basic units of speech are unknown, and it is unclear when sound perception changes to speech perception (Bybee & McClelland, 2005; Goldinger & Azuma, 2003; Lotto & Holt, 2000; Skipper et al., 2017). Of the original 2,168 articles, 72 (3.32% or 0.50% of the full Neurosynth database; n = 1321 participants) were natural speech/sound perception studies (64% speech, 12.5% sounds, 12.5% instrumental music and 11% tones) and 175 (8.07% or 1.22%; n = 3787 participants) involved speech production or articulation. See Tables S1 and S2 for a complete list of speech perception and production articles used in meta-analyses.

### Cerebellar meta-analyses

We used Neurosynth to conduct meta-analysis on our sample of natural speech perception and production studies that activate the cerebellum. Neurosynth is a database and tool for performing term-based meta-analysis (Yarkoni et al., 2011). As designed, it uses a form of kernel density analysis to compare activations reported in studies that frequently use selected psychological terms (e.g., ‘language’, ‘working memory’) to activations reported in studies in the rest of the database that do not use these terms. Instead of using Neurosynth to perform a term-based meta-analysis, we simply provided it with the articles found to involve natural speech perception or speech production. Neurosynth compared activations reported in the provided studies to activations reported in the rest of the Neurosynth database. The resulting cerebellum activity maps reflect activations that occur more consistently in our two samples as compared to other studies We examined baseline contrasts and overlaps. For baseline contrasts we used an FDR corrected threshold of q < .01 across the whole brain. We examined speech perception and production overlaps at the same individual FDR corrected thresholds. For added protection, we also required that cluster sizes be greater than 10 voxels. Results are displayed on a cerebellar flatmap using version 3.4 of the SUIT Matlab toolbox (Diedrichsen & Zotow, 2015).

### Cerebellar coactivation meta-analysis

We next did a meta-analytic coactivation analysis from the regions unique to speech perception, production and their overlap across all 14,371 neuroimaging studies in the Neurosynth database (Figure 1B). This analysis assumes that if a cerebellar region frequently coactivates with other brain regions across many studies and statistical contrasts then that region can be considered to be part of a network with the co-active brain regions. The principle here was to perform a formal contrast between studies that activate each of the three sets of regions as compared to studies that tend to activate the other sets of regions. The resulting statistical maps identify voxels throughout the brain that have a greater probability of coactivating with the identified regions. A two-way chi-square (χ2) test was used to calculate p-values for each voxel between the sets of studies. The resulting images were again thresholded using an FDR of q < .01. We again required that cluster sizes be greater than 10 voxels. This analysis and functional profile analyses (discussed in the next section) were based on de la Vega et al. (2018) and Vega et al. (2016) (https://github.com/adelavega/neurosynth-mfc/ and https://github.com/adelavega/neurosynth-lfc).

To provide descriptive functional labels of the resulting coactivation patterns, we calculated the Pearson correlation of each vectorized coactivation map with meta-analyses available in the Neurosynth database. This resulted in *r-values* that reflect the spatial similarity between each coactivation map and other large-scale meta-analysis (see Table 1).

**Table 1.** The perception, production and overlap networks in Figure 3 were individually correlated with whole brain term based meta-analyses. The following table contains the top 20 correlations, showing that these networks are neither speech or domain-specific. Given the number of voxels in each correlation, these are roughly Bonferroni corrected at a *p-value* of .01/(20*3) < 0.0001.

| Perception |  | Production |  | Overlap |  |
| --- | --- | --- | --- | --- | --- |
| Meta-Analysis | Correlation | Meta-Analysis | Correlation | Meta-Analysis | Correlation |
| vi | 0.41 | finger movements | 0.42 | cerebellar | 0.52 |
| cerebellar | 0.38 | motor | 0.41 | cerebellum | 0.52 |
| lobules | 0.35 | execution | 0.41 | vi | 0.51 |
| cerebellum | 0.34 | premotor | 0.40 | lobules | 0.46 |
| mind | 0.27 | finger | 0.40 | vermis | 0.34 |
| theory mind | 0.26 | tapping | 0.39 | nuclei | 0.24 |
| mind tom | 0.26 | premotor cortex | 0.38 | coordination | 0.23 |
| tom | 0.25 | cerebellum | 0.38 | motor | 0.22 |
| mental states | 0.23 | movements | 0.38 | production | 0.21 |
| vermis | 0.22 | supplementary | 0.35 | speech production | 0.20 |
| theory | 0.22 | finger tapping | 0.35 | hemispheres | 0.19 |
| mentalizing | 0.20 | motor imagery | 0.34 | force | 0.19 |
| autobiographical | 0.19 | movement | 0.34 | cortex cerebellum | 0.19 |
| comprehension | 0.19 | supplementary motor | 0.34 | finger | 0.19 |
| nuclei | 0.18 | hand | 0.33 | motor control | 0.19 |
| sentence | 0.15 | sequential | 0.31 | movement | 0.18 |
| sentences | 0.15 | cerebellar | 0.31 | finger tapping | 0.18 |
| thinking | 0.14 | imagery | 0.29 | tapping | 0.18 |
| semantic | 0.13 | dorsal premotor | 0.29 | loop | 0.17 |
| person | 0.13 | handed | 0.28 | sensorimotor | 0.17 |

### Cerebellar functional profile analyses

To test for domain-specificity or generality, we next generated functional profiles of the activity patterns in each of the speech perception, production and overlap regions (Figure 1C). This was done by determining which of the terms in the Neurosynth database (which are mined from the text of the 14,371 abstracts) best predicted activity in each of the three sets of cerebellum regions. Specifically, this analysis determines whether a classifier could predict if a study activated specific perception, production or overlap regions in the cerebellum given the terms mentioned in the study’s title/abstract.

A naive Bayes classifier was trained to discriminate three sets of high frequency terms associated with activation in each set of regions versus a set of studies that did not produce activation in those regions. Fourfold cross-validation was used for testing and the mean score was calculated across all folds as a summary measure of performance. Models were scored using accuracy, or the fraction of samples correctly predicted. The log odds-ratio (LOR), the probability that a term is present in active versus inactive studies, from the naive Bayes models from each set of regions was used to generate the functional profiles. The LOR indicates whether a term is predictive of activation in a given cerebellar set of regions. We output the terms with LORs that predicted activation in each of the sets of cerebellar regions at an uncorrected statistical threshold of p < .05. To conduct functional profile analyses, we went through the terms by hand and labelled each as either anatomical, fMRI or task related. The anatomical label included any term related to brain anatomy (e.g., ‘cerebellum’); the fMRI label included any terms related to the fMRI signal (e.g., ‘bold signal’), stimuli (e.g., ‘video’) and methods (e.g., ‘contrasted’); and the task label was given to any task-related terms (e.g., ‘finger tapping’) and their associated functions (e.g., ‘speech production’).

We then created four further groups of terms. First, to validate the term-based approach, we labelled terms as confirmatory if they were specifically related to the cerebellum, natural/passive speech perception or speech production. Second, to determine whether our perception, production and overlap regions were actually speech specific, we labelled each term as to whether it was remotely speech related or whether it had no obvious relationship to speech. Third, to more generally examine the domain-specificity of regions, we labelled each term with the four gross psychological domains: perceptual, motor, cognitive and social/emotional. Anything that did not fit into these categories was labelled as nonspecific. Finally, to provide some insight as to what general functional role the cerebellum may play in speech processing, we created a general category for terms associated with task demands (‘expertise’) and mechanisms (‘prediction’).

### No cerebellum meta-analysis

To test the prediction hypothesis, we did a second round of article selection (Figure 1D). Specifically, we repeated the article selection steps outlined above for cerebellum articles but in the n = 6,165 articles in the Neurosynth database that do not report activation in the cerebellar mask. The intersection of these articles and the 1,002,940 articles from our original Pubmed search resulted in a sample of 1547 articles about speech, language and/or articulation. We went through the abstract and methods of these articles by hand to find those involving natural speech perception (i.e., speech/sound/music perception in the complete absence of movement). Studies that explicitly stated that they did not scan the cerebellum were eliminated. This search resulted in 92 (5.94% or 0.64% of the whole Neurosynth database; N = 2026 participants) natural speech/sound perception studies that did not report cerebellum activation (64% speech, 18.5% tones, 11% sounds and 6.5% instrumental music). See Tables S1 and S3 for a complete list of speech perception articles with and without reported cerebellar activity used in this analysis. We did a meta-analysis of speech without a cerebellar activity, using an FDR corrected threshold of q < .01 across the whole brain and a minimum cluster size of 10 voxels. We examined how this meta-analysis differed from speech perception when the cerebellum was active by overlapping the speech perception with and without cerebellum activity meta-analyses. We also examined how these compared to speech production (regardless of whether the cerebellum was active or not) across the whole brain.

## Results

The aim of the study was to test the hypothesis that the cerebellum has a domain-general organisation whose primary function is related to prediction, with cortico-cerebellar connectivity determining what this computation is applied during speech perception. All articles in the Neurosynth database reporting cerebellar activity were identified (n = 8,206) and from this sample studies involving natural (i.e., completely motor free ‘passive’) speech perception (n = 72) and speech production (n = 175) were found and used in cerebellar meta-analysis (Figure 1A).

### Cerebellar meta-analyses

Figure 2A shows significant cerebellar activation during natural speech perception (red) and speech production (blue). Regions activated by both perception and production are in yellow. Speech activity is distributed through much of the cerebellum in a manner that does not correspond to cerebellar lobules. Activity patterns associated with speech perception and production were nearby to each other and showed abrupt transitions. This pattern was not an artifact of statistically contrasting speech perception and production. Figure 2B demonstrates that this arrangement, i.e., nearby perception and production regions, with small regions of overlap, remains when using less conservative corrections for multiple comparison (see Table S4 for the volumes and MNI centre of mass coordinates for these cerebellar meta-analyses).

**Figure 2.**
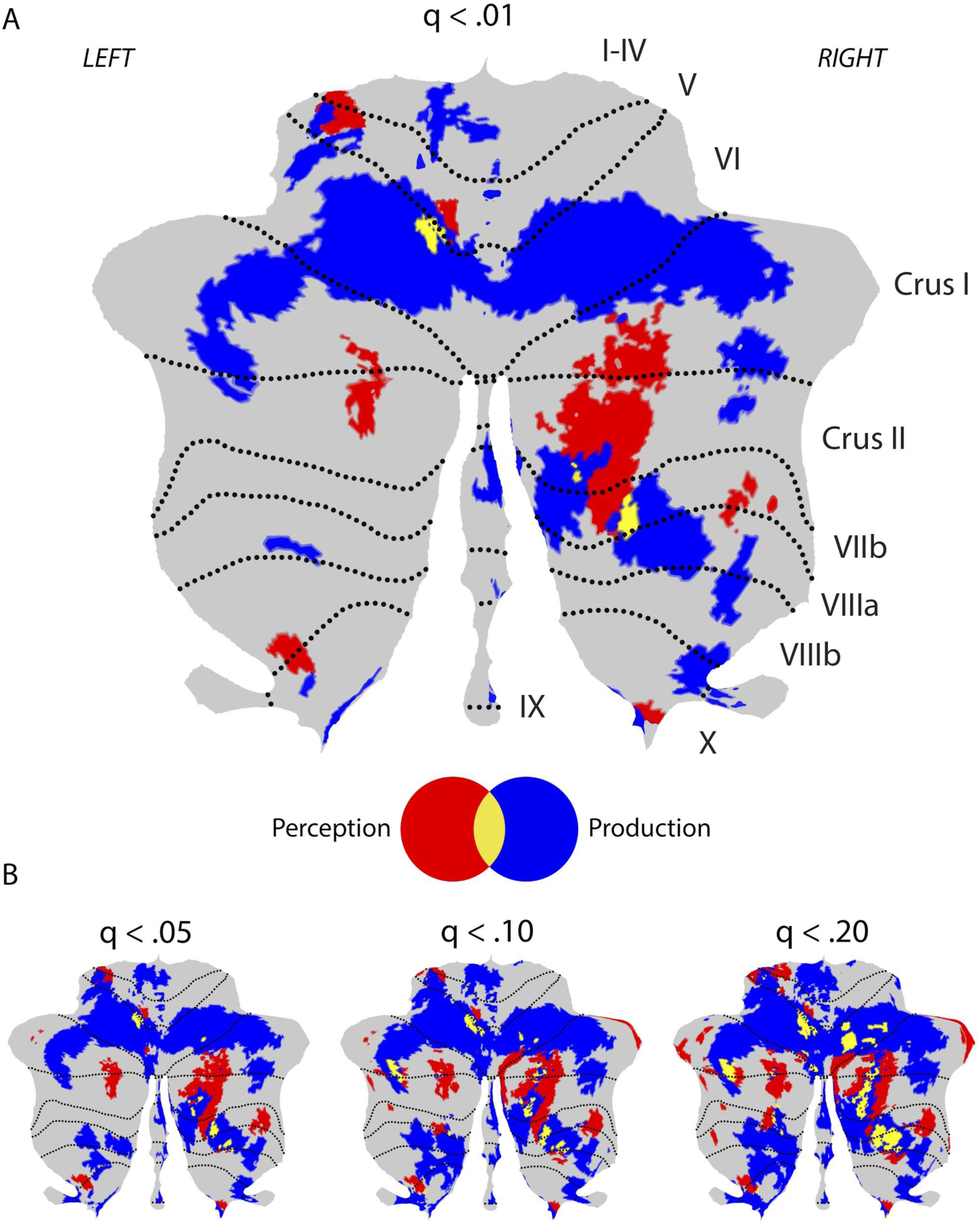
A) Flat map of the cerebellum with regions of cerebellar activity associated with natural speech perception (red), speech production (blue) and both tasks (yellow). Activity patterns are significant at q < .01 false discovery rate (FDR) corrected for multiple comparisons. B) Flat maps at less conservative FDR corrected thresholds. All results are presented with a minimum cluster size of 10 voxels.

To determine if our results were skewed by the inclusion of studies that used sound stimuli not (or only partially) producible by the human vocal tract, we reran the cerebellar meta-analyses excluding these studies. Specifically, we included only studies with speech or tones and excluding those with non-vocal music or sound stimuli. This resulted in 54 studies (75% of the original studies). The resulting spatial correlation between the image in Figure 2A and the new results was r = .99. One difference was that at our FDR corrected threshold of .01 and a cluster size of 10 voxels, only the perception and production overlap in VIIb survives correction. However, at an FDR corrected threshold of .05 and a cluster size of 10 voxels the overlapping VI voxels also survive. Similarly, when studies with tones are removed leaving 46 studies (64% of the original studies), the spatial correlation is r = .98 and the overlapping regions survive but only at a reduced corrected threshold.

### Cerebellar coactivation meta-analysis

We identified brain regions that significantly coactivate with perception, production and regions of perception-production overlap in the cerebellum across thousands of studies. Cortically, perceptual cerebellar regions had greater coactivation, predominantly with the bilateral middle and anterior temporal cortex, angular gyrus and inferior frontal gyrus (Figure 3, red, ‘Perception Network’). Perceptual coactivation also included the caudate bilaterally and the left thalamus. Production regions coactivate more with the precentral and postcentral sulcus and gyrus, the insula, as well as superior parietal regions (Figure 3, blue, ‘Production Network’). Medially, production coactivation regions also included the superior frontal gyrus and, subcortically, the putamen and thalamus bilaterally. Finally, regions in the cerebellum activated by both perception and production coactivate with the central sulcus, inferior parietal cortex, the transverse temporal gyrus and sulcus and nearby superior temporal regions (Figure 3, yellow, ‘Overlap Network’). Medially, overlap coactivation regions also included the superior frontal gyrus and, subcortically, the caudate on the left and putamen and thalamus bilaterally. These subcortical regions were in different locations than clusters in the same structures associated with perceptual and production coactivation. See Table S4 for the volumes and MNI centre of mass coordinates for regions in this coactivation meta-analysis.

**Figure 3.**
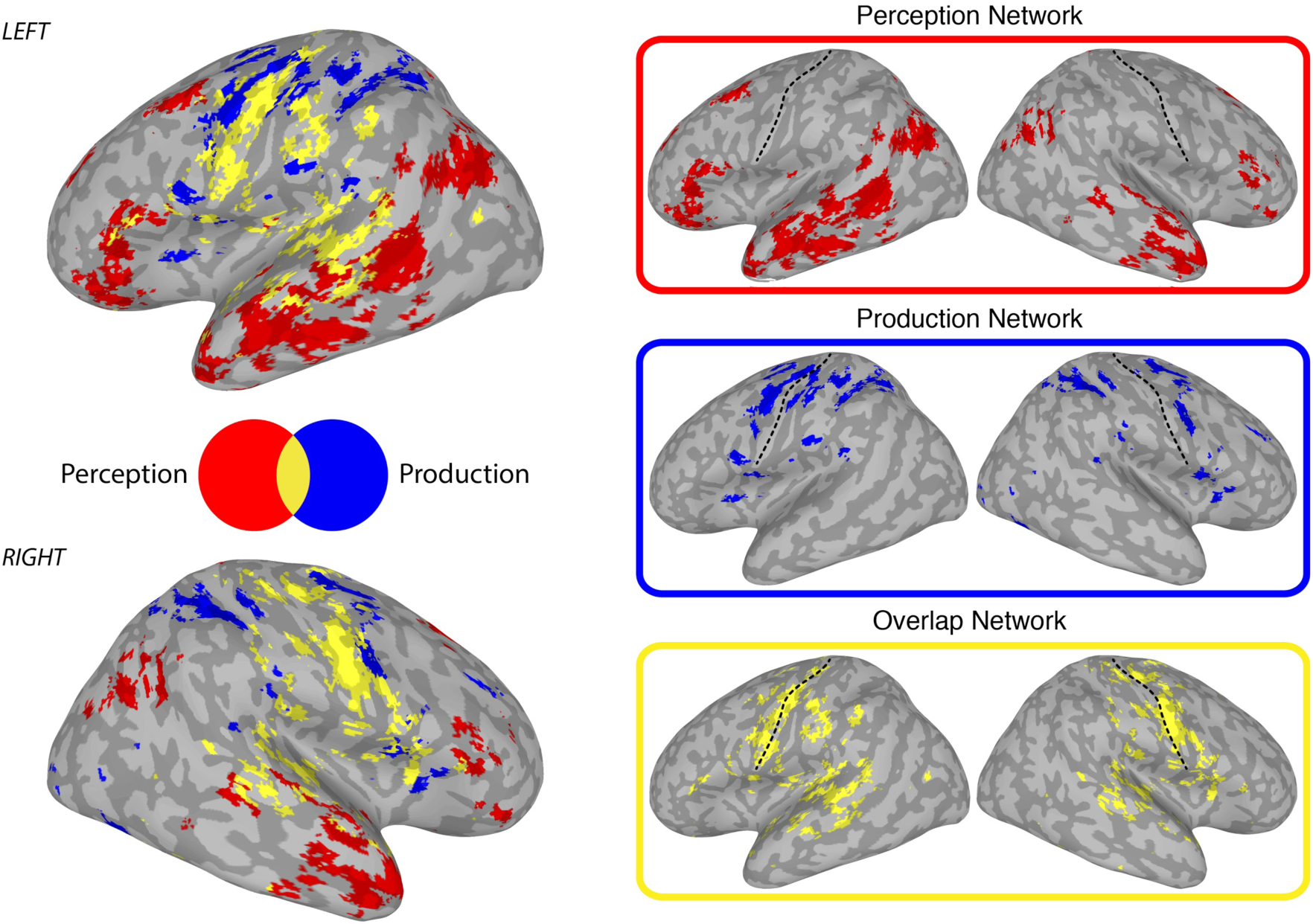
Coactivation networks. Regions in red depict a ‘Perception Network’ that was coactive with regions in the cerebellum associated with natural/passive speech perception. Regions in blue depict a ‘Production Network’ that was coactive with regions in the cerebellum associated with speech production. Regions in yellow depict an ‘Overlap Network’ that was coactive with cerebellar regions associated with both speech perception and speech production. Activity patterns are significant at q < .01 false discovery rate (FDR) corrected for multiple comparisons and presented with a minimum cluster size of 10 voxels. The images on the left are a composite of the three networks on the right. The dashed line divides the central sulcus in half.

The perception, production and overlap networks are similar to prior meta-analyses associated with language and semantic processing, motor planning/sequencing and sensorimotor control, respectively. However, they were also similar to meta-analyses associated with tasks that seemingly have little or nothing to do with speech. For instance, the ‘Perception Network’ was correlated with ‘Theory of Mind’ meta-analyses and the ‘Overlap Network’ with finger tapping (Table 1). This suggests that, although cerebellar regions were identified using studies involving only sounds and speech (without any motor response), the cerebellar networks originating from those regions are not necessarily specific to speech or language.

### Cerebellar functional profile analyses

Functional profiles of perception, production and overlap regions in the cerebellum were probed to assess domain-specificity or generality. We found all terms in the Neurosynth database that predicted activity in each set of regions at an uncorrected statistical threshold of p < .05. A ‘confirmatory’ qualitative analysis revealed that all three sets of perception, production and overlap regions were associated with many cerebellar anatomical and speech related terms. Speech perception regions were uniquely associated with ‘listened’, ‘listening’, ‘passive’ and ‘passively’. They were also associated with speech and language terms and corresponding anatomical regions (e.g., ‘comprehension’, ‘semantic’, ‘temporal’). The speech production and overlap regions (as compared to the speech perception regions) were uniquely associated with ‘active’, ‘overt’ and ‘speech production’. Speech production regions were associated with production terms and corresponding motor regions (e.g., ‘articulatory’, ‘motor cortex’). Activity in overlap regions was associated with terms that tended to be more sensorimotor in nature (e.g., ‘sensorimotor cortex’), but were not as high-level as terms that predicted activity in speech perception regions (e.g., ‘phonological’). Taken together, these results support the validity of the term-based approach.

We next filtered out all anatomical and fMRI terms, leaving 168 task related terms. Figure 4 shows the top ten task-related terms (ranked by their standardized log odds ratio) that predicted activity in cerebellar regions associated with speech perception, production and the overlap of the two. Despite the qualitative differences noted above that conform to task differences, a wide range of tasks were associated with activity in each set of regions and there were also similarities in terms between the sets (e.g., ‘motor’ was the top term associated with each). To quantify this, we calculated the percentage of speech and non-speech terms and the percentage of terms in the perceptual, cognitive, social/emotional and motor domains associated with the speech perception, production and overlap regions (Table 2). Only 23.81% of terms that predicted activity in the three sets of regions were speech specific, even though the regions were identified using a large number of studies that involved only speech and sounds. Speech related terms were roughly equally distributed across perception (5.36%), production (10.71%) and overlap regions (7.74%; χ² = 3.05, p > .05).

**Figure 4.**
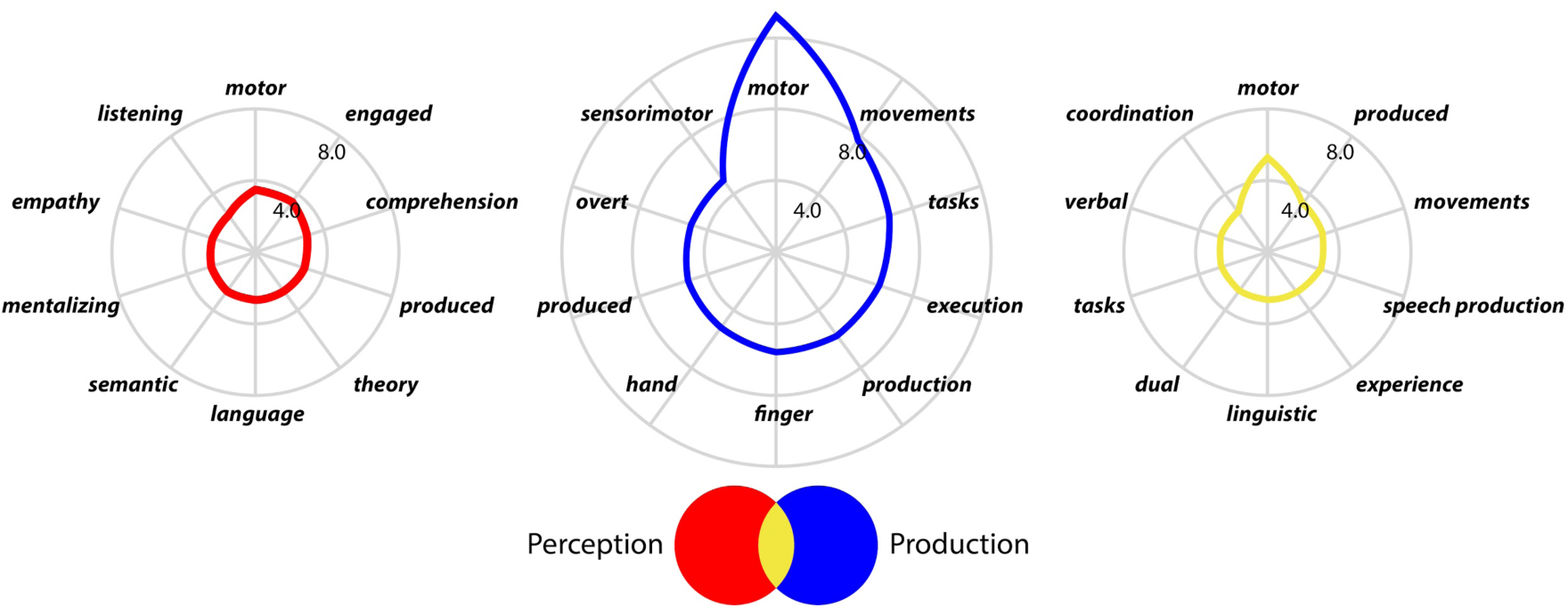
Top ten task related terms from articles that predicted activity in cerebellar regions associated with natural speech perception (red), speech production (blue) and regions activated by both tasks (yellow). The axes display the log odd ratio (in z-scores) for each term.

**Table 2.** All 168 task related terms associated with each set of regions in the cerebellum were categorised as being speech related or not or into four gross psychological domains, counted and presented as percentages showing that the three sets of cerebellar regions are not speech or domain specific.

|  |  | Domains |  |  |  |  |
| --- | --- | --- | --- | --- | --- | --- |
| Regions | Speech | Perceptual | Cognitive | Emotional | Motor | Unspecified |
| Perception | 5.36% | 2.98% | 4.17% | 5.95% | 2.98% | 3.57% |
| Production | 10.71% | 8.33% | 8.33% | 0.00% | 27.38% | 16.67% |
| Overlap | 7.74% | 1.19% | 5.95% | 0.00% | 8.33% | 4.17% |
| Sum | 23.81% | 12.50% | 18.45% | 5.95% | 38.69% | 24.40% |

Including all terms (speech and not speech), we examined whether the terms predictive of cerebellar activity in speech perception, production and overlap regions were equally distributed across our four psychological domains: perceptual, cognitive, social/emotional and motor. The distribution of terms was uniform in the case of the speech perception regions (χ² = 2.48, p > .05), but not uniform in the case of the speech production (χ² = 61.57, p < .001) and overlap regions (χ² = 20.15, p < .01). In explanation, more than half of the terms associated with the speech production regions were from the motor category (though *not* speech/articulatory specific) and no terms from the social/emotional category were associated with either the speech production or overlap regions. About a quarter of all terms (24.40%) did not fall into a psychological domain and were classified as nonspecific. These terms were associated with each set of regions, but the distribution was not uniform (χ² = 22.56, p < .001); four times as many of these terms were associated with production regions in the cerebellum. Taken together, the results suggest that speech perception, production and overlap regions in the cerebellum are associated with a range of perceptual, cognitive and motor tasks well beyond the domains of speech and language.

Finally, about 31.55% of the terms could be labelled as either associated with demands or mechanisms. About 71.70% of these came from terms that could not be labelled with the perceptual, cognitive, social/emotional or motor categories. Demand related terms were associated with increasing task difficulty and/or expertise (e.g., ‘faster’). Mechanism related terms included ‘predictions’ (speech perception and production regions), ‘sequence’ and ‘sequential’ (speech production related regions) and ‘coordination’ and ‘timing’ (speech production and overlap regions). These task-independent terms are consistent with prior accounts that link cerebellar functioning to predictive processing. See Table S5 for all terms and categorisations.

### No cerebellum meta-analysis

Functional profiles demonstrate that regions in the cerebellum associated with speech perception, speech production and their overlap are also associated with a wide range of tasks well outside of the domain of speech, language and vocal motor control. This result supports a domain-general view of cerebellar processing. A domain-general process often attributed to the cerebellum is prediction or expectancy/timing signals that are an aspect of prediction. A hallmark of predictive processing is metabolic savings (i.e., decreases in activity when events are predicted). To test whether there is a decrease in activity when the cerebellum is active, we identified studies in the Neurosynth database involving natural speech perception that did not report activity in the cerebellum (n = 92; Figure 1D). These studies were used in a second meta-analysis that compared whole-brain speech perception related activity when the cerebellum is active versus not active.

As shown in Figure 5, during speech perception, there are striking differences in brain activity as a function of whether the cerebellum is active or not. Specifically, when the cerebellum is active (in red and yellow), cortical activity related to speech perception is primarily located in the superior temporal plane, posterior inferior frontal gyrus and posterior aspect of the superior frontal gyrus. When the cerebellum is not active during speech perception (blue and yellow), there is 1.68 times more brain activity overall that is distributed over a much larger area of the brain. This activity encompasses the same regions as when the cerebellum is active and, additionally, more posterior aspects of the superior and middle temporal gyrus and sulcus, inferior parietal lobule, postcentral gyrus and sulcus, precentral gyrus and sulcus, the inferior frontal gyrus and thalamus. These additional regions are partially captured by a meta-analysis of speech production (shown in white outline). This result is consistent with the idea that the cerebellum plays a role in prediction during passive speech perception.

**Figure 5.**
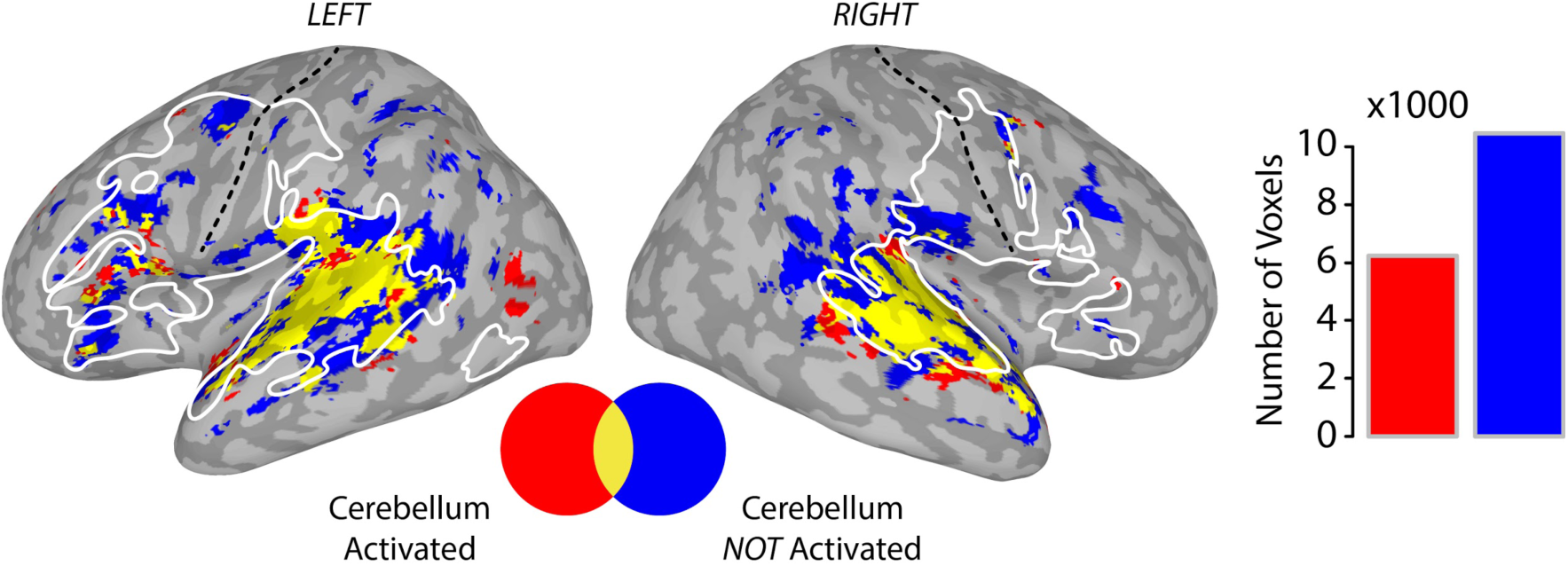
Extra-cerebellar activity patterns associated with natural speech perception when the cerebellum is active (red), not active (blue) and their overlap (yellow) displayed on the lateral surface of the left and right hemispheres. The white outline shows regions active during speech production. Activity patterns are significant at q < .01 FDR corrected and presented with a minimum cluster size of 10 voxels. The bar plot depicts the number of significantly activated voxels during passive speech perception when the cerebellum is active (red) or not (blue).

## Discussion

We tested the hypothesis that the cerebellum implements a domain-general, predictive mechanism that is deployed during speech perception as a function of connectivity. We identified studies from a large neuroimaging database reporting cerebellar activity during naturalistic (i.e., ‘passive’) speech perception without any motor response and compared these to speech production studies. We also found studies involving natural speech perception that did not report cerebellar activity. We used these in neuroimaging meta-analyses and coactivation meta-analyses and term-based cerebellar functional profile analyses (Figure 1; Tables S1-S3). We observed multiple regions of activity throughout the cerebellum related to both speech perception and production that were largely separate, but with some overlap (Figure 2; Table S4). These regions had unique patterns of functional connectivity (Figure 3, red, blue and yellow; Table S4). Across thousands of studies, the functional profiles of these networks (Table 1) and their seed cerebellar regions (Table 2) were not speech or domain-specific (see also Table S5). Regions of the cerebellum activated by speech perception studies were also associated with mechanistic terms like ‘timing’ and ‘prediction’. Finally, when the cerebellum was inactive, there was more cortical activity than when it was active (Figure 5). Here, we review these results in relation to the literature on cerebellar topology and use this to discuss how the results support a domain-general, predictive account of the cerebellum in speech perception.

### Topology

#### Modularity

Consistent with other studies using a range of tasks (Buckner et al., 2011; Diedrichsen et al., 2019; Guell, Gabrieli, et al., 2018; Guell, Schmahmann, et al., 2018; King et al., 2019; Stoodley et al., 2012), the pattern of activity observed for speech traversed lobular boundaries (Figure 2). Though activity patterns do not conform to any obvious anatomy, they do seem to be functionally modular, with sharp boundaries between speech perception and production regions. This is consistent with studies suggesting different functions map to distinct cerebellar regions with abrupt transitions between them (Guell, Gabrieli, et al., 2018; Guell, Schmahmann, et al., 2018; Imamizu et al., 2003; King et al., 2019; Marek et al., 2018). The sharp boundaries in our results are even more striking given that activity patterns were *not* the result of a direct contrast between speech perception and production.

From a functional modularity perspective, speech perception and production activity in different regions likely corresponds to the different subtasks that these functions can be decomposed into. Overlapping regions are likely associated with subtasks similar to both. Indeed, after task decomposition, subtasks engage different cerebellar regions. For example, working-memory can be broken down into an ‘articulatory loop’ and ‘phonological store’ and these subtasks consistently activate different cerebellar regions (Chen & Desmond, 2005a, 2005b; E et al., 2014). Conversely, language and working memory (Ashida et al., 2019; Stoodley et al., 2012) and social mentalizing (Van Overwalle et al., 2014) may overlap in the cerebellum because they share common subtasks.

#### Zones

Consistent with the appearance of functional modularity, it is claimed that there is a higher level of cerebellar organisation into a nonmotor (cognitive, emotional and/or social) zone and two motor zones, corresponding to different connectivity patterns. Specifically, past work has observed a nonmotor zone in posterior lateral regions, especially crus I and II, and two motor zones, one in the anterior lobe and the other around lobule VIII (Buckner et al., 2011; Gellersen et al., 2017; Stoodley & Schmahmann, 2018). These have even been further subdivided into triple nonmotor and double motor zones (Buckner et al., 2011; Guell, Gabrieli, et al., 2018; Guell, Schmahmann, et al., 2018).

If one focuses only on the largest regions of activity in our data, there seems to be a medial speech perception zone (Crus I/IIl; HVIIb; Figure 2, red) and two speech production zones, one in the anterior and one in the posterior lobe (dorsal and ventral to the medial perception zone in Figure 2, blue). Speech perception also overlaps with speech production in two somatotopically organised motor zones at the same location of lip and tongue representations (e.g., compare Figure 2 to Figure 7B in Boillat et al. 2020) (Boillat et al., 2020; Buckner et al., 2011; Grodd et al., 2001; Guell, Gabrieli, et al., 2018).

However, nonmotor vs two motor zones oversimplifies the observed pattern of activity (Diedrichsen et al., 2019). Our results suggest that there are up to 10 distinct zones for speech perception and more for speech production both distributed throughout both nonmotor and motor zones. This complexity might again be attributed to the fact that speech is a sensorimotor task that can be decomposed into many subtasks that do not neatly conform to ‘cognitive’ and ‘motor’ categories (Skipper et al., 2017). The latter cerebellar ‘sandwich’ (Hurley, 2001) view (of a cognitive between motor zones) may derive from attempting to map multiple gross functions onto the cerebellum using winner-takes-all-like strategies. Our data maps a single ‘function’ and shows that large swathes of the cerebellum are involved in speech perception, arguing for finer task decomposition to understand individual regions of activity.

### Domain-generality

Based on the uniformity of cerebellar structure, theories like the Universal Cerebellar Transform (UCT)(Schmahmann, 2019; Schmahmann et al., 2019), claim that cerebellar functions are domain-general, performing a similar computation throughout. Functional specialization in these models is determined by variations in cerebellar location and corresponding cortico-cerebellar connectivity. The fact that our data appears functionally modular while not conforming to any obvious anatomical boundaries could be consistent with either domain-specific or general accounts.

However, domain-generality was supported by the functional profile analyses. Specifically, speech perception regions in the cerebellum were not speech specific. They were also equally associated with perceptual, motor, cognitive and emotional terms generally (see Tables 1 and 2) and (in alphabetical order) attention, audition, finger and hand movements, language, memory, pain, speech, theory of mind and vision terms more specifically (see Table S1). Our speech perception studies were all sound and speech related, with no movements associated with them. It would, therefore, be hard to explain the results with a domain-specific theory as it is unlikely that every one of the nonspeech terms associated with speech perception regions have a linguistic explanation.

It is important to note that although domain generality implies a common computation in the cerebellum, it is entirely possible that this computation is used by different cognitive and motor processes in different ways. For instance, the cerebellum may contribute a timing signal that is used for prediction in some tasks, coordination in other tasks, and learning in a third set of tasks. Just as the prefrontal cortex contributes working memory to a variety of tasks that use this resource in a variety of ways, the cerebellum’s common computation may be utilized in different ways depending on the process it is contributing to.

### Connectivity

If the cerebellum does perform a domain-general role in speech perception, language comprehension and any other domain, that role must be determined by cortico-cerebellar connectivity. As we reviewed, it has been observed that there are abrupt functional divisions in the cerebellum and these have been shown to conform to structural and functional connectivity, perhaps determining the functional specialisation of those regions (Schmahmann, 2019; Schmahmann et al., 2019). Furthermore, this connectivity is said to conform to the division of the cerebellum into (multiple) nonmotor zones and (two) motor zones (Buckner et al., 2011). Indeed, the speech perception, production and overlap sets of regions formed surprisingly distinct networks with other subcortical and cortical regions as determined by coactivation meta-analysis. In functional terms, speech perception regions tended to form networks with “higher-level” language regions, speech production regions formed networks with premotor and other “higher-level” motor/speech production regions and overlap regions formed networks with primary auditory and primary motor regions. That is, overlap regions were distinctly sensorimotor (Skipper et al., 2017; Skipper & Hasson, 2017). However, these connectivity patterns were also domain-general (Table 1), suggesting, again, that cerebellar regions are not speech specific. They may become so in speech-only circumstances, but we could not determine this as we used *all* studies to do the coactivation analysis.

### Prediction

Prediction is an increasingly accepted mechanistic account of how the brain works generally (Clark, 2013; Keller & Mrsic-Flogel, 2018) and the cerebellum more specifically, especially in the domain of motor control (Moberget & Ivry, 2019; Popa & Ebner, 2018). Domain-general theories suggest that the computation the cerebellum contributes to any one domain should be similar to the computation it contributes to others. Although it is possible that different processes may use this consistent contribution in different ways, we also might expect to see commonalities between behaviours in how the cerebellum’s contribution is used. Indeed, it has been argued that the predictive role that the cerebellum plays in motor control is reused in speech perception and higher-level linguistic domains like semantics (Moberget et al., 2014; Schwartze et al., 2012).

Consistent with this account, our functional profiles of speech perception, production and overlap regions were all associated with the terms ‘coordination’, ‘timing’ and ‘prediction’. These regions were each associated with unique cerebellar locations and associated connectivity profiles, consistent with a domain-general account. This suggests the possibility that these regions and associated networks are predicting at different levels of analysis, perhaps corresponding to the superior/inferior motor zone and medial cognitive zones. Indeed, sensory prediction related processes that mediate sensorimotor adaptation have been demonstrated in the superior motor zone where we show speech perception/production overlap (Guediche et al., 2015). Furthermore, the distribution of activity in the medial portion of the cerebellum completely overlaps the peaks of activity in five fMRI studies of linguistic predictability (Bonhage et al., 2015; D’Mello et al., 2017; Lesage et al., 2017; Moberget et al., 2014; Moberget & Ivry, 2016; Tourville et al., 2008). Consistent with this, overlap networks were more associated with sensorimotor regions whereas the perception-only networks were more associated with regions mediating higher-level linguistic processes in prior studies. Indeed, perception-only regions were uniquely associated with the term ‘comprehension’.

We also generated more direct evidence for the predictive account. That is, we tested a key tenant of predictive models, that they result in less activity when predictions are accurate (Moberget et al., 2014; Skipper, 2014). If the cerebellum plays a predictive role, we hypothesized that cortical activity should be reduced when the cerebellum was active in contrast to when it is not. Indeed, we found that when the cerebellum was active there was a nearly two-fold reduction of cortical activity during speech perception compared to when the cerebellum was not active. Furthermore, much of this reduction was in the aforementioned sensorimotor and higher-level linguistic networks associated with the speech perception/production overlap and speech perception only networks, respectively. This suggests the possibility that cortico-cerebellar and cortico-cortical predictions trade off with each other. A hallmark of cerebellar function is expertise whereas a hallmark of the neocortex is flexibility. Perhaps the cerebellum is involved in predictions associated with perceiving more well learned speech whereas the cortex more flexibly applies predictions in new contexts. Consistent with this, the cerebellum seems to play a specific role in ‘automatic speech’, i.e., overlearned material (Ackermann et al., 1998; Ackermann, 2008).

### Limitations

First, our sample of natural speech perception articles includes (as a minority) studies in which participants passively listened to instrumental music, sounds and tones. We made the decision to include these studies because it is unclear when exactly sound perception changes to speech perception, many languages are tonal (Yip, 2002), and there is neuroimaging and behavioural evidence that speech and music perception draw on a common neural resource (Kunert & Slevc, 2015; Peretz et al., 2015). To examine the impact of these studies on the observed patterns of cerebellar activation, we re-ran the baseline meta-analyses with just speech and tone studies and then just speech studies alone. Patterns of activation were nearly identical to the case where all studies (speech and nonspeech) were included. However, with just speech studies regions of production-perception overlap were mostly only observed at a reduced (though still corrected) statistical threshold. This may reflect a lack of statistical power due to the reduced sample. It is also possible that the overlapping area is enhanced by nonspeech studies because of motor recruitment. There is evidence that cortical motor systems become more engaged in speech perception as auditory signals become more foreign (Wilson, 2009; Wilson & Iacoboni, 2006). The extent to which regions of perception-production overlap in the cerebellum are observed during the perception of clear, native speech, needs to be further explored.

Second, there is a possibility that our sample of passive speech perception studies in which cerebellar activation was not reported (n = 92) includes studies that actually did find—but failed to report— cerebellar activations. If such studies are in this sample, they would likely reduce differences between studies reporting cerebellar activation and studies not reporting it. Removing these studies (if they exist in our sample) would likely lead to greater observed differences in cortical activity between the groups shown in Figure 5.

Finally, our findings reflect the published literature. We cannot control for the quality of the included data, e.g., whether appropriate high-level contrasts were used. Furthermore, results may reflect theoretical biases. For instance, predictive coding is a trending topic and there is a known predictive role of the cerebellum in motor control. Thus, there may be a bias to discuss cerebellar activity in the context of a predictive framework. However, we included a large number of studies, almost none of which had any specific interest in the cerebellum (e.g., none of the speech perception articles had ‘cerebell’* in the title). They simply happened to report cerebellar activity during a task that met our criteria, likely reducing the impact of bias.

### Conclusions

What role does the cerebellum play in speech perception? Our results are consistent with the perspective that the cerebellum plays a domain-general and predictive role in all functions, including speech. Furthermore, the type of prediction (e.g., motor or semantic) must be determined by task (and subtask) specific cortico-cerebellar connectivity.

## Acknowledgments

This work was supported by grants from the British Academy, Corpus Christi College Oxford, Acadia University and the Natural Sciences and Engineering Research Council (NSERC) of Canada to DRL. It was also partially supported by EPSRC EP/M026965/1 to JIS. JIS would like to thank a Banana. The authors would also like to thank S. Bobbitt, L. Wyatt and N. Guest for help coding the manuscripts that went into the meta-analysis.

## Supplementary Tables

**Table S1.** Studies with passive speech/sound perception reporting cerebellar activation.

| # | Authors | Title | n | Task |
| --- | --- | --- | --- | --- |
| 1 | Mutschler et al. (2007) | A rapid sound-action association effect in human insular cortex. | 10 | passively listened to piano melodies after either actively or passively learning the melodies |
| 2 | Milivojevic et al. (2016) | Coding of Event Nodes and Narrative Context in the Hippocampus. | 19 | passively watched an audiovisual movie |
| 3 | Nastase et al. (2015) | Connectivity in the human brain dissociates entropy and complexity of auditory inputs. | 20 | passively listened to tones of different levels of entropy and complexity |
| 4 | Noohi et al. (2017) | Functional Brain Activation in Response to a Clinical Vestibular Test Correlates with Balance. | 16 | passively listened to tones and received skull taps |
| 5 | Li et al. (2017) | Intranasal oxytocin, but not vasopressin, augments neural responses to toddlers in human fathers. | 31 | passively viewed pictures of infants and listened to infant cries |
| 6 | Li et al. (2015) | Listening to music in a risk-reward context: The roles of the temporoparietal junction and the orbitofrontal/insular cortices in reward-anticipation, reward-gain, and reward-loss. | 23 | passively listened to music |
| 7 | Oh et al. (2018) | Multisensory vestibular, vestibular-auditory, and auditory network effects revealed by parametric sound pressure stimulation. | 20 | passively listened to tone bursts |
| 8 | Wallmark et al. (2018) | Neurophysiological Effects of Trait Empathy in Music Listening. | 35 | passively listened to different instruments (experiment 1) and songs (experiment 2) |
| 9 | Farahani et al. (2017) | Spatiotemporal reconstruction of auditory steady-state responses to acoustic amplitude modulations: Potential sources beyond the auditory pathway. | 19 | passively listened to modulated white noise while watching a silent movie |
| 10 | Li et al. (2017) | The spatiotemporal pattern of pure tone processing: A single-trial EEG-fMRI study. | 33 | passively listened to tones |
| 11 | Lopopolo et al. (2017) | Using stochastic language models (SLM) to map lexical, syntactic, and phonological information processing in the brain. | 24 | passively listened to stories |
| 12 | Downar et al. (2002) | A cortical network sensitive to stimulus salience in a neutral behavioral context across multiple sensory modalities. | 10 | passively experienced visual, auditory, and tactile stimulation |
| 13 | Ogawa et al. (2013) | Audio-visual perception of 3D cinematography: an fMRI study using condition-based and computation-based analyses. | 16 | passively watched audiovisual movie with alternating 3D and 2D sound and vision |
| 14 | Foxe et al. (2002) | Auditory-somatosensory multisensory processing in auditory association cortex: an fMRI study. | 12 | passively listened to scratching sounds and experienced somatosensory stimulation on fingers |
| 15 | Wilson et al. (2008) | Beyond superior temporal cortex: intersubject correlations in narrative speech comprehension. | 24 | passively watched an audiovisual movie of an actor describing a story |
| 16 | Klein et al. (2006) | Bilingual brain organization: a functional magnetic resonance adaptation study. | 16 | listened to a list of identical words with the final word either being identical or different and either in the same or different language |
| 17 | Hesling et al. (2005) | Cerebral mechanisms of prosodic sensory integration using low-frequency bands of connected speech. | 12 | passively listened to either filtered or unfiltered speech |
| 18 | Wiethoff et al. (2008) | Cerebral processing of emotional prosody--influence of acoustic parameters and arousal. | 24 | passively listened to words spoken with different emotional prosody |
| 19 | Jeong et al. (2011) | Congruence of happy and sad emotion in music and faces modifies cortical audiovisual activation. | 15 | passively listened to happy or sad music while viewing happy or sad faces |
| 20 | Vander Wyk et al. (2010) | Cortical integration of audio-visual speech and non-speech stimuli. | 19 | passively listened to either tones or speech while viewing either ellipses or circles |
| 21 | McNealy et al. (2006) | Cracking the language code: neural mechanisms underlying speech parsing. | 27 | passively listened to strings of syllables linked together |
| 22 | Hertrich et al. (2011) | Cross-modal interactions during perception of audiovisual speech and nonspeech signals: an fMRI study. | 20 | passively listened to speech or nonspeech (tones) |
| 23 | Engel et al. (2009) | Different categories of living and non-living sound-sources activate distinct cortical networks. | 20 | passively listened to sounds of different categories and silently determined which category each one belonged to |
| 24 | Leech et al. (2011) | Distributed processing and cortical specialization for speech and environmental sounds in human temporal cortex. | 7 | passively listened to everyday environmental sounds and speech sounds |
| 25 | Halai et al. (2015) | Dual-echo fMRI can detect activations in inferior temporal lobe during intelligible speech comprehension. | 20 | passively listened to normal speech, rotated speech, and rotated vocoded speech |
| 26 | Chapin et al. (2010) | Dynamic emotional and neural responses to music depend on performance expression and listener experience. | 21 | passively listened to music |
| 27 | Chow et al. (2014) | Embodied comprehension of stories: interactions between language regions and modality-specific neural systems. | 24 | passively listened to stories |
| 28 | Obleser et al. (2010) | Expectancy constraints in degraded speech modulate the language comprehension network. | 16 | passively listened to noise-vocoded sentences |
| 29 | Woods et al. (2014) | Expert athletes activate somatosensory and motor planning regions of the brain when passively listening to familiar sports sounds. | 57 | passively listened to sports sounds and non-sports sounds |
| 30 | Eldar et al. (2007) | Feeling the real world: limbic response to music depends on related content. | 14 | passively watched film clips while listening to music of different emotional valences |
| 31 | Dick et al. (2014) | Frontal and temporal contributions to understanding the iconic co-speech gestures that accompany speech. | 14 | passively watched audiovisual clip of speaker using gestures |
| 32 | Smirnov et al. (2014) | Fronto-parietal network supports context-dependent speech comprehension. | 20 | passively listened to narratives |
| 33 | Specht et al. (2003) | Functional segregation of the temporal lobes into highly differentiated subsystems for auditory perception: an auditory rapid event-related fMRI-task. | 12 | passively listened to tones, sounds of instruments and animals, or words with one or two syllables |
| 34 | Holle et al. (2010) | Integration of iconic gestures and speech in left superior temporal areas boosts speech comprehension under adverse listening conditions. | 16 | passively watched audiovisual clips of an actor speaking while making gestures that matched what was being said |
| 35 | Mueller et al. (2011) | Investigating brain response to music: a comparison of different fMRI acquisition schemes. | 20 | passively listened to manipulated and non-manipulated music clips |
| 36 | Herdener et al. (2014) | Jazz drummers recruit language-specific areas for the processing of rhythmic structure. | 22 | passively listened to a rhythmic sequence |
| 37 | Alluri et al. (2012) | Large-scale brain networks emerge from dynamic processing of musical timbre, key and rhythm. | 11 | passively listened to music |
| 38 | Boldt et al. (2013) | Listening to an audio drama activates two processing networks, one for all sounds, another exclusively for speech. | 13 | passively watched an audiovisual movie |
| 39 | Bengtsson et al. (2009) | Listening to rhythms activates motor and premotor cortices. | 17 | passively listened to rhythms |
| 40 | Skipper et al. (2005) | Listening to talking faces: motor cortical activation during speech perception. | 9 | passively listened to stories, watched audiovisual movies of stories, or watched silent video clips |
| 41 | Lee et al. (2011) | Long-term music training tunes how the brain temporally binds signals from multiple senses. | 37 | passively watched audiovisual movie of a speaker uttering short sentences or a man playing piano |
| 42 | Raposo et al. (2009) | Modulation of motor and premotor cortices by actions, action words and action sentences. | 22 | passively listened to action words and noise bursts |
| 43 | Lambert et al. (2002) | Neural substrates of animal mental imagery: calcarine sulcus and dorsal pathway involvement--an fMRI study. | 6 | passively listened to animal names |
| 44 | Nagels et al. (2013) | Neural substrates of figurative language during natural speech perception: an fMRI study. | 16 | passively listened to stories |
| 45 | Strobel et al. (2008) | Novelty and target processing during an auditory novelty oddball: a simultaneous event-related potential and functional magnetic resonance imaging study. | 14 | passively listened to sine tones and environmental sounds |
| 46 | Holloway et al. (2015) | Orthographic dependency in the neural correlates of reading: evidence from audiovisual integration in English readers. | 18 | passively listened to letter names, number names, and phonemes |
| 47 | Chow et al. (2015) | Personal experience with narrated events modulates functional connectivity within visual and motor systems during story comprehension. | 24 | passively listened to stories |
| 48 | Park et al. (2013) | Personality traits modulate neural responses to emotions expressed in music. | 12 | passively listened to musical excerpts that conveyed happiness, sadness, or fear |
| 49 | Willems et al. (2016) | Prediction During Natural Language Comprehension. | 24 | passively listened to excerpts from novels |
| 50 | Campbell et al. (2007) | Primary and secondary neural networks of auditory prepulse inhibition: a functional magnetic resonance imaging study of sensorimotor gating of the human acoustic startle response. | 16 | passively listened to white noise (startle stimulus) |
| 51 | Lewald et al. (2008) | Processing of sound location in human cortex. | 11 | passively listened to white noise bursts |
| 52 | Giraud et al. (2000) | Representation of the temporal envelope of sounds in the human brain. | 5 | passively listened to different frequencies of sound |
| 53 | Laufer et al. (2008) | Sensory and cognitive mechanisms of change detection in the context of speech. | 20 | passively listened to nonwords and words |
| 54 | Londei et al. (2010) | Sensory-motor brain network connectivity for speech comprehension. | 8 | passively listened to words, pseudowords, or reversed words |
| 55 | Deschamps et al. (2014) | Sequencing at the syllabic and supra-syllabic levels during speech perception: an fMRI study. | 15 | passively listened to syllables |
| 56 | Straube et al. (2010) | Social cues, mentalizing and the neural processing of speech accompanied by gestures. | 18 | passively watched audio visual movie of speaker performing gestures |
| 57 | Nardo et al. (2014) | Spatial orienting in complex audiovisual environments. | 26 | passively watched audio visual movie portraying everyday situations |
| 58 | Yakunina et al. (2013) | Spatiotemporal Segregation of Neural Response to Auditory Stimulation: An fMRI Study Using Independent Component Analysis and Frequency-Domain Analysis. | 19 | passively listened to continuous broadband noise |
| 59 | Rohl et al. (2011) | Spectral loudness summation takes place in the primary auditory cortex. | 22 | passively listened to pink noise of different bandwidths |
| 60 | McGettigan et al. (2012) | Speech comprehension aided by multiple modalities: behavioural and neural interactions. | 12 | passively watched audiovisual movies of a speaker with varied auditory clarity, visual clarity, and linguistic predictability |
| 61 | Paulesu et al. (2009) | Supercalifragilisticexpialidocious: how the brain learns words never heard before. | 6 | passively listened to and learned a list of nonwords |
| 62 | Hugdahl et al. (2003) | The effects of attention on speech perception: an fMRI study. | 13 | passively listened to single vowels, three-phoneme pseudowords and three- and four-phoneme real nouns and words |
| 63 | Lockwood et al. (1999) | The functional anatomy of the normal human auditory system: responses to 0.5 and 4.0 kHz tones at varied intensities. | 12 | passively listened to sine-wave tones presented to right ear |
| 64 | Anders et al. (2008) | The human amygdala is sensitive to the valence of pictures and sounds irrespective of arousal: an fMRI study. | 24 | passively listened to non-linguistic auditory stimuli |
| 65 | Abutalebi et al. (2007) | The neural cost of the auditory perception of language switches: an event-related functional magnetic resonance imaging study in bilinguals. | 12 | passively listened to stories that switched between L1 and L2 |
| 66 | Mayer et al. (2009) | The neural networks underlying auditory sensory gating. | 21 | passively listened to tones |
| 67 | Evans et al. (2014) | The pathways for intelligible speech: multivariate and univariate perspectives. | 12 | passively listened to natural speech, noise-vocoded speech, spectrally rotated speech, and spectrally rotated-noise-vocoded speech |
| 68 | Lyons et al. (2010) | The role of personal experience in the neural processing of action-related language. | 21 | passively listened to sentences |
| 69 | Bekinschtein et al. (2011) | Why clowns taste funny: the relationship between humor and semantic ambiguity. | 12 | passively listened to ambiguous sentences, unambiguous sentences, ambiguous jokes, low-ambiguity jokes, and signal-correlated noise |
| 70 | Osaka et al. (2004) | A word expressing affective pain activates the anterior cingulate cortex in the human brain: an fMRI study. | 20 | passively listened to onomatopoeic words highly suggestive of pain |
| 71 | Lepping et al. (2016) | Neural Processing of Emotional Musical and Non-musical Stimuli in Depression. | 22 | passively listened to music and pure tones |
| 72 | Raz et al. (2017) | Robust inter-subject audiovisual decoding in functional magnetic resonance imaging using high-dimensional regression. | 23 | passively viewed films or passively listened to music |

**Table S2.**
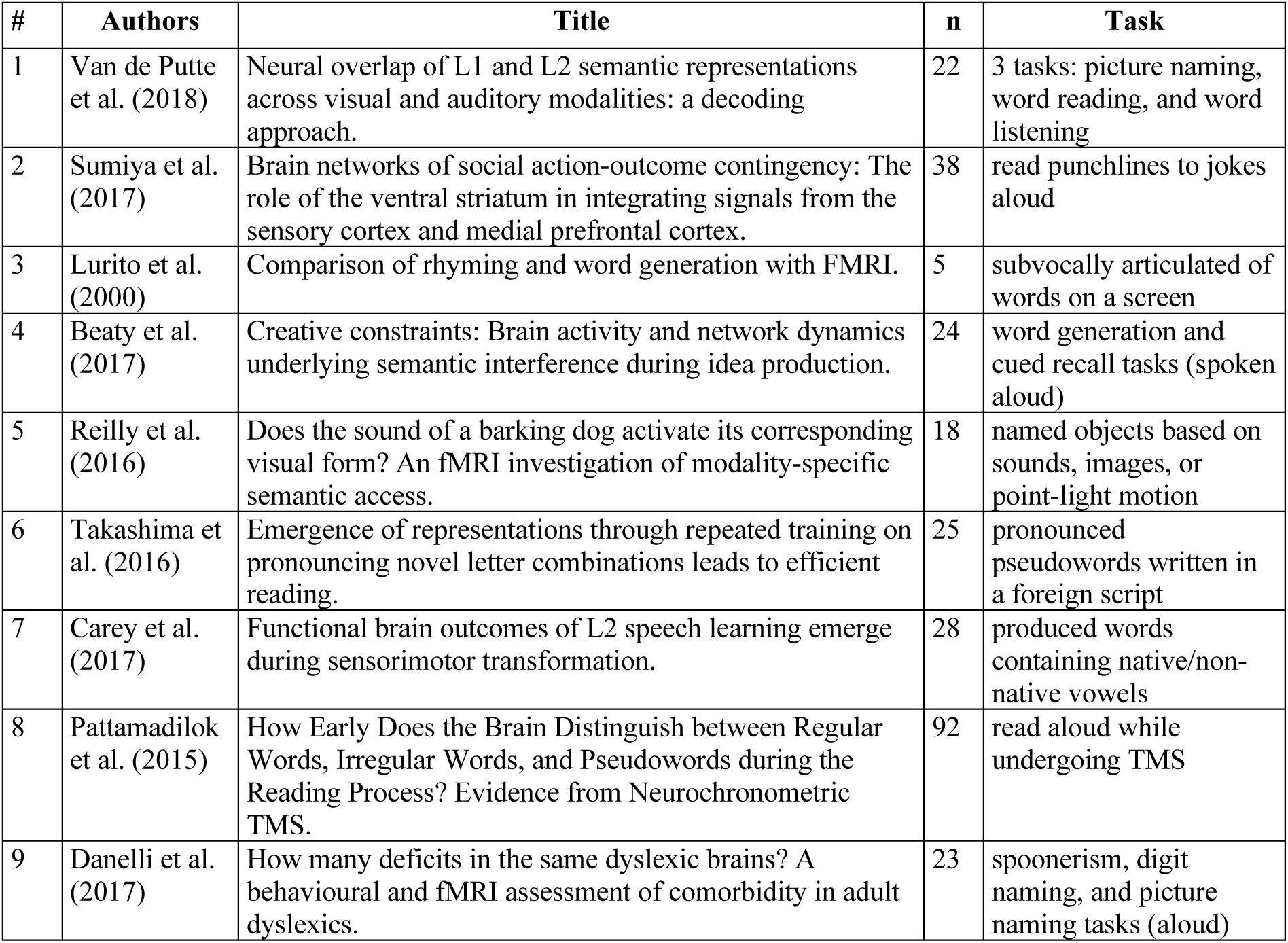
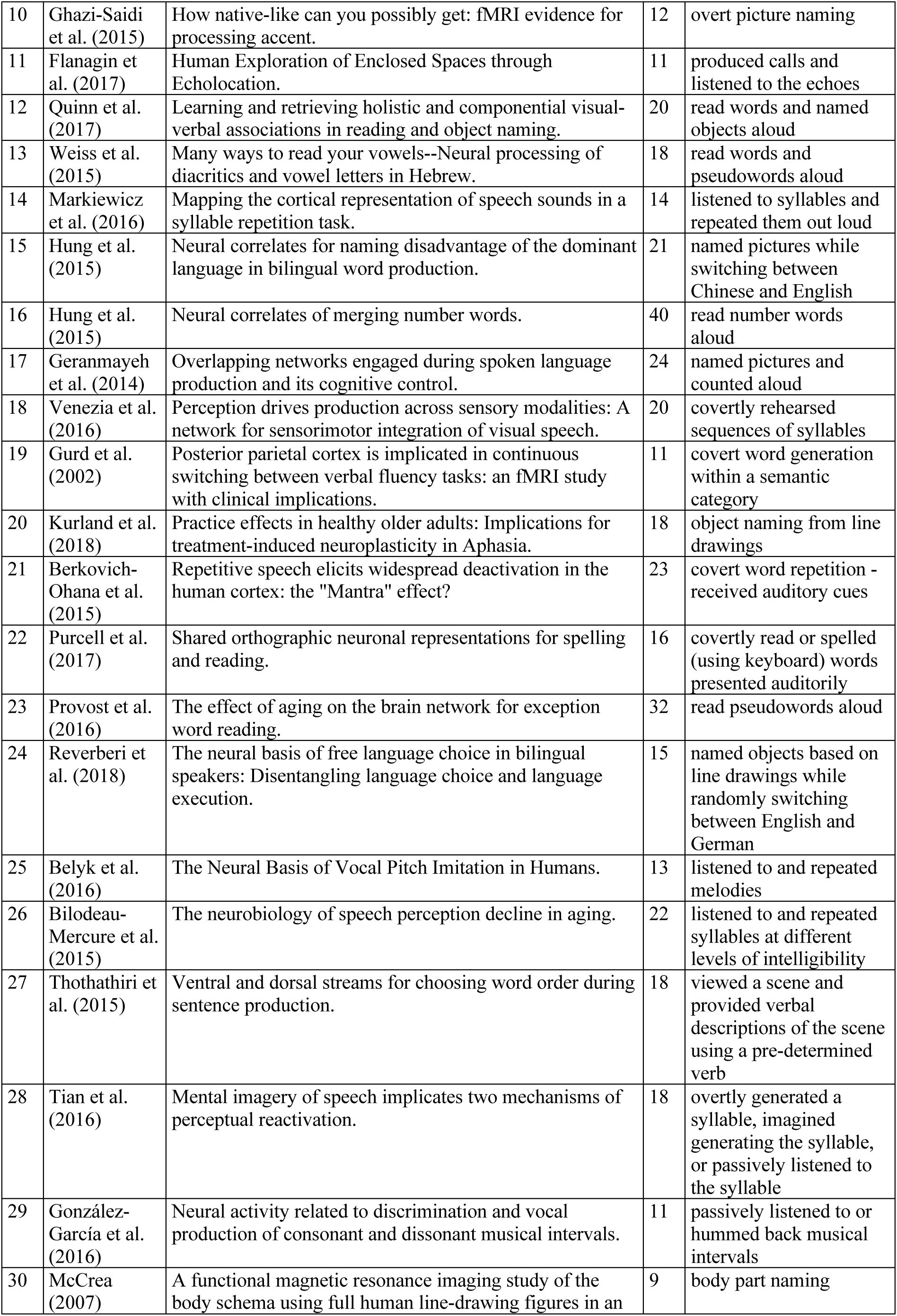
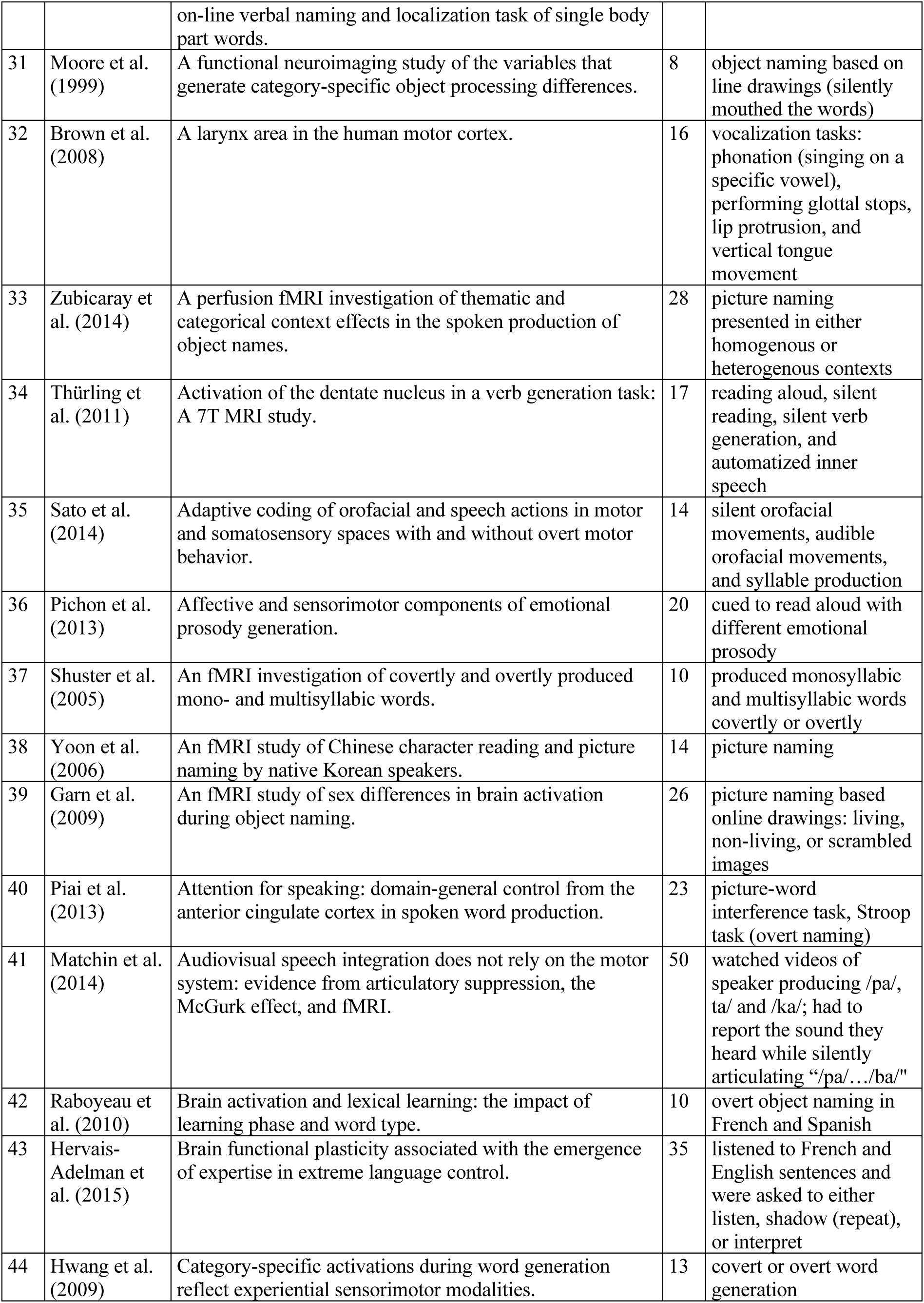
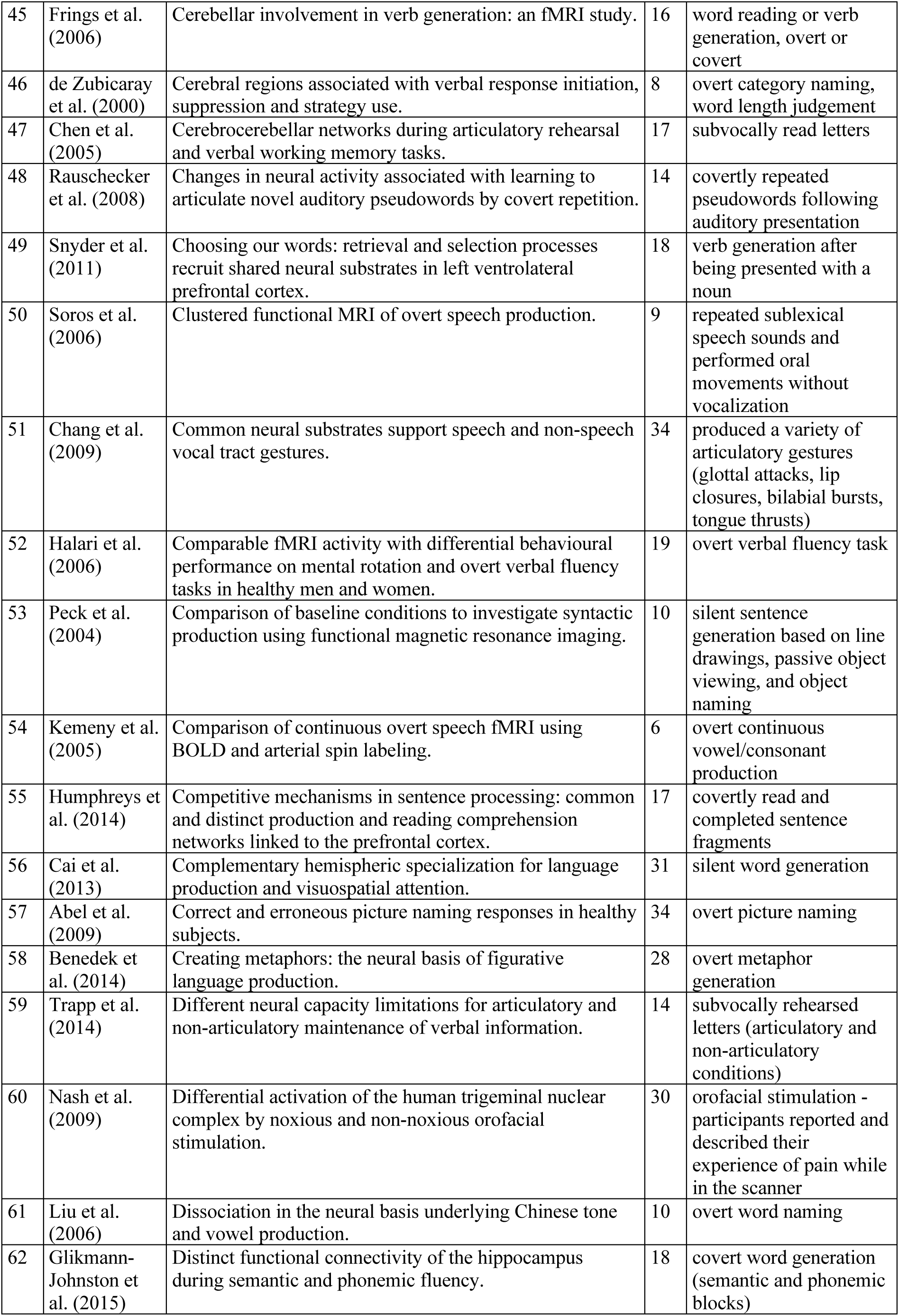
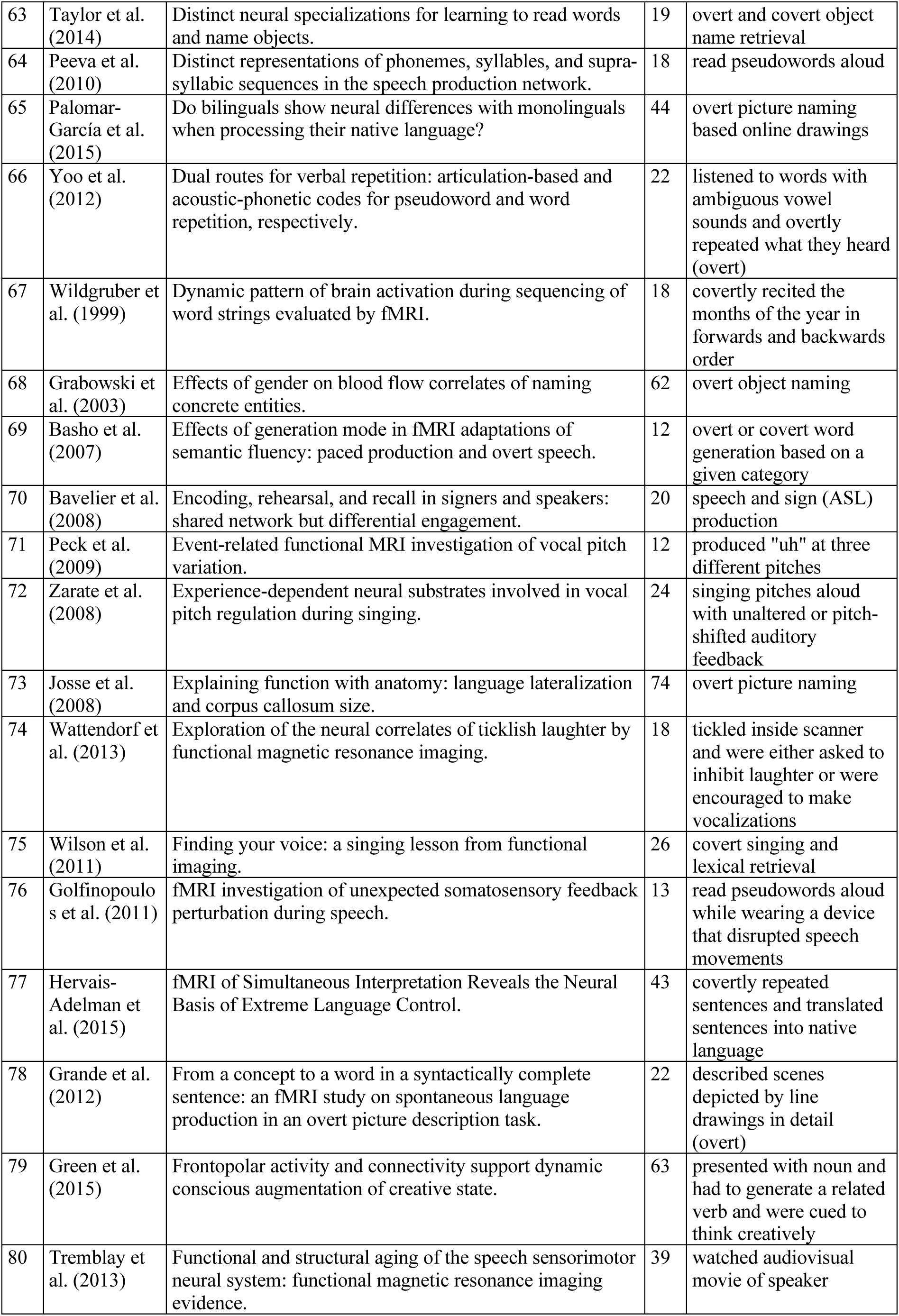
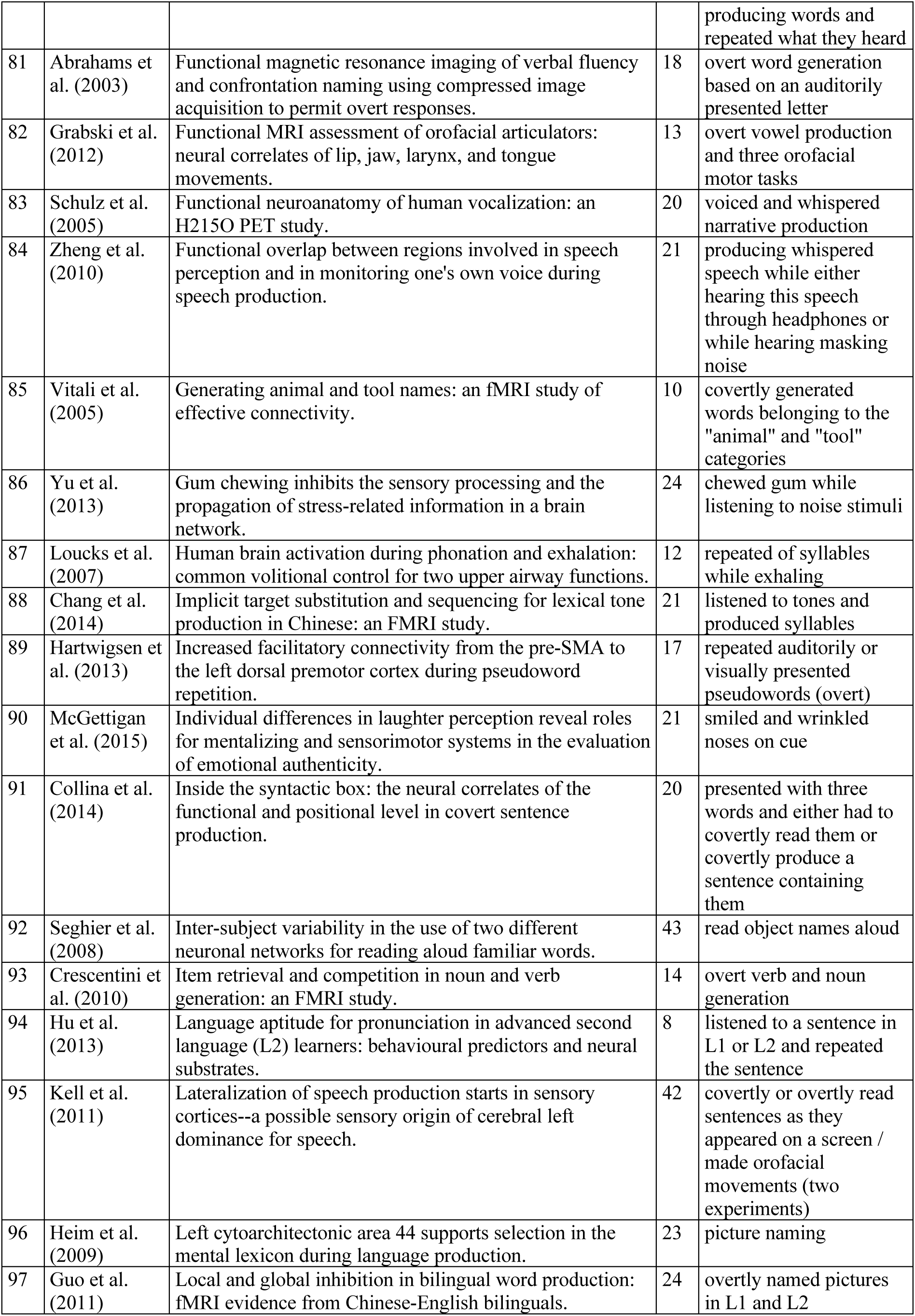
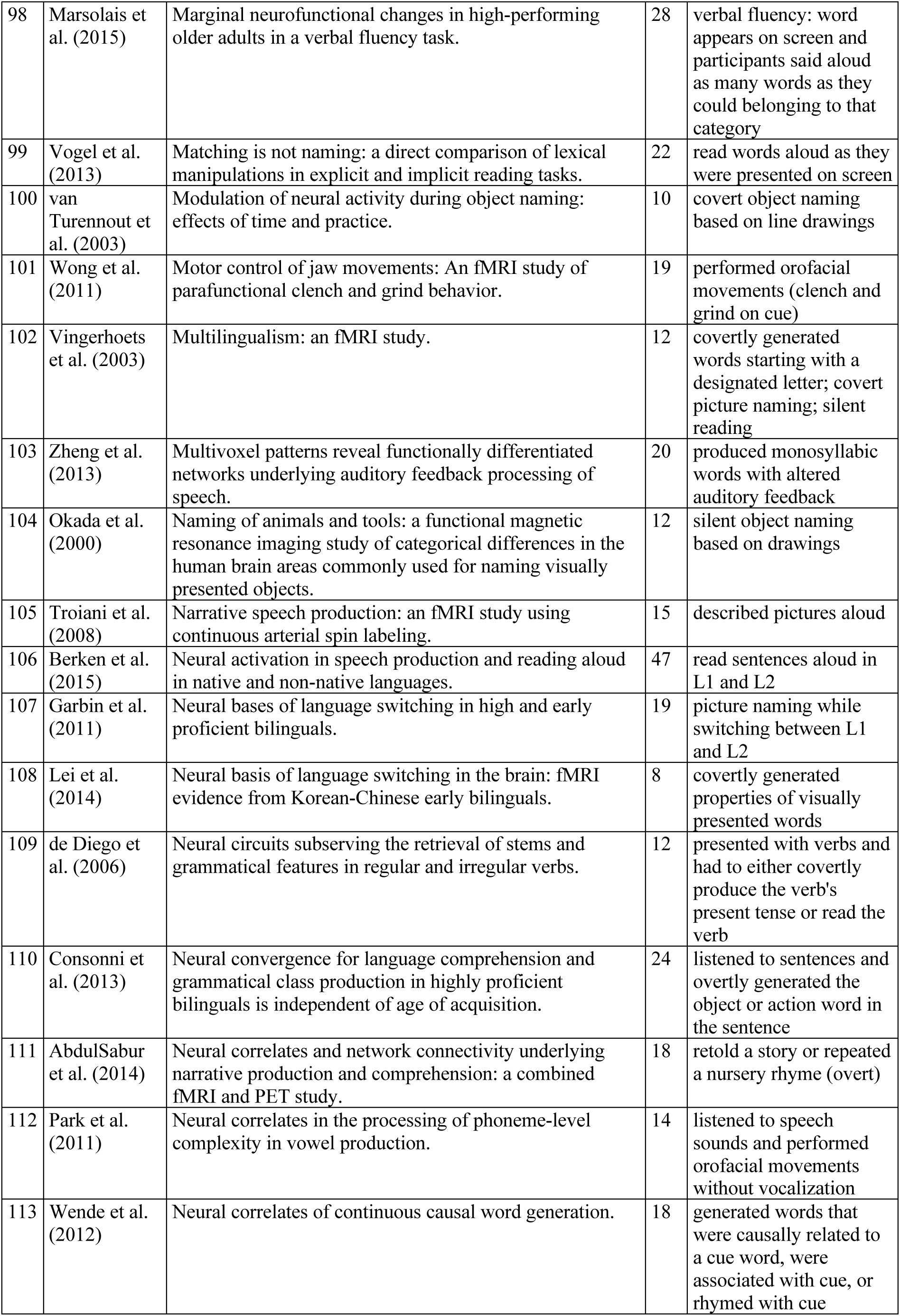
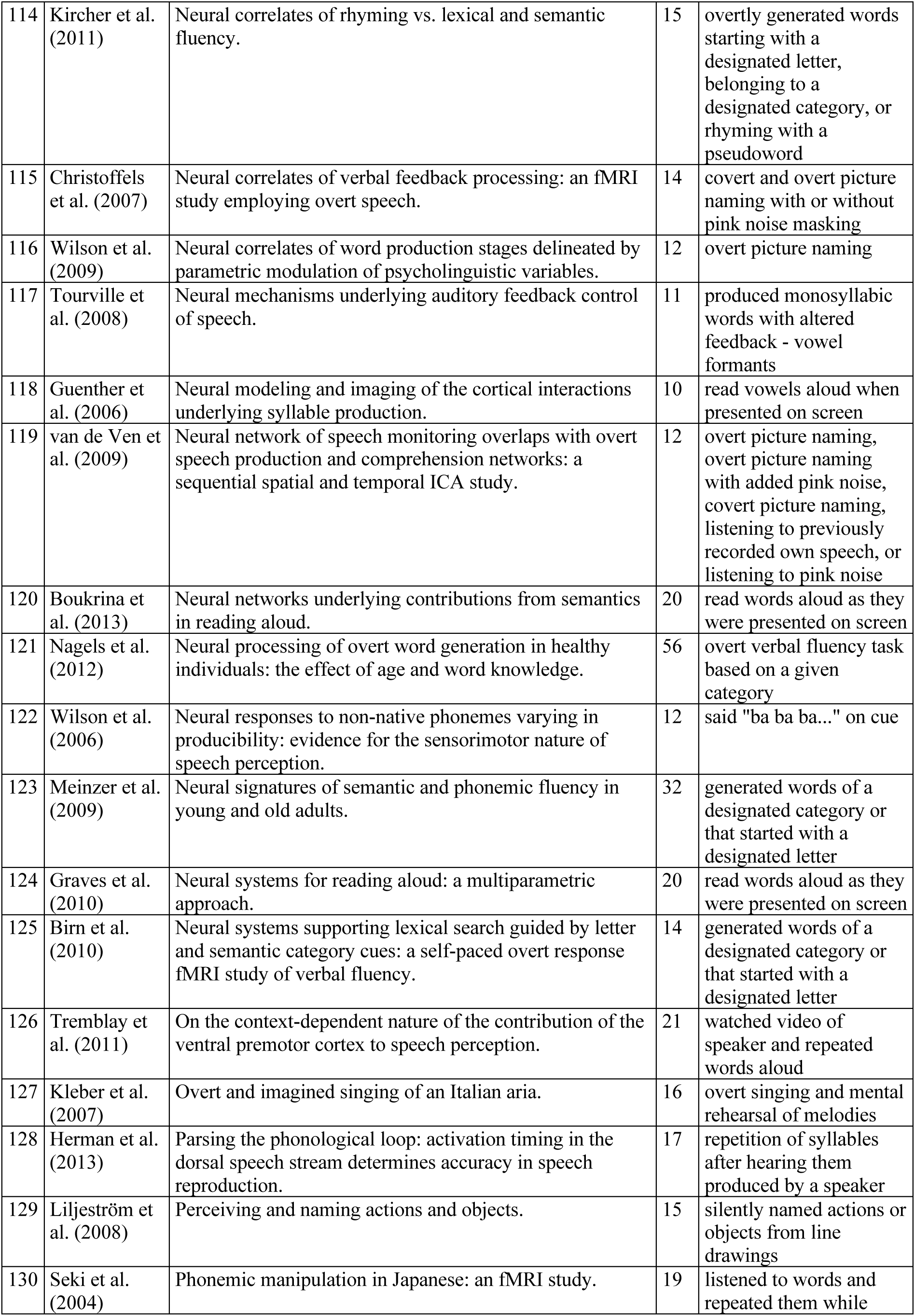
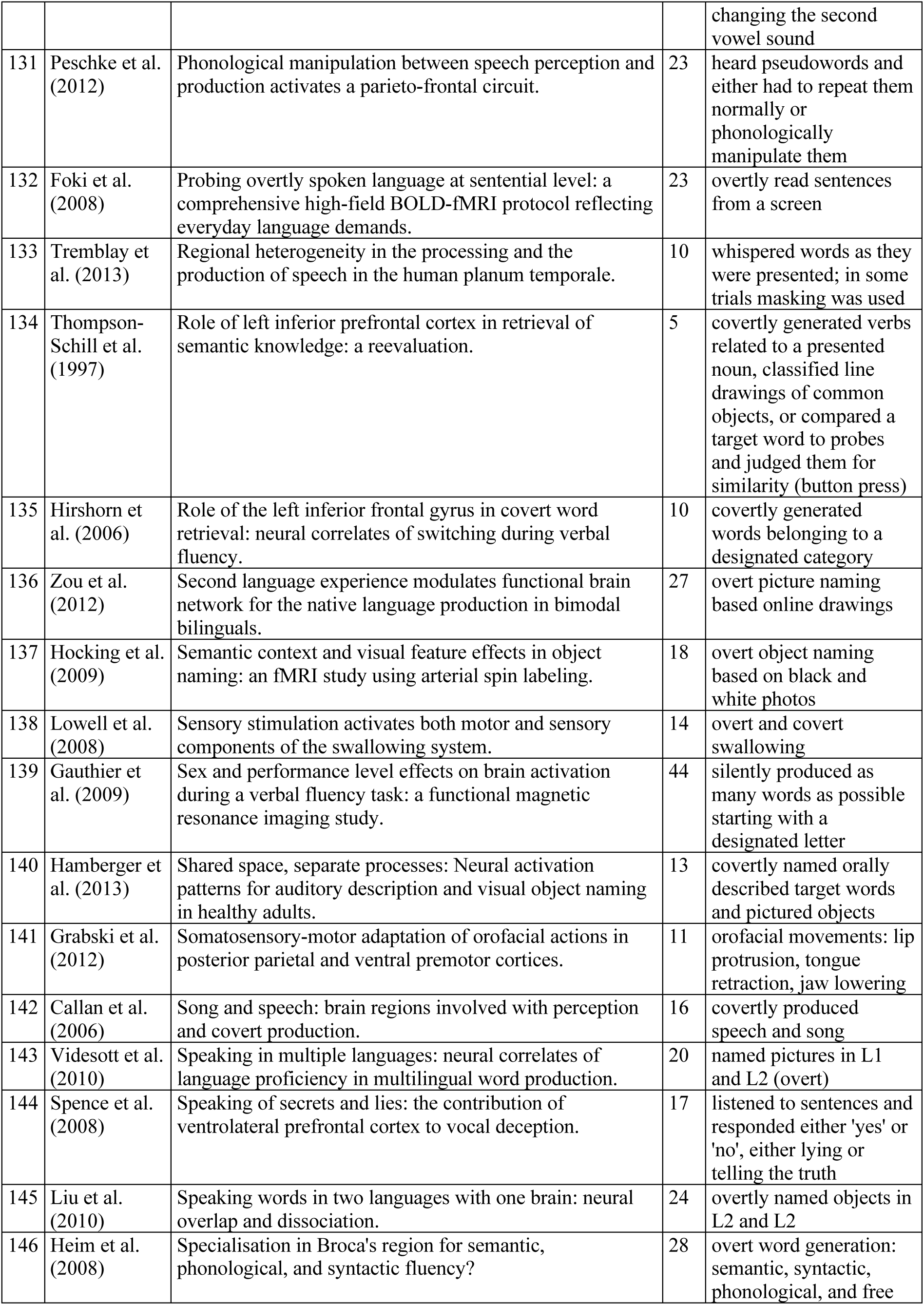
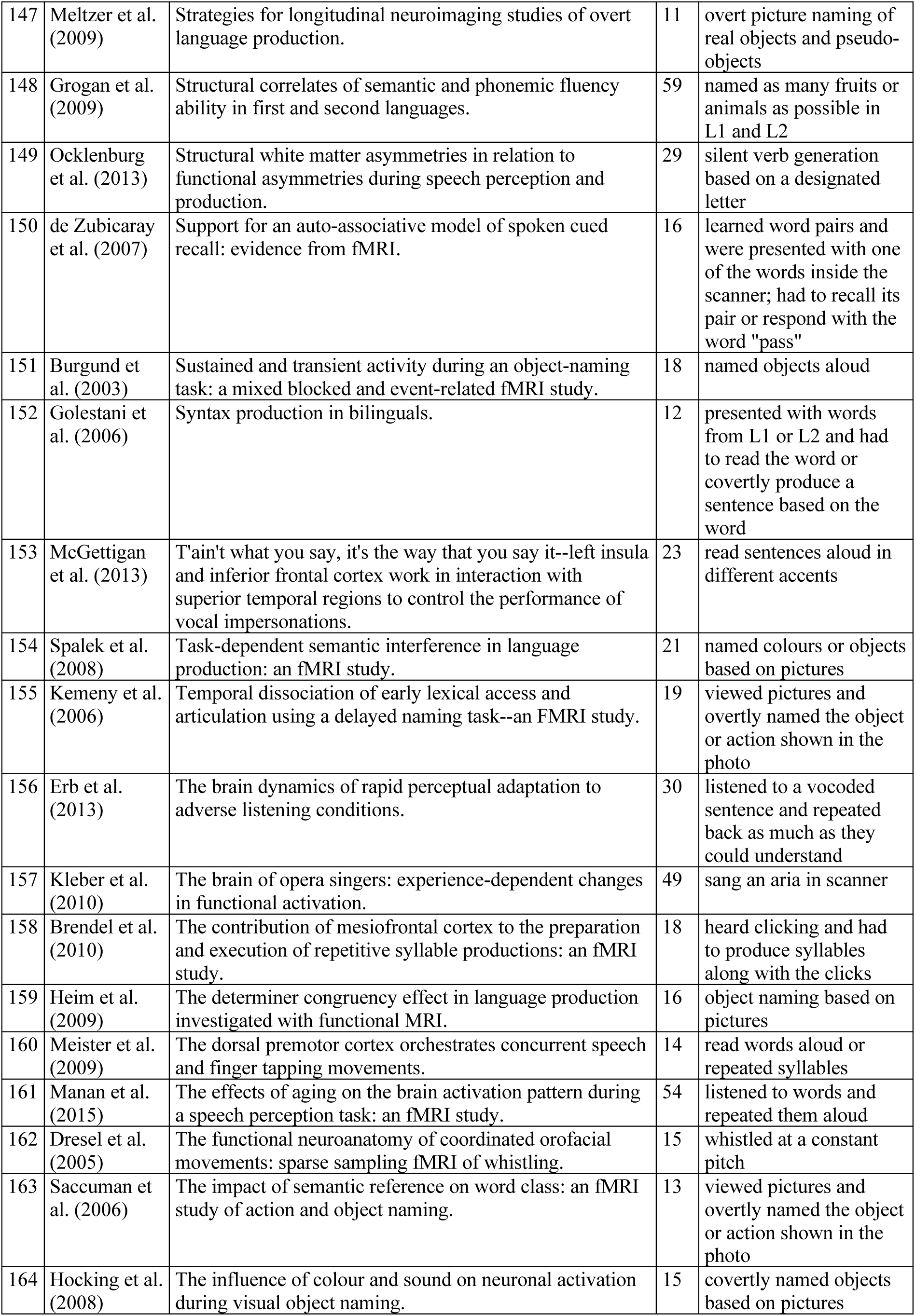
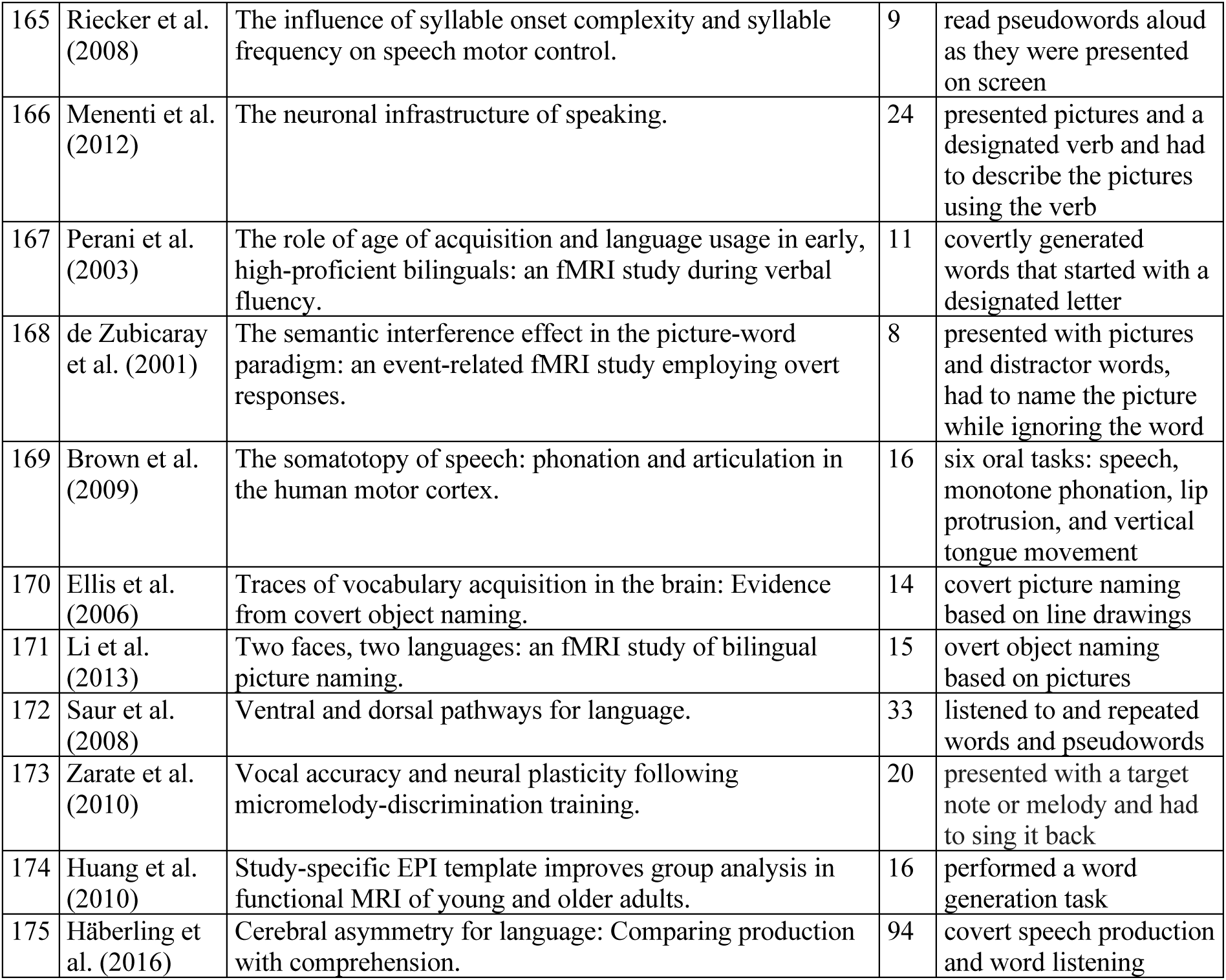
Studies involving speech production reporting cerebellum activation.

| # | Authors | Title | n | Task |
| --- | --- | --- | --- | --- |
| 1 | Van de Putte et al. (2018) | Neural overlap of L1 and L2 semantic representations across visual and auditory modalities: a decoding approach. | 22 | 3 tasks: picture naming, word reading, and word listening |
| 2 | Sumiya et al. (2017) | Brain networks of social action-outcome contingency: The role of the ventral striatum in integrating signals from the sensory cortex and medial prefrontal cortex. | 38 | read punchlines to jokes aloud |
| 3 | Lurito et al. (2000) | Comparison of rhyming and word generation with FMRI. | 5 | subvocally articulated of words on a screen |
| 4 | Beaty et al. (2017) | Creative constraints: Brain activity and network dynamics underlying semantic interference during idea production. | 24 | word generation and cued recall tasks (spoken aloud) |
| 5 | Reilly et al. (2016) | Does the sound of a barking dog activate its corresponding visual form? An fMRI investigation of modality-specific semantic access. | 18 | named objects based on sounds, images, or point-light motion |
| 6 | Takashima et al. (2016) | Emergence of representations through repeated training on pronouncing novel letter combinations leads to efficient reading. | 25 | pronounced pseudowords written in a foreign script |
| 7 | Carey et al. (2017) | Functional brain outcomes of L2 speech learning emerge during sensorimotor transformation. | 28 | produced words containing native/non-native vowels |
| 8 | Pattamadilok et al. (2015) | How Early Does the Brain Distinguish between Regular Words, Irregular Words, and Pseudowords during the Reading Process? Evidence from Neurochronometric TMS. | 92 | read aloud while undergoing TMS |
| 9 | Danelli et al. (2017) | How many deficits in the same dyslexic brains? A behavioural and fMRI assessment of comorbidity in adult dyslexics. | 23 | spoonerism, digit naming, and picture naming tasks (aloud) |
| 10 | Ghazi-Saidi et al. (2015) | How native-like can you possibly get: fMRI evidence for processing accent. | 12 | overt picture naming |
| 11 | Flanagin et al. (2017) | Human Exploration of Enclosed Spaces through Echolocation. | 11 | produced calls and listened to the echoes |
| 12 | Quinn et al. (2017) | Learning and retrieving holistic and componential visual-verbal associations in reading and object naming. | 20 | read words and named objects aloud |
| 13 | Weiss et al. (2015) | Many ways to read your vowels--Neural processing of diacritics and vowel letters in Hebrew. | 18 | read words and pseudowords aloud |
| 14 | Markiewicz et al. (2016) | Mapping the cortical representation of speech sounds in a syllable repetition task. | 14 | listened to syllables and repeated them out loud |
| 15 | Hung et al. (2015) | Neural correlates for naming disadvantage of the dominant language in bilingual word production. | 21 | named pictures while switching between Chinese and English |
| 16 | Hung et al. (2015) | Neural correlates of merging number words. | 40 | read number words aloud |
| 17 | Geranmayeh et al. (2014) | Overlapping networks engaged during spoken language production and its cognitive control. | 24 | named pictures and counted aloud |
| 18 | Venezia et al. (2016) | Perception drives production across sensory modalities: A network for sensorimotor integration of visual speech. | 20 | covertly rehearsed sequences of syllables |
| 19 | Gurd et al. (2002) | Posterior parietal cortex is implicated in continuous switching between verbal fluency tasks: an fMRI study with clinical implications. | 11 | covert word generation within a semantic category |
| 20 | Kurland et al. (2018) | Practice effects in healthy older adults: Implications for treatment-induced neuroplasticity in Aphasia. | 18 | object naming from line drawings |
| 21 | Berkovich-Ohana et al. (2015) | Repetitive speech elicits widespread deactivation in the human cortex: the "Mantra" effect? | 23 | covert word repetition - received auditory cues |
| 22 | Purcell et al. (2017) | Shared orthographic neuronal representations for spelling and reading. | 16 | covertly read or spelled (using keyboard) words presented auditorily |
| 23 | Provost et al. (2016) | The effect of aging on the brain network for exception word reading. | 32 | read pseudowords aloud |
| 24 | Reverberi et al. (2018) | The neural basis of free language choice in bilingual speakers: Disentangling language choice and language execution. | 15 | named objects based on line drawings while randomly switching between English and German |
| 25 | Belyk et al. (2016) | The Neural Basis of Vocal Pitch Imitation in Humans. | 13 | listened to and repeated melodies |
| 26 | Bilodeau-Mercure et al. (2015) | The neurobiology of speech perception decline in aging. | 22 | listened to and repeated syllables at different levels of intelligibility |
| 27 | Thothathiri et al. (2015) | Ventral and dorsal streams for choosing word order during sentence production. | 18 | viewed a scene and provided verbal descriptions of the scene using a pre-determined verb |
| 28 | Tian et al. (2016) | Mental imagery of speech implicates two mechanisms of perceptual reactivation. | 18 | overtly generated a syllable, imagined generating the syllable, or passively listened to the syllable |
| 29 | González-García et al. (2016) | Neural activity related to discrimination and vocal production of consonant and dissonant musical intervals. | 11 | passively listened to or hummed back musical intervals |
| 30 | McCrea (2007) | A functional magnetic resonance imaging study of the body schema using full human line-drawing figures in an on-line verbal naming and localization task of single body part words. | 9 | body part naming |
| 31 | Moore et al. (1999) | A functional neuroimaging study of the variables that generate category-specific object processing differences. | 8 | object naming based on line drawings (silently mouthed the words) |
| 32 | Brown et al. (2008) | A larynx area in the human motor cortex. | 16 | vocalization tasks: phonation (singing on a specific vowel), performing glottal stops, lip protrusion, and vertical tongue movement |
| 33 | Zubicaray et al. (2014) | A perfusion fMRI investigation of thematic and categorical context effects in the spoken production of object names. | 28 | picture naming presented in either homogenous or heterogenous contexts |
| 34 | Thürling et al. (2011) | Activation of the dentate nucleus in a verb generation task: A 7T MRI study. | 17 | reading aloud, silent reading, silent verb generation, and automatized inner speech |
| 35 | Sato et al. (2014) | Adaptive coding of orofacial and speech actions in motor and somatosensory spaces with and without overt motor behavior. | 14 | silent orofacial movements, audible orofacial movements, and syllable production |
| 36 | Pichon et al. (2013) | Affective and sensorimotor components of emotional prosody generation. | 20 | cued to read aloud with different emotional prosody |
| 37 | Shuster et al. (2005) | An fMRI investigation of covertly and overtly produced mono- and multisyllabic words. | 10 | produced monosyllabic and multisyllabic words covertly or overtly |
| 38 | Yoon et al. (2006) | An fMRI study of Chinese character reading and picture naming by native Korean speakers. | 14 | picture naming |
| 39 | Garn et al. (2009) | An fMRI study of sex differences in brain activation during object naming. | 26 | picture naming based on line drawings: living, non-living, or scrambled images |
| 40 | Piai et al. (2013) | Attention for speaking: domain-general control from the anterior cingulate cortex in spoken word production. | 23 | picture-word interference task, Stroop task (overt naming) |
| 41 | Matchin et al. (2014) | Audiovisual speech integration does not rely on the motor system: evidence from articulatory suppression, the McGurk effect, and fMRI. | 50 | watched videos of speaker producing /pa/, /ta/ and /ka/; had to report the sound they heard while silently articulating “/pa/.../ba/” |
| 42 | Raboyeau et al. (2010) | Brain activation and lexical learning: the impact of learning phase and word type. | 10 | overt object naming in French and Spanish |
| 43 | Hervais-Adelman et al. (2015) | Brain functional plasticity associated with the emergence of expertise in extreme language control. | 35 | listened to French and English sentences and were asked to either listen, shadow (repeat), or interpret |
| 44 | Hwang et al. (2009) | Category-specific activations during word generation reflect experiential sensorimotor modalities. | 13 | covert or overt word generation |
| 45 | Frings et al. (2006) | Cerebellar involvement in verb generation: an fMRI study. | 16 | word reading or verb generation, overt or covert |
| 46 | de Zubicaray et al. (2000) | Cerebral regions associated with verbal response initiation, suppression and strategy use. | 8 | overt category naming, word length judgement |
| 47 | Chen et al. (2005) | Cerebrocerebellar networks during articulatory rehearsal and verbal working memory tasks. | 17 | subvocally read letters |
| 48 | Rauschecker et al. (2008) | Changes in neural activity associated with learning to articulate novel auditory pseudowords by covert repetition. | 14 | covertly repeated pseudowords following auditory presentation |
| 49 | Snyder et al. (2011) | Choosing our words: retrieval and selection processes recruit shared neural substrates in left ventrolateral prefrontal cortex. | 18 | verb generation after being presented with a noun |
| 50 | Soros et al. (2006) | Clustered functional MRI of overt speech production. | 9 | repeated sublexical speech sounds and performed oral movements without vocalization |
| 51 | Chang et al. (2009) | Common neural substrates support speech and non-speech vocal tract gestures. | 34 | produced a variety of articulatory gestures (glottal attacks, lip closures, bilabial bursts, tongue thrusts) |
| 52 | Halari et al. (2006) | Comparable fMRI activity with differential behavioural performance on mental rotation and overt verbal fluency tasks in healthy men and women. | 19 | overt verbal fluency task |
| 53 | Peck et al. (2004) | Comparison of baseline conditions to investigate syntactic production using functional magnetic resonance imaging. | 10 | silent sentence generation based on line drawings, passive object viewing, and object naming |
| 54 | Kemeny et al. (2005) | Comparison of continuous overt speech fMRI using BOLD and arterial spin labeling. | 6 | overt continuous vowel/consonant production |
| 55 | Humphreys et al. (2014) | Competitive mechanisms in sentence processing: common and distinct production and reading comprehension networks linked to the prefrontal cortex. | 17 | covertly read and completed sentence fragments |
| 56 | Cai et al. (2013) | Complementary hemispheric specialization for language production and visuospatial attention. | 31 | silent word generation |
| 57 | Abel et al. (2009) | Correct and erroneous picture naming responses in healthy subjects. | 34 | overt picture naming |
| 58 | Benedek et al. (2014) | Creating metaphors: the neural basis of figurative language production. | 28 | overt metaphor generation |
| 59 | Trapp et al. (2014) | Different neural capacity limitations for articulatory and non-articulatory maintenance of verbal information. | 14 | subvocally rehearsed letters (articulatory and non-articulatory conditions) |
| 60 | Nash et al. (2009) | Differential activation of the human trigeminal nuclear complex by noxious and non-noxious orofacial stimulation. | 30 | orofacial stimulation - participants reported and described their experience of pain while in the scanner |
| 61 | Liu et al. (2006) | Dissociation in the neural basis underlying Chinese tone and vowel production. | 10 | overt word naming |
| 62 | Glikmann-Johnston et al. (2015) | Distinct functional connectivity of the hippocampus during semantic and phonemic fluency. | 18 | covert word generation (semantic and phonemic blocks) |
| 63 | Taylor et al. (2014) | Distinct neural specializations for learning to read words and name objects. | 19 | overt and covert object name retrieval |
| 64 | Peeva et al. (2010) | Distinct representations of phonemes, syllables, and supra-syllabic sequences in the speech production network. | 18 | read pseudowords aloud |
| 65 | Palomar-García et al. (2015) | Do bilinguals show neural differences with monolinguals when processing their native language? | 44 | overt picture naming based online drawings |
| 66 | Yoo et al. (2012) | Dual routes for verbal repetition: articulation-based and acoustic-phonetic codes for pseudoword and word repetition, respectively. | 22 | listened to words with ambiguous vowel sounds and overtly repeated what they heard (overt) |
| 67 | Wildgruber et al. (1999) | Dynamic pattern of brain activation during sequencing of word strings evaluated by fMRI. | 18 | covertly recited the months of the year in forwards and backwards order |
| 68 | Grabowski et al. (2003) | Effects of gender on blood flow correlates of naming concrete entities. | 62 | overt object naming |
| 69 | Basho et al. (2007) | Effects of generation mode in fMRI adaptations of semantic fluency: paced production and overt speech. | 12 | overt or covert word generation based on a given category |
| 70 | Bavelier et al. (2008) | Encoding, rehearsal, and recall in signers and speakers: shared network but differential engagement. | 20 | speech and sign (ASL) production |
| 71 | Peck et al. (2009) | Event-related functional MRI investigation of vocal pitch variation. | 12 | produced "uh" at three different pitches |
| 72 | Zarate et al. (2008) | Experience-dependent neural substrates involved in vocal pitch regulation during singing. | 24 | singing pitches aloud with unaltered or pitch-shifted auditory feedback |
| 73 | Josse et al. (2008) | Explaining function with anatomy: language lateralization and corpus callosum size. | 74 | overt picture naming |
| 74 | Wattendorf et al. (2013) | Exploration of the neural correlates of ticklish laughter by functional magnetic resonance imaging. | 18 | tickled inside scanner and were either asked to inhibit laughter or were encouraged to make vocalizations |
| 75 | Wilson et al. (2011) | Finding your voice: a singing lesson from functional imaging. | 26 | covert singing and lexical retrieval |
| 76 | Golfinopoulou et al. (2011) | fMRI investigation of unexpected somatosensory feedback perturbation during speech. | 13 | read pseudowords aloud while wearing a device that disrupted speech movements |
| 77 | Hervais-Adelman et al. (2015) | fMRI of Simultaneous Interpretation Reveals the Neural Basis of Extreme Language Control. | 43 | covertly repeated sentences and translated sentences into native language |
| 78 | Grande et al. (2012) | From a concept to a word in a syntactically complete sentence: an fMRI study on spontaneous language production in an overt picture description task. | 22 | described scenes depicted by line drawings in detail (overt) |
| 79 | Green et al. (2015) | Frontopolar activity and connectivity support dynamic conscious augmentation of creative state. | 63 | presented with noun and had to generate a related verb and were cued to think creatively |
| 80 | Tremblay et al. (2013) | Functional and structural aging of the speech sensorimotor neural system: functional magnetic resonance imaging evidence. | 39 | watched audiovisual movie of speaker producing words and repeated what they heard |
| 81 | Abrahams et al. (2003) | Functional magnetic resonance imaging of verbal fluency and confrontation naming using compressed image acquisition to permit overt responses. | 18 | overt word generation based on an auditorily presented letter |
| 82 | Grabski et al. (2012) | Functional MRI assessment of orofacial articulators: neural correlates of lip, jaw, larynx, and tongue movements. | 13 | overt vowel production and three orofacial motor tasks |
| 83 | Schulz et al. (2005) | Functional neuroanatomy of human vocalization: an H215O PET study. | 20 | voiced and whispered narrative production |
| 84 | Zheng et al. (2010) | Functional overlap between regions involved in speech perception and in monitoring one's own voice during speech production. | 21 | producing whispered speech while either hearing this speech through headphones or while hearing masking noise |
| 85 | Vitali et al. (2005) | Generating animal and tool names: an fMRI study of effective connectivity. | 10 | covertly generated words belonging to the "animal" and "tool" categories |
| 86 | Yu et al. (2013) | Gum chewing inhibits the sensory processing and the propagation of stress-related information in a brain network. | 24 | chewed gum while listening to noise stimuli |
| 87 | Loucks et al. (2007) | Human brain activation during phonation and exhalation: common volitional control for two upper airway functions. | 12 | repeated of syllables while exhaling |
| 88 | Chang et al. (2014) | Implicit target substitution and sequencing for lexical tone production in Chinese: an FMRI study. | 21 | listened to tones and produced syllables |
| 89 | Hartwigsen et al. (2013) | Increased facilitatory connectivity from the pre-SMA to the left dorsal premotor cortex during pseudoword repetition. | 17 | repeated auditorily or visually presented pseudowords (overt) |
| 90 | McGettigan et al. (2015) | Individual differences in laughter perception reveal roles for mentalizing and sensorimotor systems in the evaluation of emotional authenticity. | 21 | smiled and wrinkled noses on cue |
| 91 | Collina et al. (2014) | Inside the syntactic box: the neural correlates of the functional and positional level in covert sentence production. | 20 | presented with three words and either had to covertly read them or covertly produce a sentence containing them |
| 92 | Seghier et al. (2008) | Inter-subject variability in the use of two different neuronal networks for reading aloud familiar words. | 43 | read object names aloud |
| 93 | Crescentini et al. (2010) | Item retrieval and competition in noun and verb generation: an FMRI study. | 14 | overt verb and noun generation |
| 94 | Hu et al. (2013) | Language aptitude for pronunciation in advanced second language (L2) learners: behavioural predictors and neural substrates. | 8 | listened to a sentence in L1 or L2 and repeated the sentence |
| 95 | Kell et al. (2011) | Lateralization of speech production starts in sensory cortices--a possible sensory origin of cerebral left dominance for speech. | 42 | covertly or overtly read sentences as they appeared on a screen / made orofacial movements (two experiments) |
| 96 | Heim et al. (2009) | Left cytoarchitectonic area 44 supports selection in the mental lexicon during language production. | 23 | picture naming |
| 97 | Guo et al. (2011) | Local and global inhibition in bilingual word production: fMRI evidence from Chinese-English bilinguals. | 24 | overtly named pictures in L1 and L2 |
| 98 | Marsolais et al. (2015) | Marginal neurofunctional changes in high-performing older adults in a verbal fluency task. | 28 | verbal fluency: word appears on screen and participants said aloud as many words as they could belonging to that category |
| 99 | Vogel et al. (2013) | Matching is not naming: a direct comparison of lexical manipulations in explicit and implicit reading tasks. | 22 | read words aloud as they were presented on screen |
| 100 | van Turennout et al. (2003) | Modulation of neural activity during object naming: effects of time and practice. | 10 | covert object naming based on line drawings |
| 101 | Wong et al. (2011) | Motor control of jaw movements: An fMRI study of parafunctional clench and grind behavior. | 19 | performed orofacial movements (clench and grind on cue) |
| 102 | Vingerhoets et al. (2003) | Multilingualism: an fMRI study. | 12 | covertly generated words starting with a designated letter; covert picture naming; silent reading |
| 103 | Zheng et al. (2013) | Multivoxel patterns reveal functionally differentiated networks underlying auditory feedback processing of speech. | 20 | produced monosyllabic words with altered auditory feedback |
| 104 | Okada et al. (2000) | Naming of animals and tools: a functional magnetic resonance imaging study of categorical differences in the human brain areas commonly used for naming visually presented objects. | 12 | silent object naming based on drawings |
| 105 | Troiani et al. (2008) | Narrative speech production: an fMRI study using continuous arterial spin labeling. | 15 | described pictures aloud |
| 106 | Berken et al. (2015) | Neural activation in speech production and reading aloud in native and non-native languages. | 47 | read sentences aloud in L1 and L2 |
| 107 | Garbin et al. (2011) | Neural bases of language switching in high and early proficient bilinguals. | 19 | picture naming while switching between L1 and L2 |
| 108 | Lei et al. (2014) | Neural basis of language switching in the brain: fMRI evidence from Korean-Chinese early bilinguals. | 8 | covertly generated properties of visually presented words |
| 109 | de Diego et al. (2006) | Neural circuits subserving the retrieval of stems and grammatical features in regular and irregular verbs. | 12 | presented with verbs and had to either covertly produce the verb's present tense or read the verb |
| 110 | Consonni et al. (2013) | Neural convergence for language comprehension and grammatical class production in highly proficient bilinguals is independent of age of acquisition. | 24 | listened to sentences and overtly generated the object or action word in the sentence |
| 111 | AbdulSabur et al. (2014) | Neural correlates and network connectivity underlying narrative production and comprehension: a combined fMRI and PET study. | 18 | retold a story or repeated a nursery rhyme (overt) |
| 112 | Park et al. (2011) | Neural correlates in the processing of phoneme-level complexity in vowel production. | 14 | listened to speech sounds and performed orofacial movements without vocalization |
| 113 | Wende et al. (2012) | Neural correlates of continuous causal word generation. | 18 | generated words that were causally related to a cue word, were associated with cue, or rhymed with cue |
| 114 | Kircher et al. (2011) | Neural correlates of rhyming vs. lexical and semantic fluency. | 15 | overtly generated words starting with a designated letter, belonging to a designated category, or rhyming with a pseudoword |
| 115 | Christoffels et al. (2007) | Neural correlates of verbal feedback processing: an fMRI study employing overt speech. | 14 | covert and overt picture naming with or without pink noise masking |
| 116 | Wilson et al. (2009) | Neural correlates of word production stages delineated by parametric modulation of psycholinguistic variables. | 12 | overt picture naming |
| 117 | Tourville et al. (2008) | Neural mechanisms underlying auditory feedback control of speech. | 11 | produced monosyllabic words with altered feedback - vowel formants |
| 118 | Guenther et al. (2006) | Neural modeling and imaging of the cortical interactions underlying syllable production. | 10 | read vowels aloud when presented on screen |
| 119 | van de Ven et al. (2009) | Neural network of speech monitoring overlaps with overt speech production and comprehension networks: a sequential spatial and temporal ICA study. | 12 | overt picture naming, overt picture naming with added pink noise, covert picture naming, listening to previously recorded own speech, or listening to pink noise |
| 120 | Boukrina et al. (2013) | Neural networks underlying contributions from semantics in reading aloud. | 20 | read words aloud as they were presented on screen |
| 121 | Nagels et al. (2012) | Neural processing of overt word generation in healthy individuals: the effect of age and word knowledge. | 56 | overt verbal fluency task based on a given category |
| 122 | Wilson et al. (2006) | Neural responses to non-native phonemes varying in producibility: evidence for the sensorimotor nature of speech perception. | 12 | said "ba ba ba..." on cue |
| 123 | Meinzer et al. (2009) | Neural signatures of semantic and phonemic fluency in young and old adults. | 32 | generated words of a designated category or that started with a designated letter |
| 124 | Graves et al. (2010) | Neural systems for reading aloud: a multiparametric approach. | 20 | read words aloud as they were presented on screen |
| 125 | Birn et al. (2010) | Neural systems supporting lexical search guided by letter and semantic category cues: a self-paced overt response fMRI study of verbal fluency. | 14 | generated words of a designated category or that started with a designated letter |
| 126 | Tremblay et al. (2011) | On the context-dependent nature of the contribution of the ventral premotor cortex to speech perception. | 21 | watched video of speaker and repeated words aloud |
| 127 | Kleber et al. (2007) | Overt and imagined singing of an Italian aria. | 16 | overt singing and mental rehearsal of melodies |
| 128 | Herman et al. (2013) | Parsing the phonological loop: activation timing in the dorsal speech stream determines accuracy in speech reproduction. | 17 | repetition of syllables after hearing them produced by a speaker |
| 129 | Liljeström et al. (2008) | Perceiving and naming actions and objects. | 15 | silently named actions or objects from line drawings |
| 130 | Seki et al. (2004) | Phonemic manipulation in Japanese: an fMRI study. | 19 | listened to words and repeated them while changing the second vowel sound |
| 131 | Peschke et al. (2012) | Phonological manipulation between speech perception and production activates a parieto-frontal circuit. | 23 | heard pseudowords and either had to repeat them normally or phonologically manipulate them |
| 132 | Foki et al. (2008) | Probing overtly spoken language at sentential level: a comprehensive high-field BOLD-fMRI protocol reflecting everyday language demands. | 23 | overtly read sentences from a screen |
| 133 | Tremblay et al. (2013) | Regional heterogeneity in the processing and the production of speech in the human planum temporale. | 10 | whispered words as they were presented; in some trials masking was used |
| 134 | Thompson-Schill et al. (1997) | Role of left inferior prefrontal cortex in retrieval of semantic knowledge: a reevaluation. | 5 | covertly generated verbs related to a presented noun, classified line drawings of common objects, or compared a target word to probes and judged them for similarity (button press) |
| 135 | Hirshorn et al. (2006) | Role of the left inferior frontal gyrus in covert word retrieval: neural correlates of switching during verbal fluency. | 10 | covertly generated words belonging to a designated category |
| 136 | Zou et al. (2012) | Second language experience modulates functional brain network for the native language production in bimodal bilinguals. | 27 | overt picture naming based online drawings |
| 137 | Hocking et al. (2009) | Semantic context and visual feature effects in object naming: an fMRI study using arterial spin labeling. | 18 | overt object naming based on black and white photos |
| 138 | Lowell et al. (2008) | Sensory stimulation activates both motor and sensory components of the swallowing system. | 14 | overt and covert swallowing |
| 139 | Gauthier et al. (2009) | Sex and performance level effects on brain activation during a verbal fluency task: a functional magnetic resonance imaging study. | 44 | silently produced as many words as possible starting with a designated letter |
| 140 | Hamberger et al. (2013) | Shared space, separate processes: Neural activation patterns for auditory description and visual object naming in healthy adults. | 13 | covertly named orally described target words and pictured objects |
| 141 | Grabski et al. (2012) | Somatosensory-motor adaptation of orofacial actions in posterior parietal and ventral premotor cortices. | 11 | orofacial movements: lip protrusion, tongue retraction, jaw lowering |
| 142 | Callan et al. (2006) | Song and speech: brain regions involved with perception and covert production. | 16 | covertly produced speech and song |
| 143 | Videsott et al. (2010) | Speaking in multiple languages: neural correlates of language proficiency in multilingual word production. | 20 | named pictures in L1 and L2 (overt) |
| 144 | Spence et al. (2008) | Speaking of secrets and lies: the contribution of ventrolateral prefrontal cortex to vocal deception. | 17 | listened to sentences and responded either 'yes' or 'no', either lying or telling the truth |
| 145 | Liu et al. (2010) | Speaking words in two languages with one brain: neural overlap and dissociation. | 24 | overtly named objects in L1 and L2 |
| 146 | Heim et al. (2008) | Specialisation in Broca's region for semantic, phonological, and syntactic fluency? | 28 | overt word generation: semantic, syntactic, phonological, and free |
| 147 | Meltzer et al. (2009) | Strategies for longitudinal neuroimaging studies of overt language production. | 11 | overt picture naming of real objects and pseudo-objects |
| 148 | Grogan et al. (2009) | Structural correlates of semantic and phonemic fluency ability in first and second languages. | 59 | named as many fruits or animals as possible in L1 and L2 |
| 149 | Ocklenburg et al. (2013) | Structural white matter asymmetries in relation to functional asymmetries during speech perception and production. | 29 | silent verb generation based on a designated letter |
| 150 | de Zubicaray et al. (2007) | Support for an auto-associative model of spoken cued recall: evidence from fMRI. | 16 | learned word pairs and were presented with one of the words inside the scanner; had to recall its pair or respond with the word "pass" |
| 151 | Burgund et al. (2003) | Sustained and transient activity during an object-naming task: a mixed blocked and event-related fMRI study. | 18 | named objects aloud |
| 152 | Golestani et al. (2006) | Syntax production in bilinguals. | 12 | presented with words from L1 or L2 and had to read the word or covertly produce a sentence based on the word |
| 153 | McGettigan et al. (2013) | T'ain't what you say, it's the way that you say it--left insula and inferior frontal cortex work in interaction with superior temporal regions to control the performance of vocal impersonations. | 23 | read sentences aloud in different accents |
| 154 | Spalek et al. (2008) | Task-dependent semantic interference in language production: an fMRI study. | 21 | named colours or objects based on pictures |
| 155 | Kemeny et al. (2006) | Temporal dissociation of early lexical access and articulation using a delayed naming task--an FMRI study. | 19 | viewed pictures and overtly named the object or action shown in the photo |
| 156 | Erb et al. (2013) | The brain dynamics of rapid perceptual adaptation to adverse listening conditions. | 30 | listened to a vocoded sentence and repeated back as much as they could understand |
| 157 | Kleber et al. (2010) | The brain of opera singers: experience-dependent changes in functional activation. | 49 | sang an aria in scanner |
| 158 | Brendel et al. (2010) | The contribution of mesiofrontal cortex to the preparation and execution of repetitive syllable productions: an fMRI study. | 18 | heard clicking and had to produce syllables along with the clicks |
| 159 | Heim et al. (2009) | The determiner congruency effect in language production investigated with functional MRI. | 16 | object naming based on pictures |
| 160 | Meister et al. (2009) | The dorsal premotor cortex orchestrates concurrent speech and finger tapping movements. | 14 | read words aloud or repeated syllables |
| 161 | Manan et al. (2015) | The effects of aging on the brain activation pattern during a speech perception task: an fMRI study. | 54 | listened to words and repeated them aloud |
| 162 | Dresel et al. (2005) | The functional neuroanatomy of coordinated orofacial movements: sparse sampling fMRI of whistling. | 15 | whistled at a constant pitch |
| 163 | Saccuman et al. (2006) | The impact of semantic reference on word class: an fMRI study of action and object naming. | 13 | viewed pictures and overtly named the object or action shown in the photo |
| 164 | Hocking et al. (2008) | The influence of colour and sound on neuronal activation during visual object naming. | 15 | covertly named objects based on pictures |
| 165 | Riecker et al. (2008) | The influence of syllable onset complexity and syllable frequency on speech motor control. | 9 | read pseudowords aloud as they were presented on screen |
| 166 | Menenti et al. (2012) | The neuronal infrastructure of speaking. | 24 | presented pictures and a designated verb and had to describe the pictures using the verb |
| 167 | Perani et al. (2003) | The role of age of acquisition and language usage in early, high-proficient bilinguals: an fMRI study during verbal fluency. | 11 | covertly generated words that started with a designated letter |
| 168 | de Zubicaray et al. (2001) | The semantic interference effect in the picture-word paradigm: an event-related fMRI study employing overt responses. | 8 | presented with pictures and distractor words, had to name the picture while ignoring the word |
| 169 | Brown et al. (2009) | The somatotopy of speech: phonation and articulation in the human motor cortex. | 16 | six oral tasks: speech, monotone phonation, lip protrusion, and vertical tongue movement |
| 170 | Ellis et al. (2006) | Traces of vocabulary acquisition in the brain: Evidence from covert object naming. | 14 | covert picture naming based on line drawings |
| 171 | Li et al. (2013) | Two faces, two languages: an fMRI study of bilingual picture naming. | 15 | overt object naming based on pictures |
| 172 | Saur et al. (2008) | Ventral and dorsal pathways for language. | 33 | listened to and repeated words and pseudowords |
| 173 | Zarate et al. (2010) | Vocal accuracy and neural plasticity following micromelody-discrimination training. | 20 | presented with a target note or melody and had to sing it back |
| 174 | Huang et al. (2010) | Study-specific EPI template improves group analysis in functional MRI of young and older adults. | 16 | performed a word generation task |
| 175 | Häberling et al. (2016) | Cerebral asymmetry for language: Comparing production with comprehension. | 94 | covert speech production and word listening |

**Table S3.**
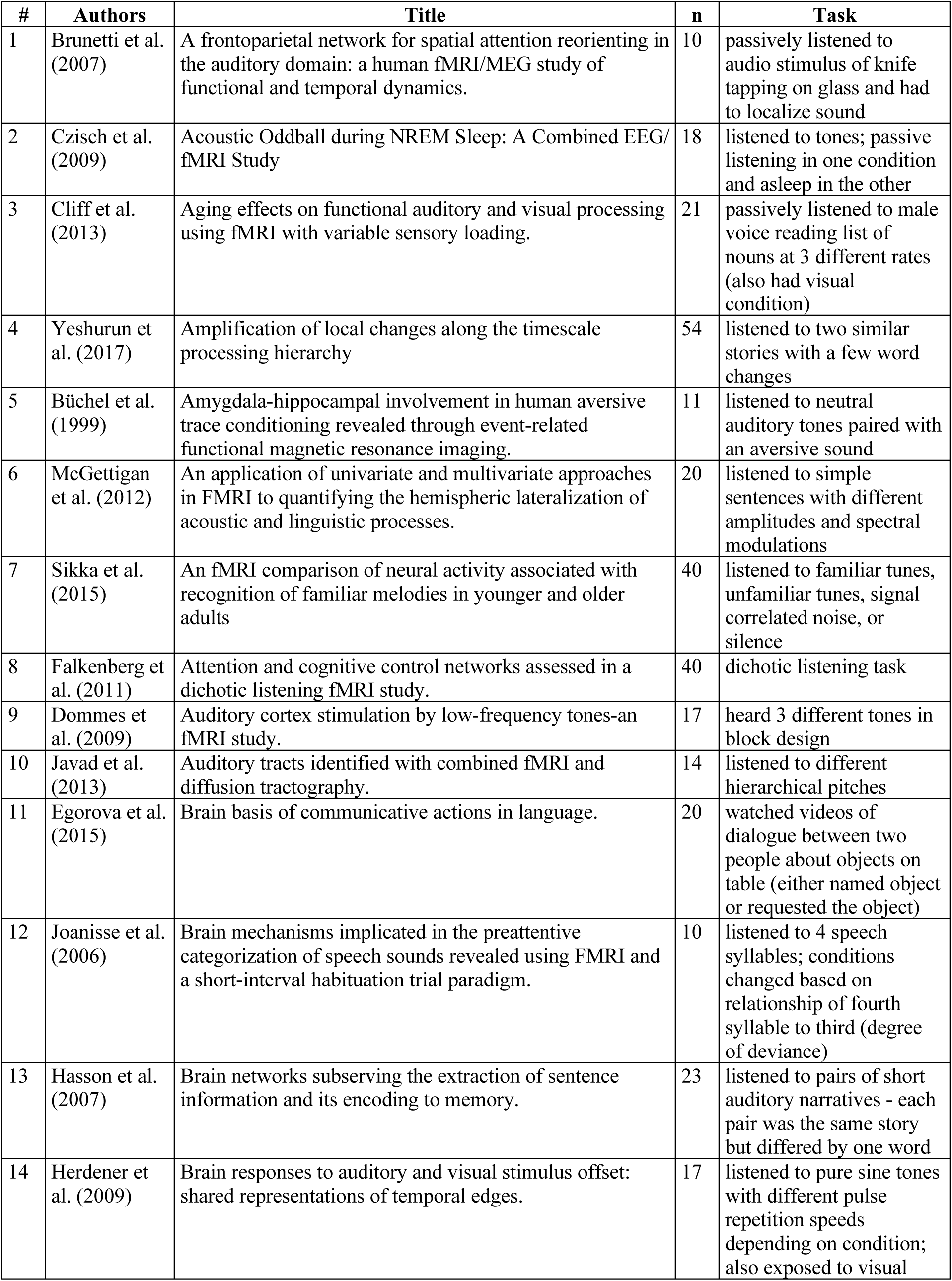
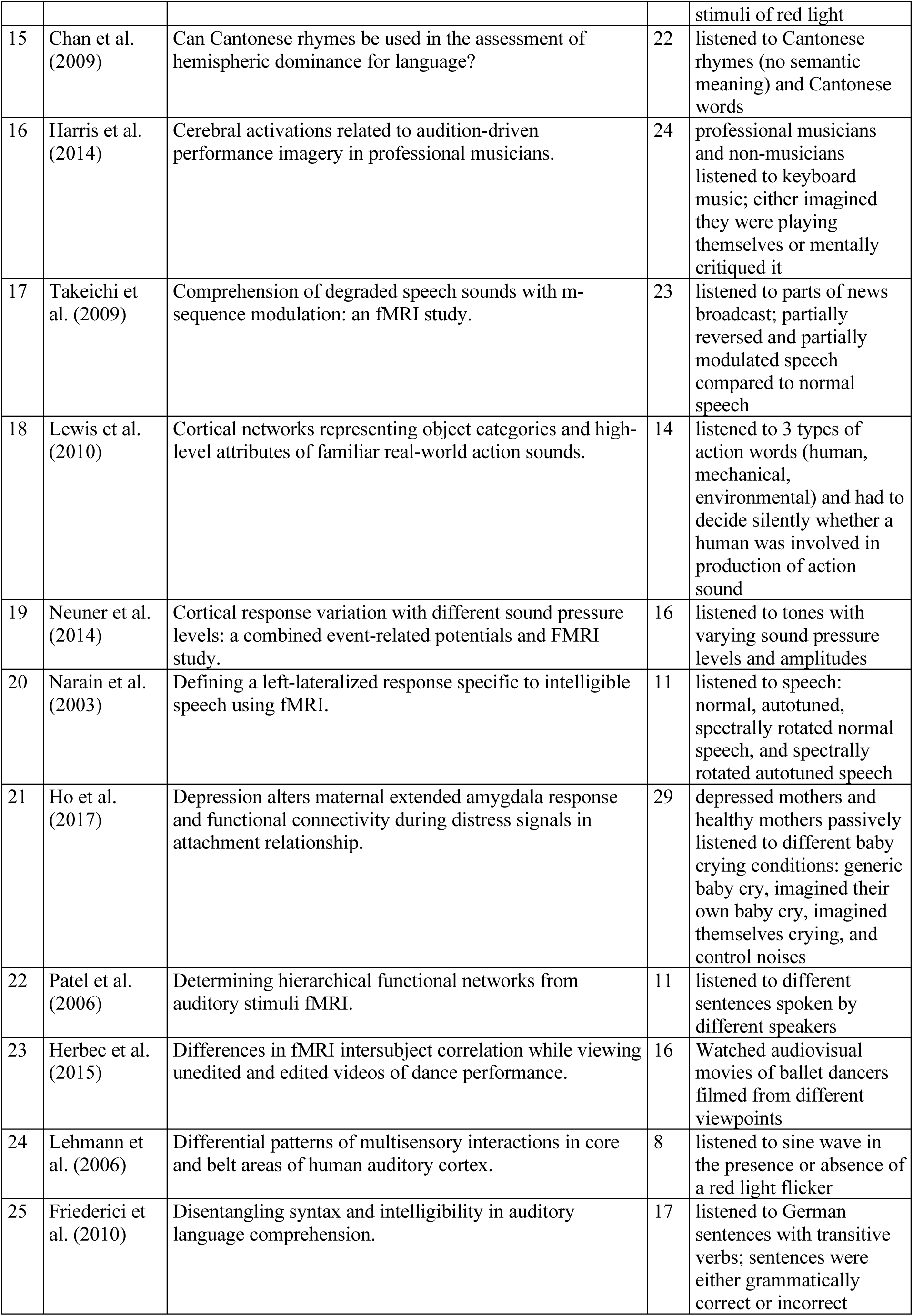
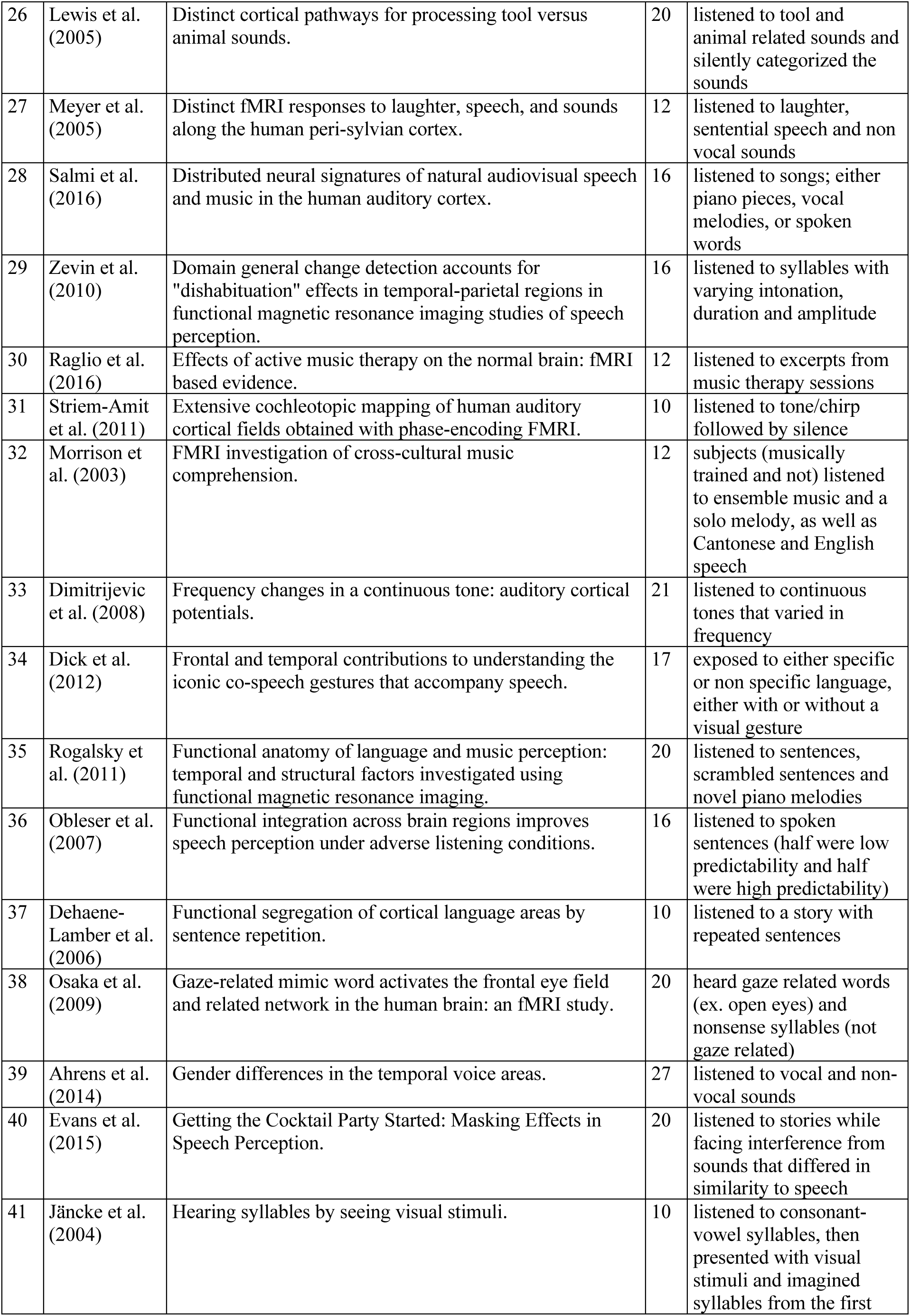
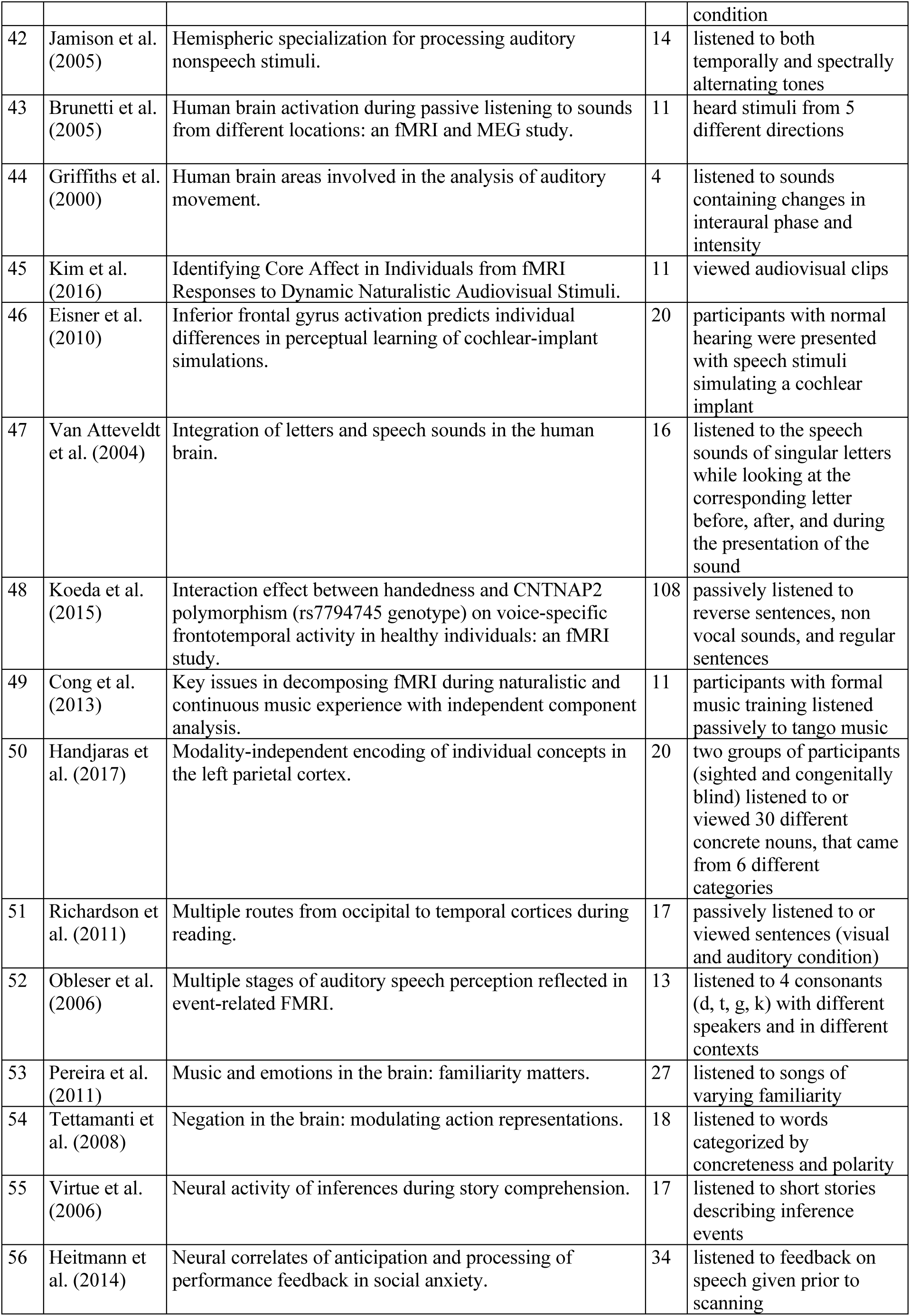
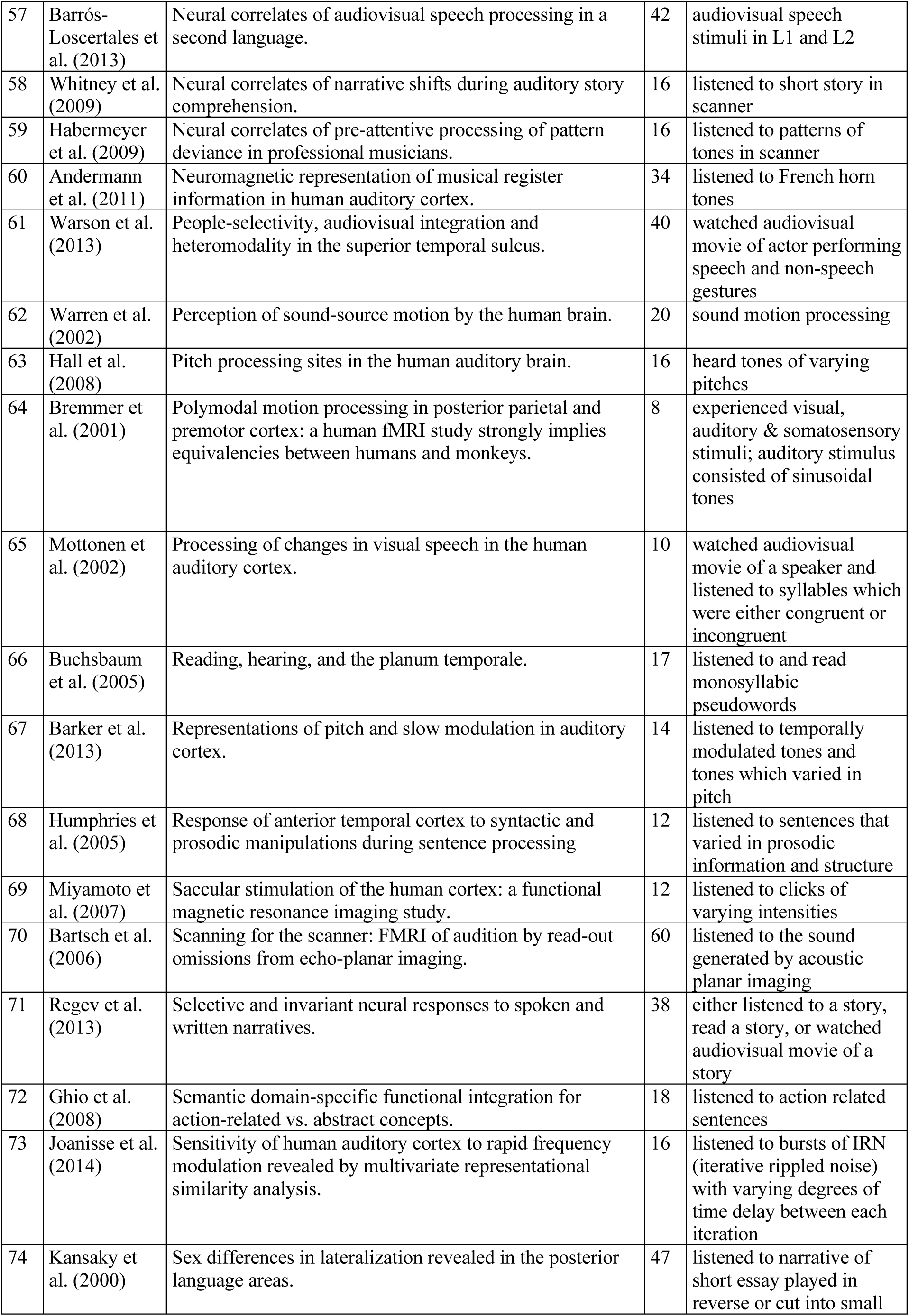
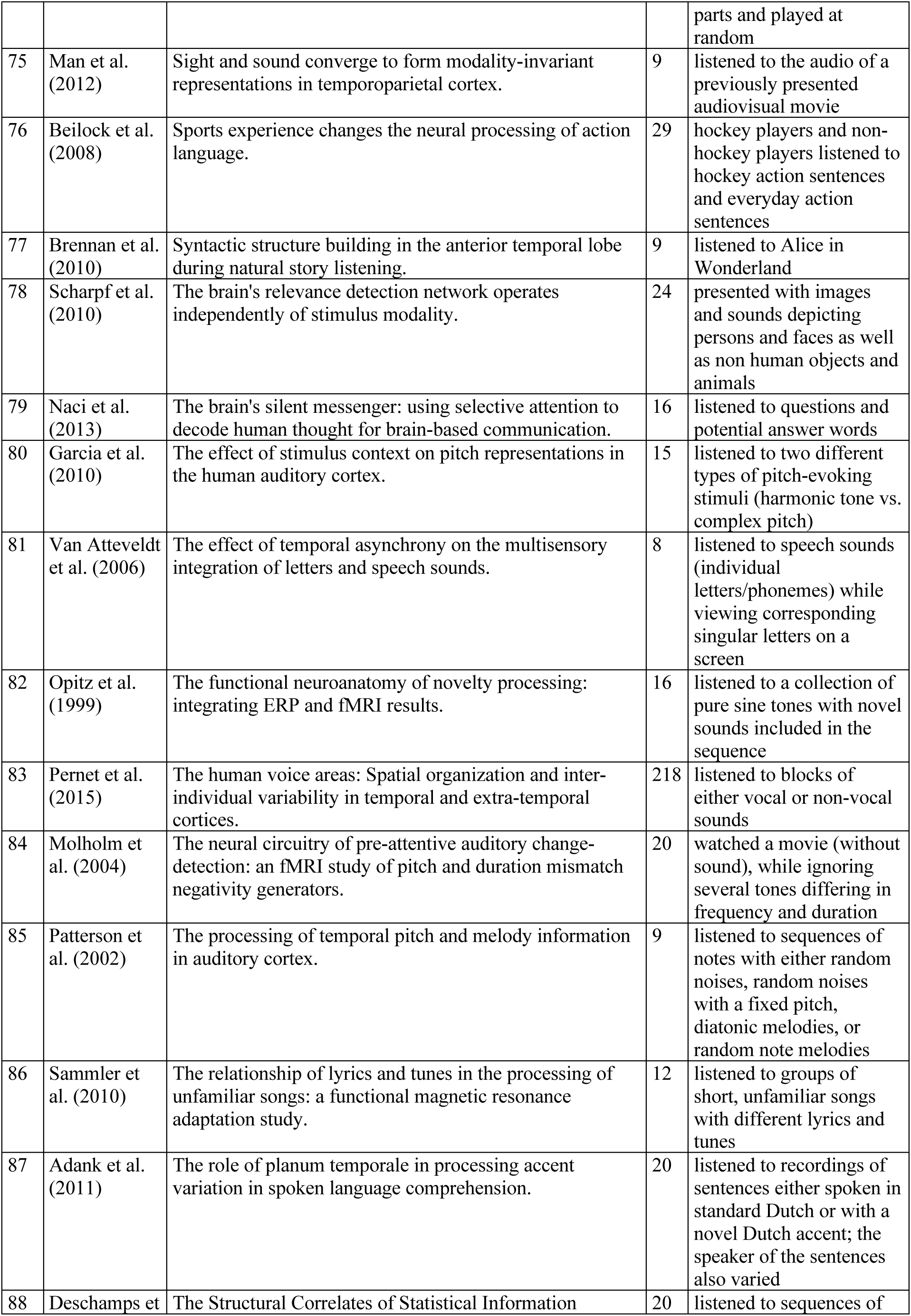
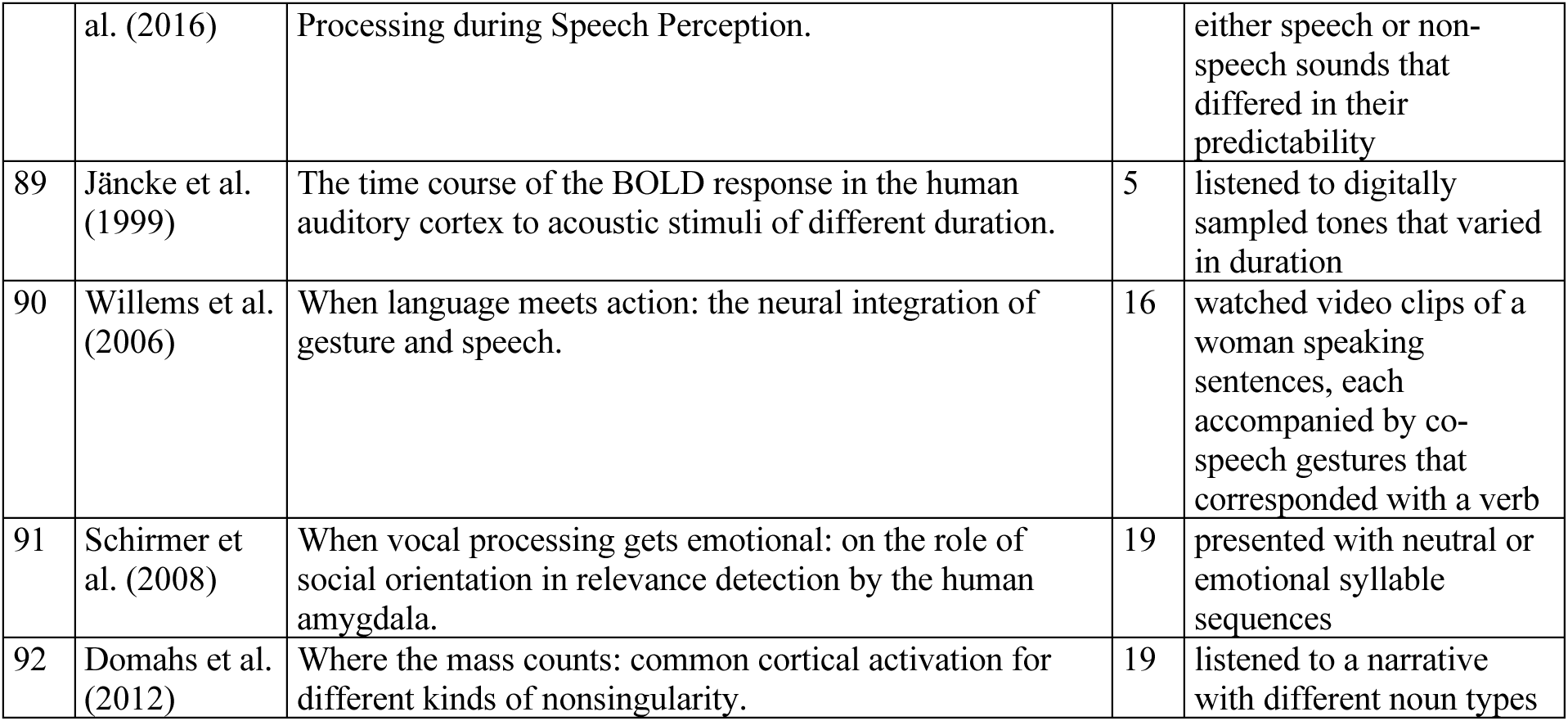
Studies with passive speech/sound perception without reported cerebellum activity.

**Table S4.** Cerebellar volume size in voxels and MNI x/y/z centre of mass (CM) coordinates for speech perception (red), production (blue) and overlap (yellow) regions for cerebellar meta-analyses (Figure 2) and coactivation meta-analysis (Figure 3). RL is right left; AP is anterior/posterior and IS is inferior/superior. Only clusters with a 10 or more voxels are shown for brevity.

| **Regions** | **Meta-Analysis** | **Volume** | **CM RL** | **CM AP** | **CM IS** |
| --- | --- | --- | --- | --- | --- |
| Overlap | Coactivation Meta-Analysis | 1895 | -49.8 | 6.0 | 30.8 |
| Overlap | Coactivation Meta-Analysis | 1269 | 49.2 | 10.7 | 29.4 |
| Overlap | Coactivation Meta-Analysis | 862 | 52.8 | 27.5 | 5.5 |
| Overlap | Coactivation Meta-Analysis | 674 | -20.7 | 9.5 | 2.0 |
| Overlap | Coactivation Meta-Analysis | 671 | 21.9 | 7.1 | 2.7 |
| Overlap | Coactivation Meta-Analysis | 479 | 15.7 | 69.0 | -3.8 |
| Overlap | Coactivation Meta-Analysis | 422 | -53.1 | 24.4 | 6.4 |
| Overlap | Coactivation Meta-Analysis | 391 | 3.5 | -1.7 | 52.5 |
| Overlap | Coactivation Meta-Analysis | 190 | -16.9 | 58.1 | -3.9 |
| Overlap | Coactivation Meta-Analysis | 176 | 35.0 | 21.7 | 53.0 |
| Overlap | Coactivation Meta-Analysis | 92 | 15.7 | 49.9 | 3.1 |
| Overlap | Coactivation Meta-Analysis | 78 | 55.2 | -15.5 | 11.9 |
| Overlap | Coactivation Meta-Analysis | 76 | 49.4 | -2.2 | 46.9 |
| Overlap | Coactivation Meta-Analysis | 61 | -44.4 | 29.7 | 31.4 |
| Overlap | Coactivation Meta-Analysis | 60 | -28.0 | 30.7 | 7.3 |
| Overlap | Coactivation Meta-Analysis | 57 | -11.5 | 42.5 | 56.0 |
| Overlap | Coactivation Meta-Analysis | 51 | -51.6 | -14.9 | -16.5 |
| Overlap | Coactivation Meta-Analysis | 50 | 36.8 | 12.9 | 16.8 |
| Overlap | Coactivation Meta-Analysis | 49 | 7.8 | 89.5 | 30.7 |
| Overlap | Coactivation Meta-Analysis | 44 | 52.0 | -13.6 | -15.1 |
| Overlap | Coactivation Meta-Analysis | 39 | 22.3 | -32.6 | -8.0 |
| Overlap | Coactivation Meta-Analysis | 39 | -20.9 | -4.5 | 22.8 |
| Overlap | Coactivation Meta-Analysis | 38 | -39.5 | -11.8 | -24.0 |
| Overlap | Coactivation Meta-Analysis | 37 | -39.2 | 19.5 | 5.1 |
| Overlap | Coactivation Meta-Analysis | 33 | 14.1 | 53.3 | 54.8 |
| Overlap | Coactivation Meta-Analysis | 32 | -25.3 | 60.4 | 8.8 |
| Overlap | Coactivation Meta-Analysis | 31 | 12.9 | -11.9 | -2.7 |
| Overlap | Coactivation Meta-Analysis | 30 | 25.4 | 32.1 | 3.9 |
| Overlap | Coactivation Meta-Analysis | 29 | 5.9 | 54.3 | 36.0 |
| Overlap | Coactivation Meta-Analysis | 28 | -8.2 | 85.7 | 25.6 |
| Overlap | Coactivation Meta-Analysis | 28 | 21.6 | 26.1 | 73.7 |
| Overlap | Coactivation Meta-Analysis | 27 | 13.6 | 27.6 | -36.9 |
| Overlap | Coactivation Meta-Analysis | 26 | 14.5 | 40.7 | -36.5 |
| Overlap | Coactivation Meta-Analysis | 26 | -65.4 | 9.8 | -0.5 |
| Overlap | Coactivation Meta-Analysis | 25 | 59.4 | -1.6 | -17.8 |
| Overlap | Coactivation Meta-Analysis | 25 | 35.8 | 29.1 | -13.4 |
| Overlap | Coactivation Meta-Analysis | 25 | -56.0 | 39.8 | 9.8 |
| Overlap | Coactivation Meta-Analysis | 22 | -59.0 | 8.2 | -11.8 |
| Overlap | Coactivation Meta-Analysis | 22 | 51.7 | -23.8 | -1.3 |
| Overlap | Coactivation Meta-Analysis | 22 | -11.7 | 95.0 | 17.4 |
| Overlap | Coactivation Meta-Analysis | 22 | -21.7 | 9.7 | 55.9 |
| Overlap | Coactivation Meta-Analysis | 21 | 17.9 | 100.5 | -7.6 |
| Overlap | Coactivation Meta-Analysis | 21 | 43.0 | 46.7 | 1.3 |
| Overlap | Coactivation Meta-Analysis | 21 | -25.1 | -28.5 | 5.6 |
| Overlap | Coactivation Meta-Analysis | 20 | 10.8 | 38.1 | 26.6 |
| Overlap | Coactivation Meta-Analysis | 20 | 14.2 | 10.1 | 44.8 |
| Overlap | Coactivation Meta-Analysis | 19 | -43.3 | 17.3 | -30.9 |
| Overlap | Coactivation Meta-Analysis | 19 | -49.2 | 21.3 | -20.1 |
| Overlap | Coactivation Meta-Analysis | 19 | 48.4 | 62.2 | 6.4 |
| Overlap | Coactivation Meta-Analysis | 19 | -38.1 | 36.9 | 64.0 |
| Overlap | Coactivation Meta-Analysis | 18 | -48.2 | 55.4 | -7.5 |
| Overlap | Coactivation Meta-Analysis | 18 | 21.9 | -2.7 | 30.3 |
| Overlap | Coactivation Meta-Analysis | 17 | -1.1 | 67.9 | 28.7 |
| Overlap | Coactivation Meta-Analysis | 17 | 20.1 | 25.9 | 57.3 |
| Overlap | Coactivation Meta-Analysis | 15 | -2.7 | 18.5 | -41.0 |
| Overlap | Coactivation Meta-Analysis | 15 | 13.9 | -19.1 | 3.8 |
| Overlap | Coactivation Meta-Analysis | 15 | -0.1 | 8.2 | 5.2 |
| Overlap | Coactivation Meta-Analysis | 15 | -30.6 | 55.5 | 6.7 |
| Overlap | Coactivation Meta-Analysis | 14 | -22.9 | 78.1 | 39.7 |
| Overlap | Coactivation Meta-Analysis | 14 | -27.3 | -30.0 | 54.7 |
| Overlap | Coactivation Meta-Analysis | 13 | -61.4 | 61.3 | 2.6 |
| Overlap | Coactivation Meta-Analysis | 12 | -11.0 | 70.2 | -6.5 |
| Overlap | Coactivation Meta-Analysis | 12 | 22.3 | 24.5 | 5.1 |
| Overlap | Coactivation Meta-Analysis | 11 | -57.7 | -5.7 | -16.7 |
| Overlap | Coactivation Meta-Analysis | 10 | -28.4 | 40.5 | 56.4 |
| Overlap | Meta-Analysis | 16 | 12.8 | 62.8 | -13.4 |
| Overlap | Meta-Analysis | 10 | -23.2 | 69.2 | -50.4 |
| Perception | Coactivation Meta-Analysis | 3518 | 54.0 | 6.8 | -10.0 |
| Perception | Coactivation Meta-Analysis | 1380 | -52.4 | -1.5 | -23.0 |
| Perception | Coactivation Meta-Analysis | 1117 | 5.5 | -48.7 | 38.6 |
| Perception | Coactivation Meta-Analysis | 875 | 46.9 | 63.1 | 28.9 |
| Perception | Coactivation Meta-Analysis | 516 | -0.6 | 52.8 | 33.3 |
| Perception | Coactivation Meta-Analysis | 465 | -50.5 | 61.2 | 27.7 |
| Perception | Coactivation Meta-Analysis | 428 | 38.4 | -13.8 | 46.5 |
| Perception | Coactivation Meta-Analysis | 308 | -53.3 | -28.6 | -1.3 |
| Perception | Coactivation Meta-Analysis | 255 | 1.2 | -48.3 | -14.4 |
| Perception | Coactivation Meta-Analysis | 238 | 6.5 | 24.5 | -38.7 |
| Perception | Coactivation Meta-Analysis | 217 | -23.4 | -29.1 | 47.5 |
| Perception | Coactivation Meta-Analysis | 169 | 16.8 | -8.5 | 11.5 |
| Perception | Coactivation Meta-Analysis | 147 | 27.8 | 32.5 | -16.6 |
| Perception | Coactivation Meta-Analysis | 94 | -47.4 | 29.9 | -2.5 |
| Perception | Coactivation Meta-Analysis | 90 | 24.0 | -42.7 | 23.5 |
| Perception | Coactivation Meta-Analysis | 73 | 27.3 | -15.2 | -32.7 |
| Perception | Coactivation Meta-Analysis | 71 | -16.0 | 41.5 | 75.4 |
| Perception | Coactivation Meta-Analysis | 61 | -23.1 | 32.5 | -16.2 |
| Perception | Coactivation Meta-Analysis | 59 | -50.7 | -38.9 | 12.3 |
| Perception | Coactivation Meta-Analysis | 58 | 13.3 | 62.9 | -5.4 |
| Perception | Coactivation Meta-Analysis | 48 | 23.6 | 99.2 | -11.7 |
| Perception | Coactivation Meta-Analysis | 45 | 17.9 | 12.5 | -31.7 |
| Perception | Coactivation Meta-Analysis | 43 | -29.3 | 6.8 | -42.8 |
| Perception | Coactivation Meta-Analysis | 43 | -15.4 | -13.4 | 15.2 |
| Perception | Coactivation Meta-Analysis | 40 | -10.4 | 58.0 | 1.0 |
| Perception | Coactivation Meta-Analysis | 36 | -41.8 | 74.5 | 26.5 |
| Perception | Coactivation Meta-Analysis | 35 | -6.3 | 21.1 | 66.0 |
| Perception | Coactivation Meta-Analysis | 29 | 7.4 | -32.6 | -21.9 |
| Perception | Coactivation Meta-Analysis | 29 | 15.7 | -40.9 | -6.1 |
| Perception | Coactivation Meta-Analysis | 24 | 35.7 | 12.2 | 19.5 |
| Perception | Coactivation Meta-Analysis | 24 | 24.3 | -39.2 | 47.7 |
| Perception | Coactivation Meta-Analysis | 22 | -2.7 | -33.6 | -5.3 |
| Perception | Coactivation Meta-Analysis | 19 | 34.5 | -16.8 | -43.1 |
| Perception | Coactivation Meta-Analysis | 18 | 27.9 | -34.7 | -13.6 |
| Perception | Coactivation Meta-Analysis | 18 | -1.0 | 33.9 | 33.4 |
| Perception | Coactivation Meta-Analysis | 17 | 22.7 | 10.0 | 32.1 |
| Perception | Coactivation Meta-Analysis | 17 | -7.8 | -18.7 | 65.2 |
| Perception | Coactivation Meta-Analysis | 16 | 13.2 | 40.1 | -38.8 |
| Perception | Coactivation Meta-Analysis | 16 | -33.0 | 30.9 | 4.7 |
| Perception | Coactivation Meta-Analysis | 15 | -6.5 | 39.7 | -47.7 |
| Perception | Coactivation Meta-Analysis | 14 | 26.6 | 3.5 | -25.7 |
| Perception | Coactivation Meta-Analysis | 14 | 13.1 | 46.6 | 5.1 |
| Perception | Coactivation Meta-Analysis | 13 | -65.9 | 46.5 | -7.7 |
| Perception | Coactivation Meta-Analysis | 13 | 7.2 | 14.0 | 8.6 |
| Perception | Coactivation Meta-Analysis | 12 | 31.7 | 1.3 | -33.8 |
| Perception | Coactivation Meta-Analysis | 12 | -0.9 | 41.0 | 54.0 |
| Perception | Coactivation Meta-Analysis | 11 | -3.0 | 31.3 | -54.7 |
| Perception | Coactivation Meta-Analysis | 10 | 18.8 | 23.2 | -23.0 |
| Perception | Coactivation Meta-Analysis | 10 | 4.1 | 44.9 | 9.0 |
| Perception | Coactivation Meta-Analysis | 10 | -42.8 | 70.3 | 40.2 |
| Perception | Coactivation Meta-Analysis | 10 | 9.8 | 30.4 | 70.2 |
| Perception | Meta-Analysis | 200 | -20.3 | 78.9 | -41.7 |
| Perception | Meta-Analysis | 46 | -26.3 | 75.2 | -32.7 |
| Perception | Meta-Analysis | 43 | -42.3 | 54.6 | -56.1 |
| Perception | Meta-Analysis | 29 | 12.2 | 40.0 | -55.9 |
| Perception | Meta-Analysis | 28 | 11.2 | 62.3 | -9.3 |
| Perception | Meta-Analysis | 23 | 20.6 | 77.3 | -35.7 |
| Perception | Meta-Analysis | 14 | 31.3 | 64.4 | -59.7 |
| Perception | Meta-Analysis | 13 | 20.9 | 33.2 | -26.6 |
| Production | Coactivation Meta-Analysis | 2116 | 42.3 | 18.5 | 44.7 |
| Production | Coactivation Meta-Analysis | 1143 | 0.3 | -3.3 | 53.5 |
| Production | Coactivation Meta-Analysis | 679 | -38.7 | 43.2 | 48.8 |
| Production | Coactivation Meta-Analysis | 522 | -34.1 | 6.2 | 55.1 |
| Production | Coactivation Meta-Analysis | 402 | 16.4 | 11.3 | 5.4 |
| Production | Coactivation Meta-Analysis | 275 | -54.7 | -5.7 | 35.2 |
| Production | Coactivation Meta-Analysis | 242 | -36.6 | 63.4 | -15.8 |
| Production | Coactivation Meta-Analysis | 222 | -13.4 | 16.2 | 7.0 |
| Production | Coactivation Meta-Analysis | 135 | -49.0 | -9.1 | 3.8 |
| Production | Coactivation Meta-Analysis | 123 | -36.0 | -20.2 | 4.2 |
| Production | Coactivation Meta-Analysis | 99 | 34.0 | 62.1 | -16.2 |
| Production | Coactivation Meta-Analysis | 93 | -59.8 | 27.2 | 21.1 |
| Production | Coactivation Meta-Analysis | 77 | -20.3 | 64.1 | 55.7 |
| Production | Coactivation Meta-Analysis | 70 | -22.1 | -3.1 | 6.2 |
| Production | Coactivation Meta-Analysis | 70 | 57.1 | 17.8 | 18.7 |
| Production | Coactivation Meta-Analysis | 70 | -36.3 | -37.6 | 26.9 |
| Production | Coactivation Meta-Analysis | 66 | 32.3 | -18.0 | 2.7 |
| Production | Coactivation Meta-Analysis | 44 | -31.9 | 86.9 | 0.2 |
| Production | Coactivation Meta-Analysis | 37 | 48.2 | 37.5 | 23.9 |
| Production | Coactivation Meta-Analysis | 23 | -8.4 | 75.5 | -10.3 |
| Production | Coactivation Meta-Analysis | 22 | -16.9 | 1.6 | -2.1 |
| Production | Coactivation Meta-Analysis | 13 | -43.8 | 64.8 | 4.0 |
| Production | Coactivation Meta-Analysis | 12 | 14.1 | 67.5 | 59.4 |
| Production | Meta-Analysis | 1929 | -9.7 | 62.9 | -22.4 |
| Production | Meta-Analysis | 184 | -26.2 | 63.2 | -53.7 |
| Production | Meta-Analysis | 118 | 40.7 | 64.6 | -31.5 |
| Production | Meta-Analysis | 82 | -12.2 | 77.7 | -44.5 |
| Production | Meta-Analysis | 48 | -14.0 | 39.5 | -46.8 |
| Production | Meta-Analysis | 25 | -5.8 | 58.9 | -31.0 |
| Production | Meta-Analysis | 21 | 7.8 | 45.0 | -17.6 |
| Production | Meta-Analysis | 20 | -25.2 | 47.0 | -46.3 |
| Production | Meta-Analysis | 16 | -11.5 | 67.9 | -38.8 |
| Production | Meta-Analysis | 15 | 10.8 | 53.9 | -32.9 |
| Production | Meta-Analysis | 13 | -0.5 | 57.4 | -0.6 |
| Production | Meta-Analysis | 12 | 29.0 | 31.3 | -27.0 |
| Production | Meta-Analysis | 10 | 1.0 | 45.4 | -7.4 |

**Table S5.**
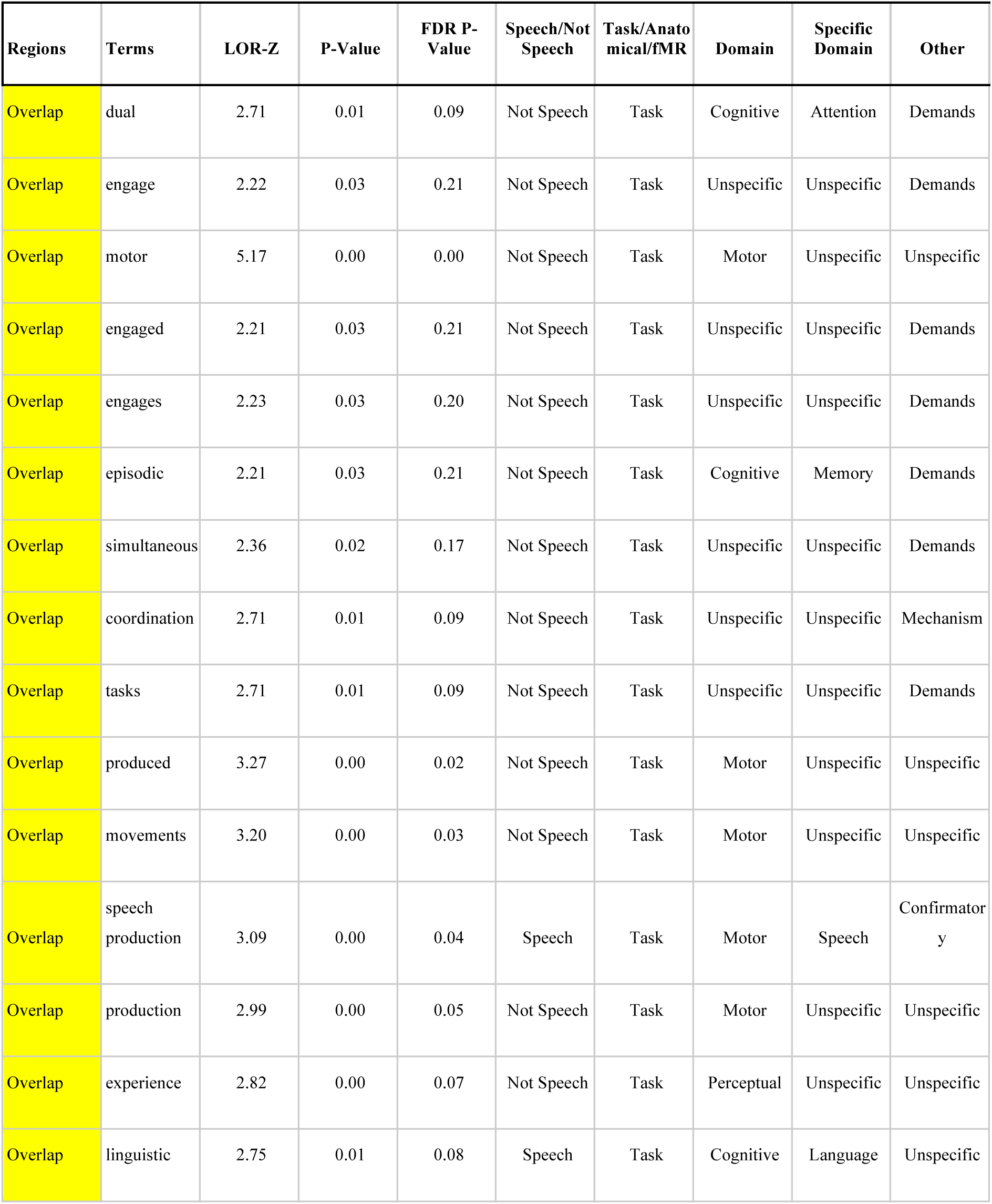
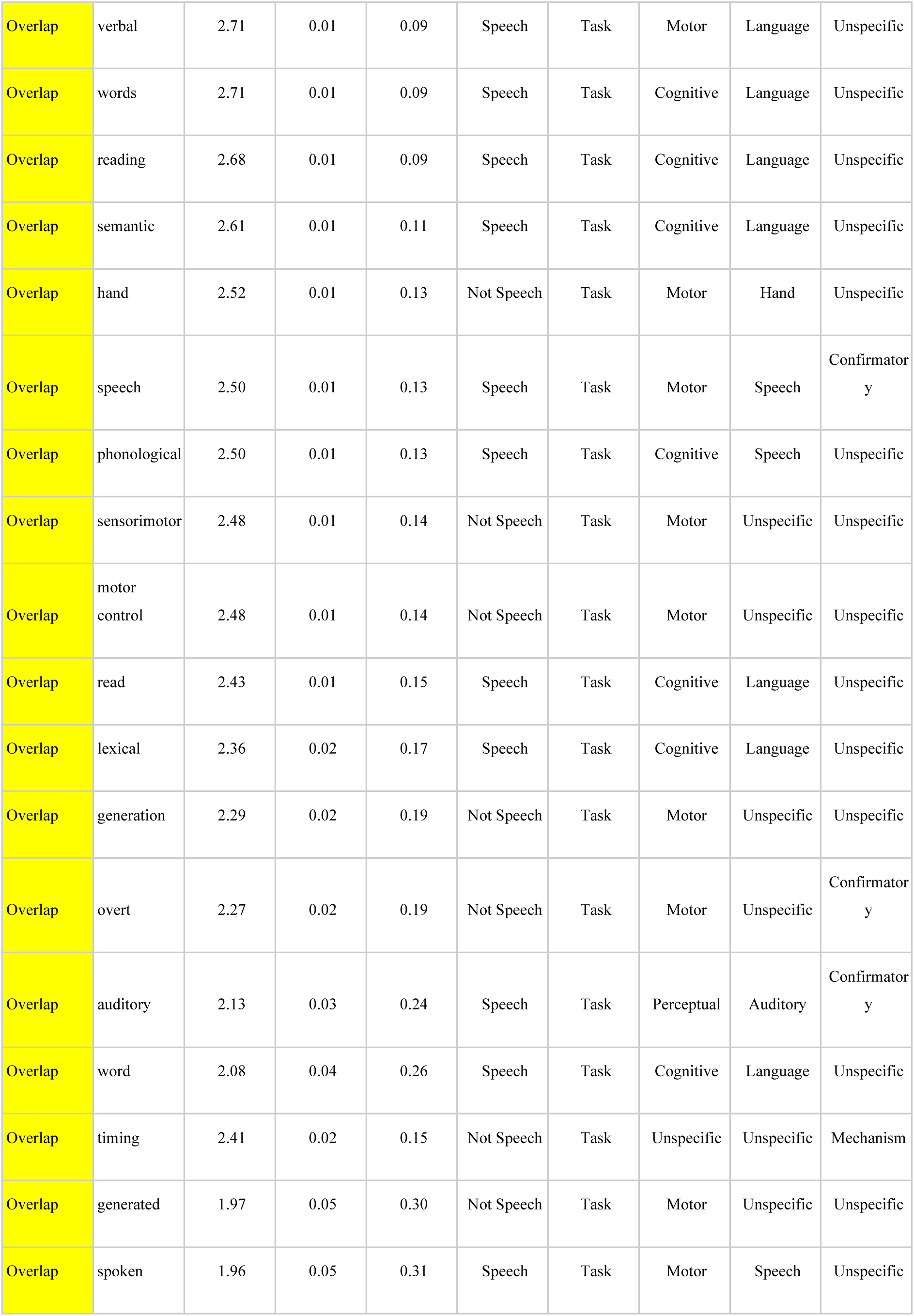
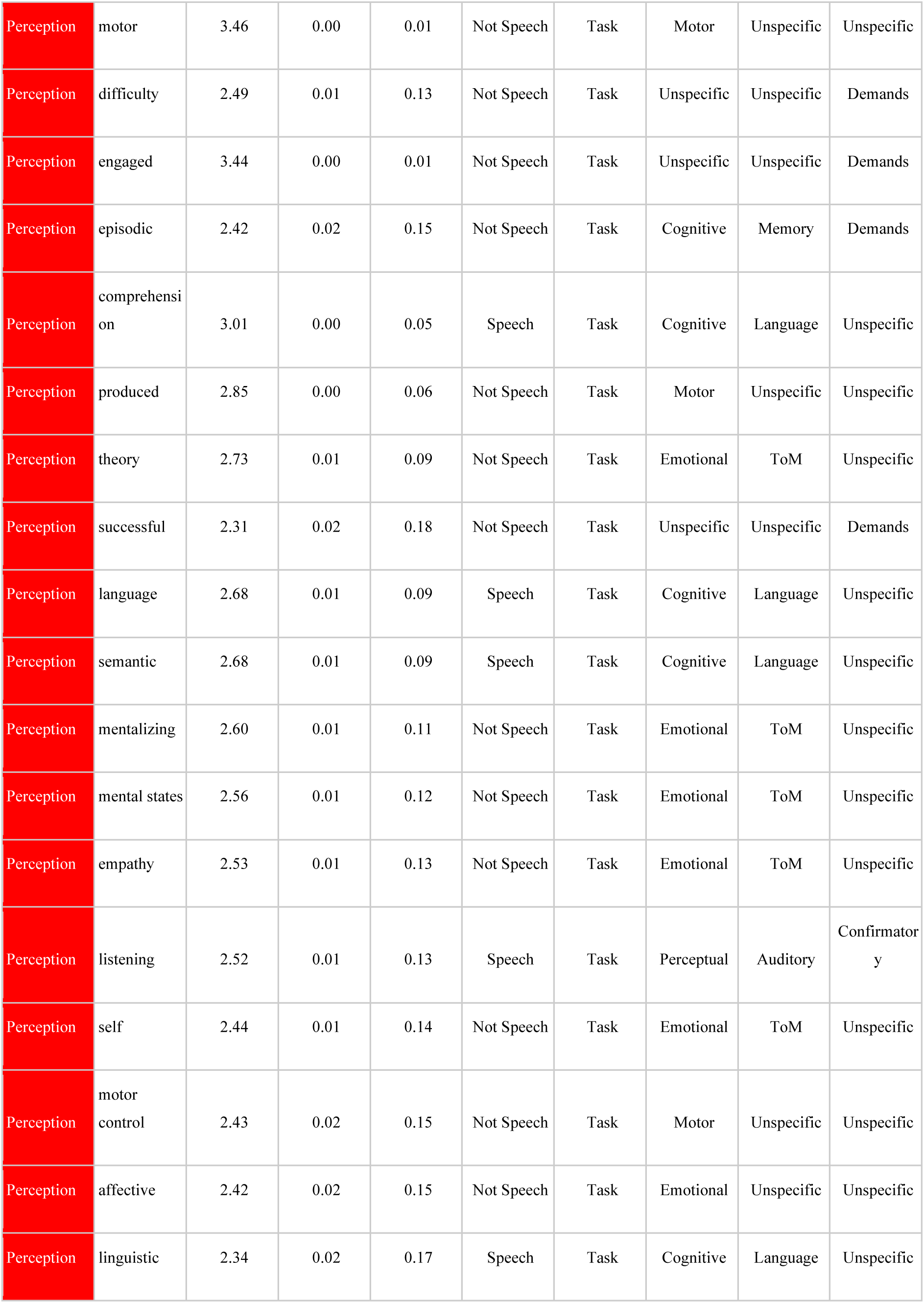
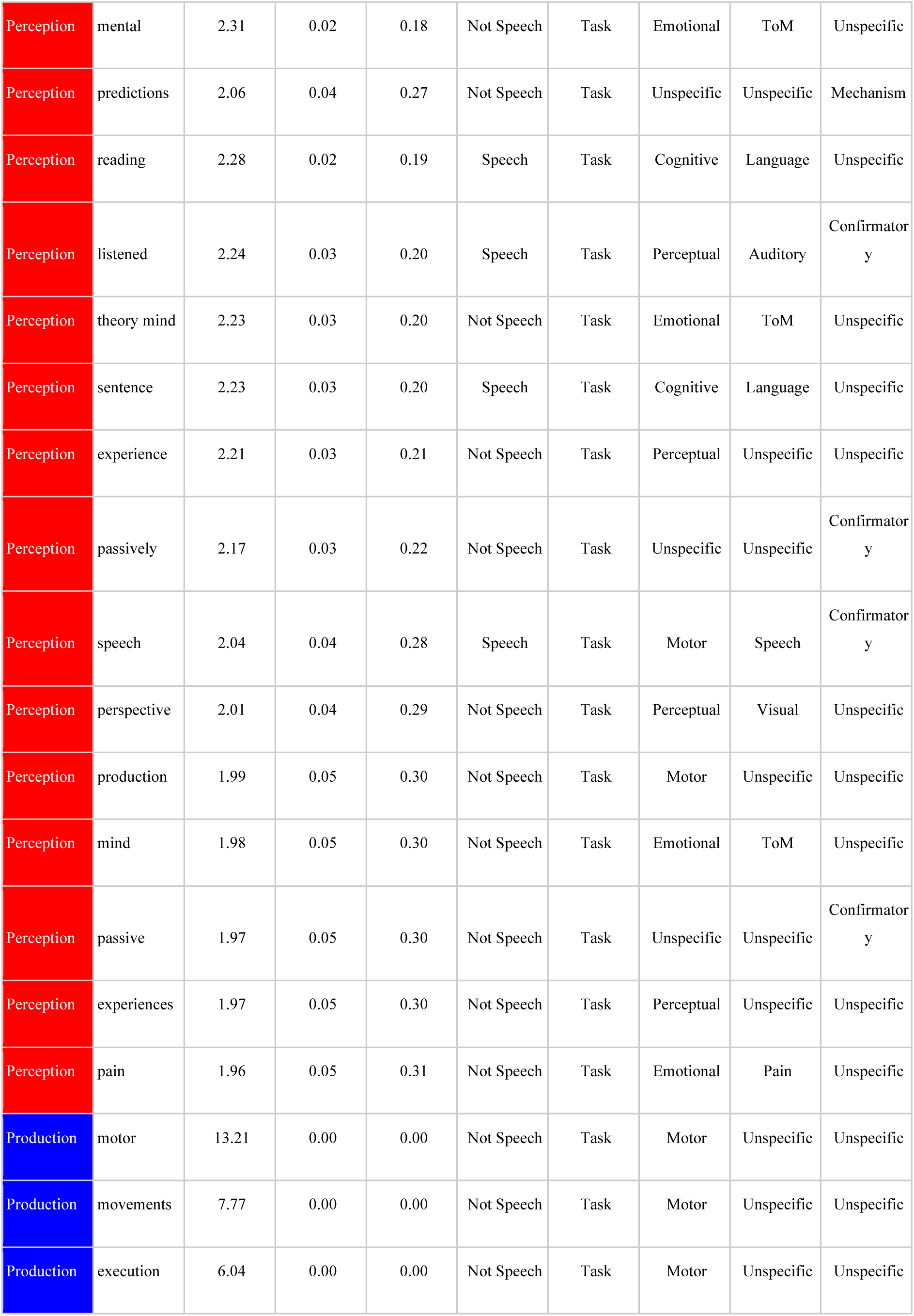
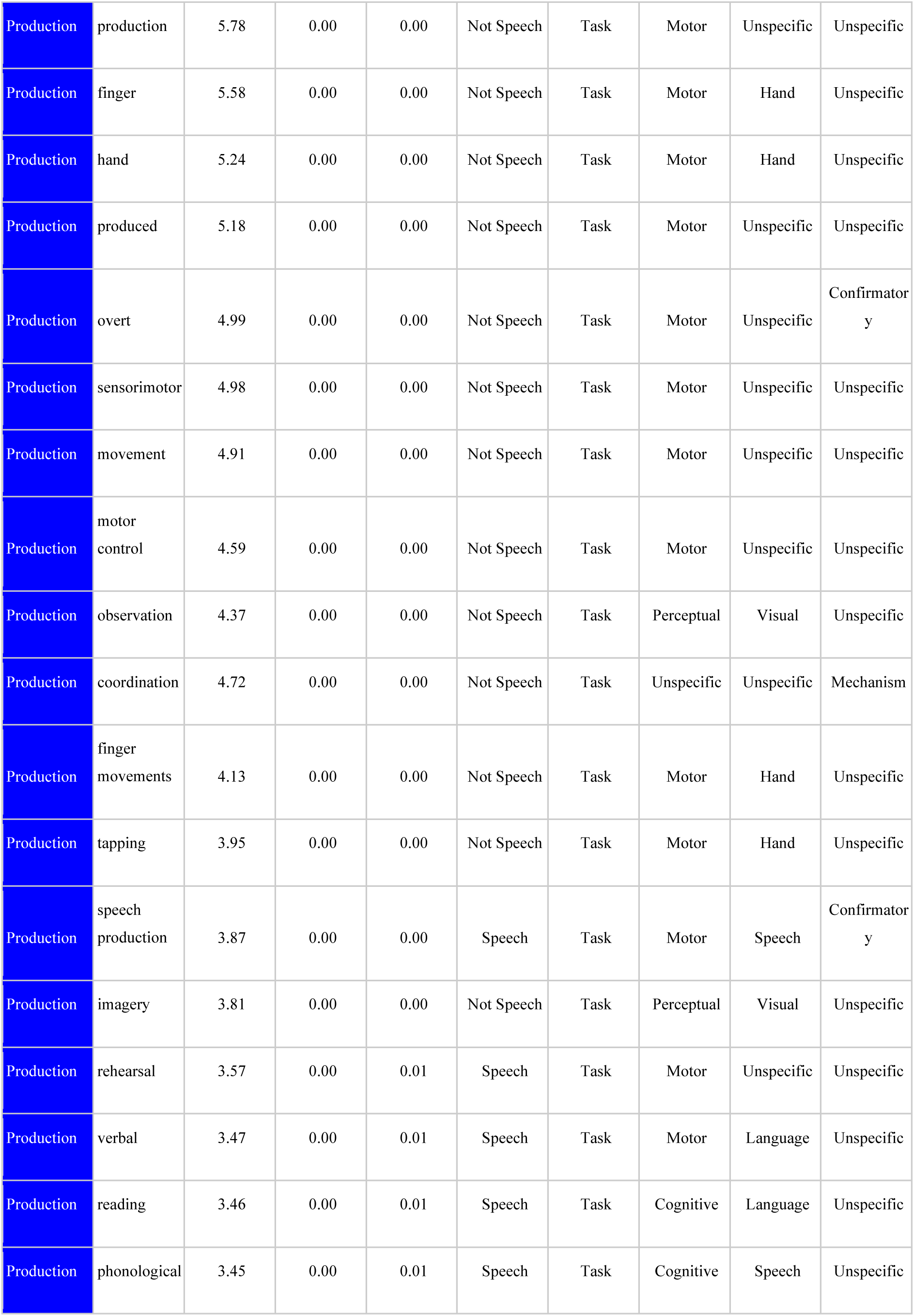
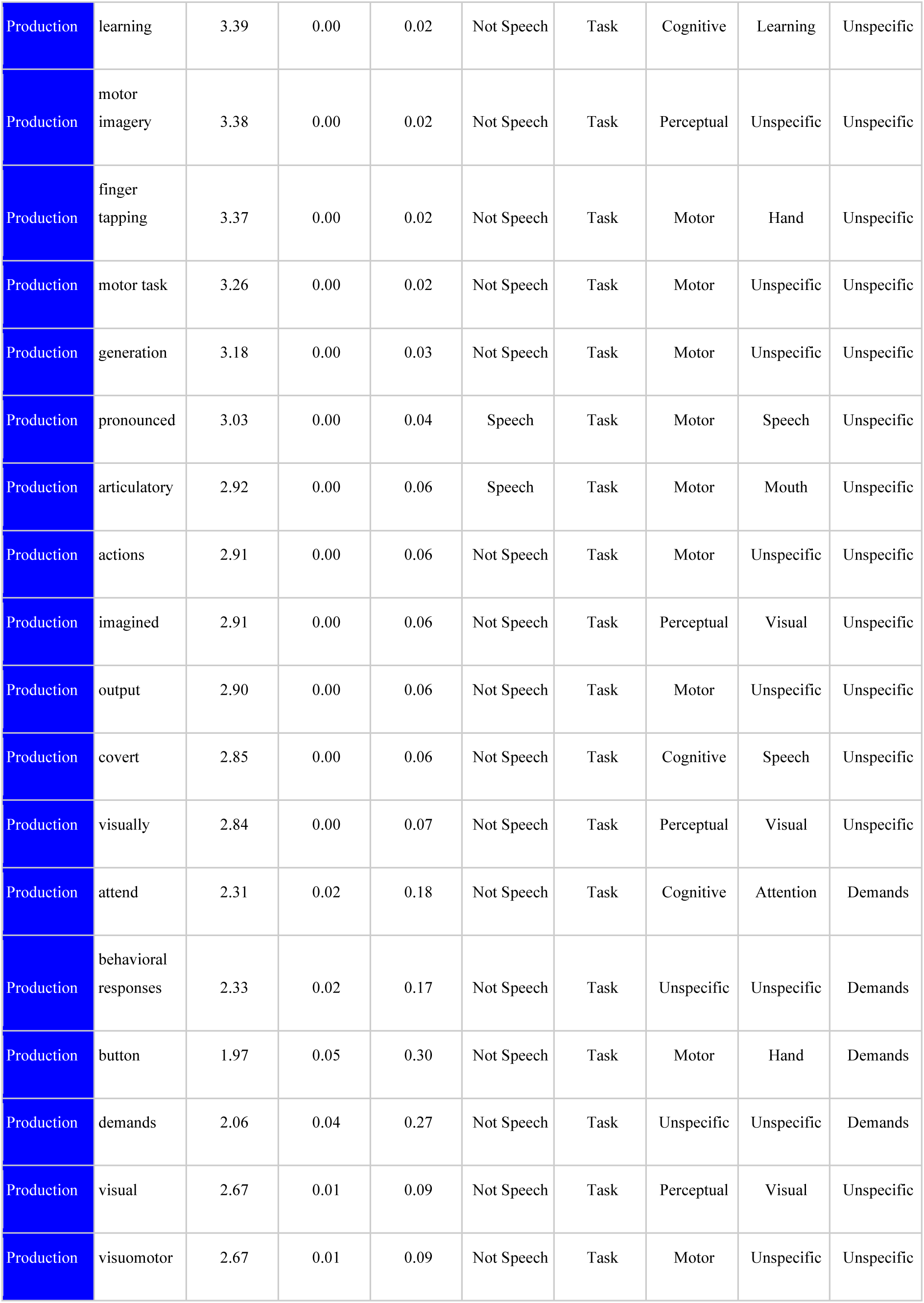
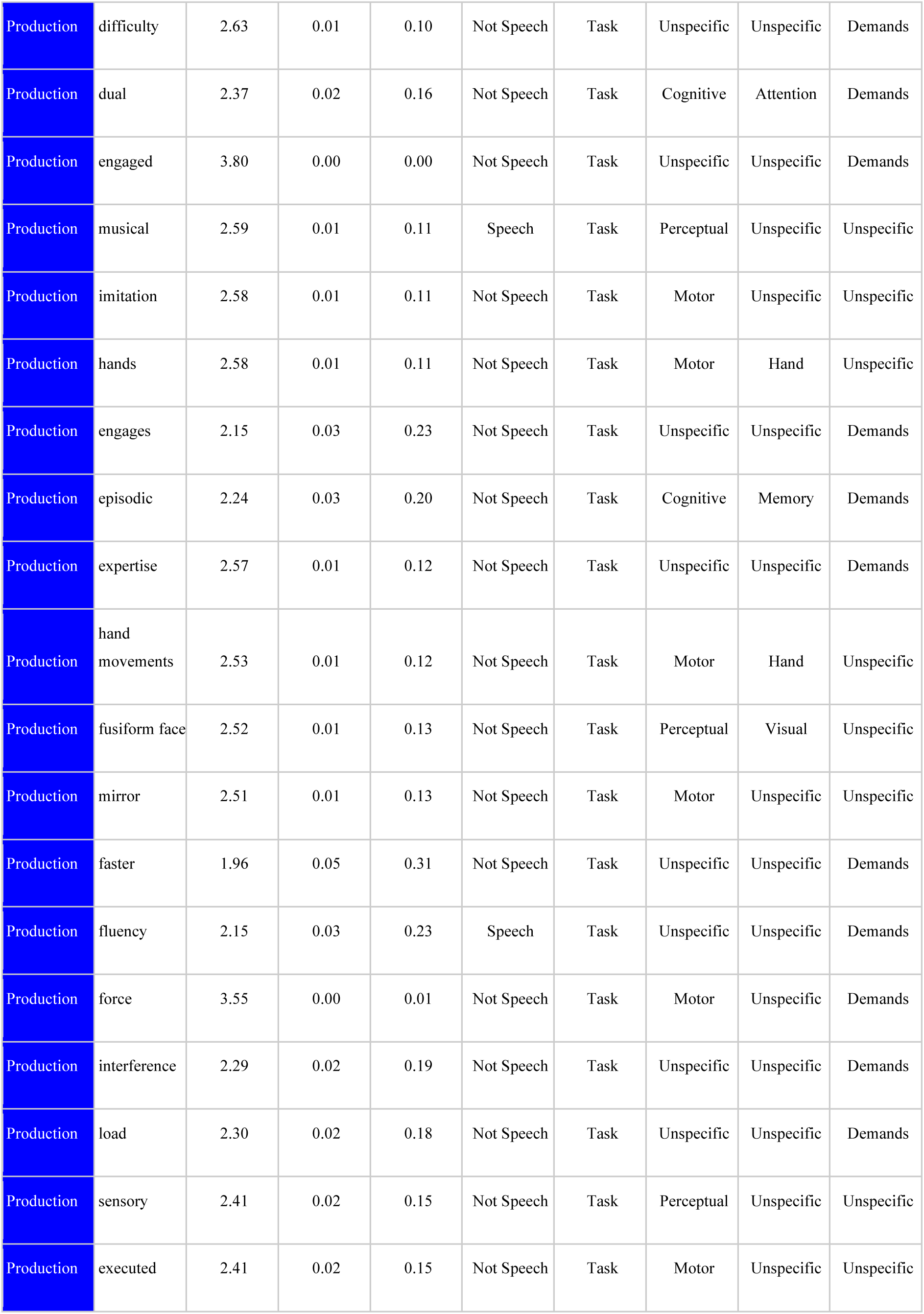
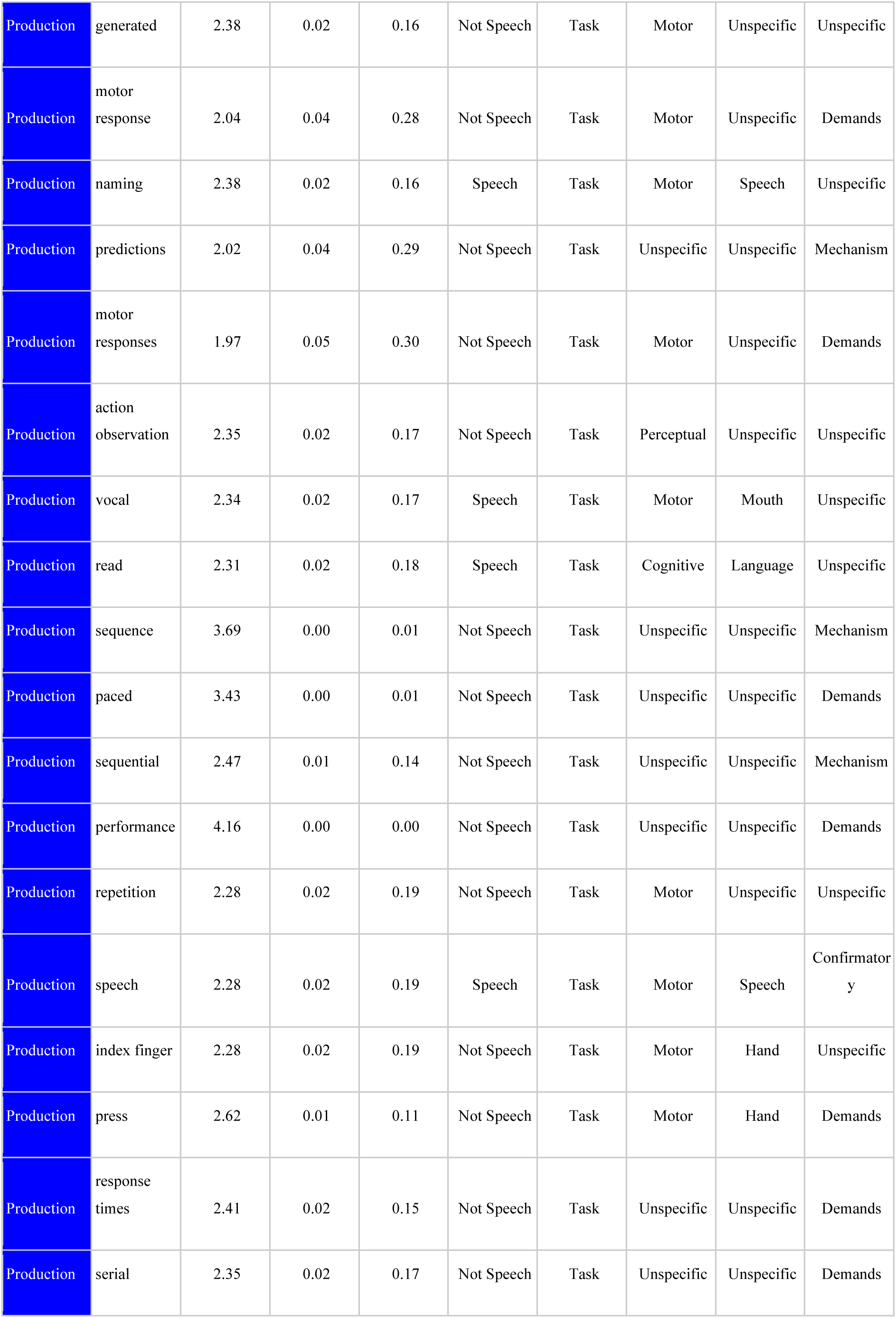
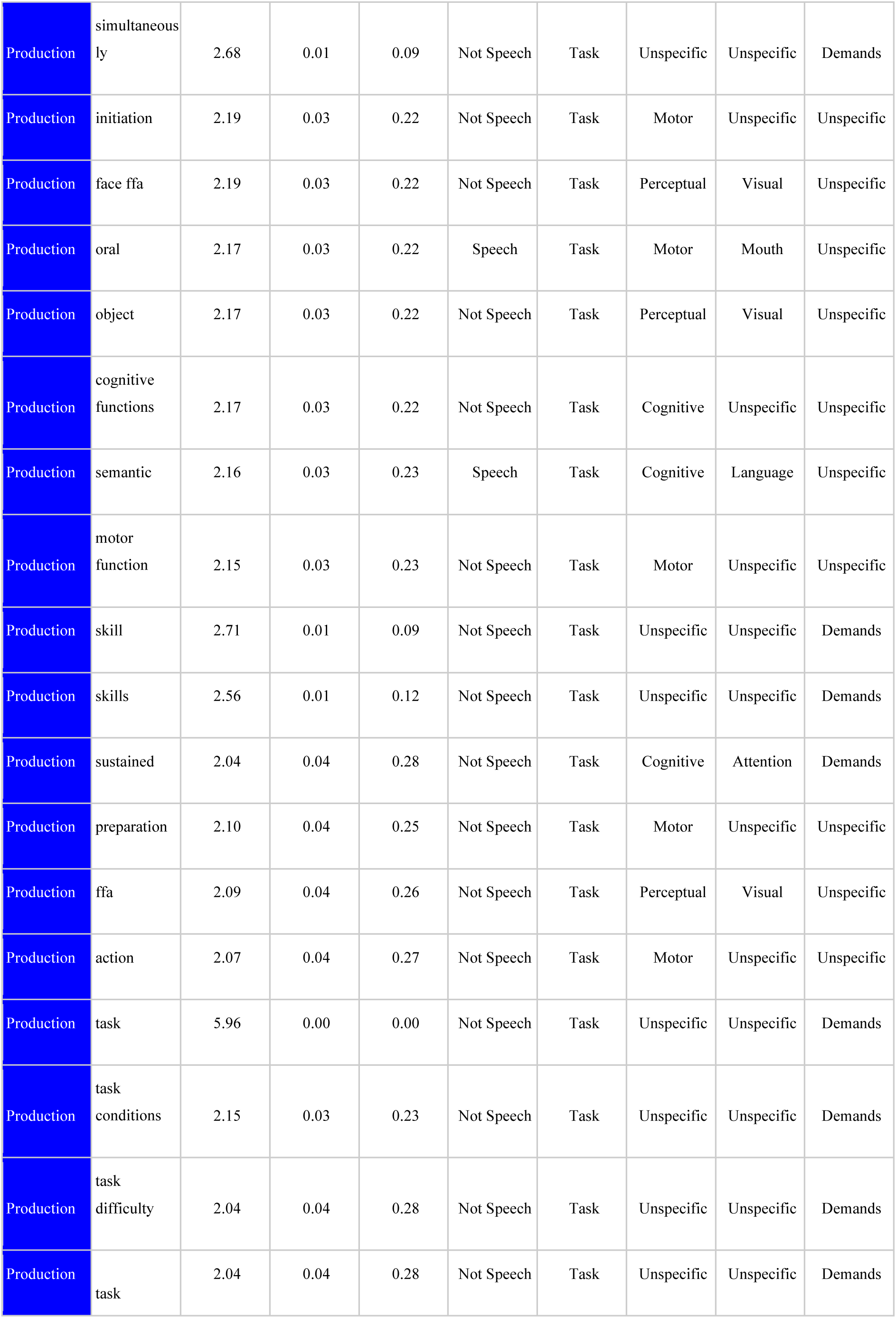
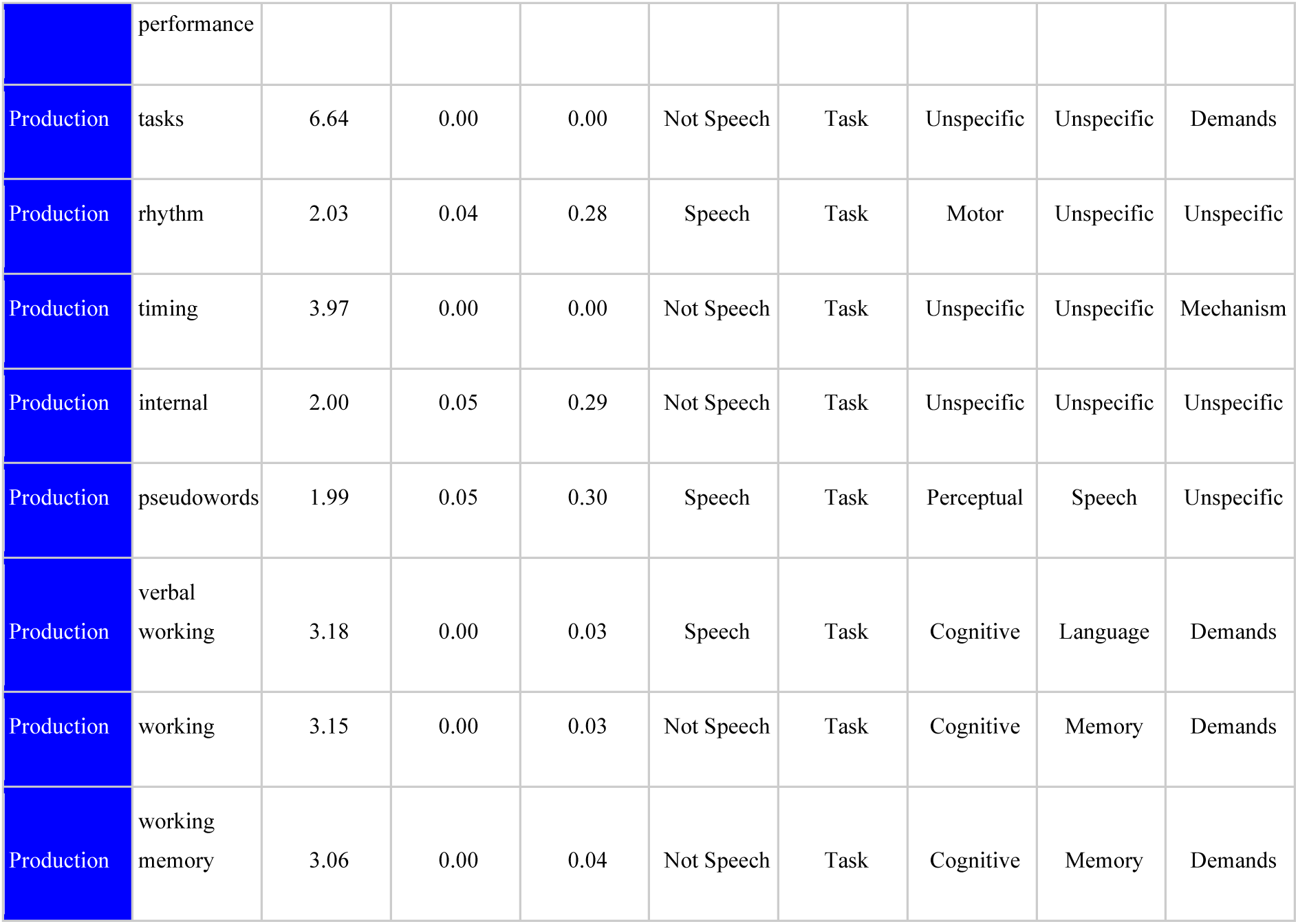
All terms associated with each set of regions in the cerebellum were categorised as being speech related or not, into five gross domains of psychological inquiry and other categorisations. These are summarised in Table 2.

## References

Ackermann, H. (2008). Cerebellar contributions to speech production and speech perception: psycholinguistic and neurobiological perspectives. Trends in Neurosciences, 31(6), 265–272. https://doi.org/10.1016/j.tins.2008.02.011

Ackermann, H., & Brendel, B. (2016). Chapter 7 - Cerebellar Contributions to Speech and Language. In G. Hickok & S. L. Small (Eds.), Neurobiology of Language (pp. 73–84). Academic Press. https://doi.org/10.1016/B978-0-12-407794-2.00007-9

Ackermann, H., Gräber, S., Hertrich, I., & Daum, I. (1997). Categorical speech perception in cerebellar disorders. Brain and Language, 60(2), 323–331. https://doi.org/10.1006/brln.1997.1826

Ackermann, H., Mathiak, K., & Riecker, A. (2007). The contribution of the cerebellum to speech production and speech perception: clinical and functional imaging data. Cerebellum, 6(3), 202–213. https://doi.org/10.1080/14734220701266742

Ackermann, H., Wildgruber, D., Daum, I., & Grodd, W. (1998). Does the cerebellum contribute to cognitive aspects of speech production? A functional magnetic resonance imaging (fMRI) study in humans. Neuroscience Letters, 247(2-3), 187–190. https://doi.org/10.1016/s0304-3940(98)00328-0

Argyropoulos, G. P. D. (2016). The cerebellum, internal models and prediction in “non-motor” aspects of language: A critical review. Brain and Language, 161, 4–17. https://doi.org/10.1016/j.bandl.2015.08.003

Ashida, R., Cerminara, N. L., Edwards, R. J., Apps, R., & Brooks, J. C. W. (2019). Sensorimotor, language, and working memory representation within the human cerebellum. Human Brain Mapping, 40(16), 4732–4747. https://doi.org/10.1002/hbm.24733

Baizer, J. S., & Glickstein, M. (1974). Proceedings: Role of cerebellum in prism adaptation. The Journal of Physiology, 236(1), 34P – 35P. https://www.ncbi.nlm.nih.gov/pubmed/4206505

Balsters, J. H., Laird, A. R., Fox, P. T., & Eickhoff, S. B. (2014). Bridging the gap between functional and anatomical features of cortico-cerebellar circuits using meta-analytic connectivity modeling. Human Brain Mapping, 35(7), 3152–3169. https://doi.org/10.1002/hbm.22392

Barton, R. A., & Venditti, C. (2017). Rapid Evolution of the Cerebellum in Humans and Other Great Apes. Current Biology: CB, 27(8), 1249–1250. https://doi.org/10.1016/j.cub.2017.03.059

Boillat, Y., Bazin, P.-L., & van der Zwaag, W. (2020). Whole-body somatotopic maps in the cerebellum revealed with 7T fMRI. NeuroImage, 116624. https://doi.org/10.1016/j.neuroimage.2020.116624

Bonhage, C. E., Mueller, J. L., Friederici, A. D., & Fiebach, C. J. (2015). Combined eye tracking and fMRI reveals neural basis of linguistic predictions during sentence comprehension. Cortex; a Journal Devoted to the Study of the Nervous System and Behavior, 68, 33–47. https://doi.org/10.1016/j.cortex.2015.04.011

Buckner, R. L., Krienen, F. M., Castellanos, A., Diaz, J. C., & Yeo, B. T. T. (2011). The organization of the human cerebellum estimated by intrinsic functional connectivity. Journal of Neurophysiology, 106(5), 2322–2345. https://doi.org/10.1152/jn.00339.2011

Bybee, J., & McClelland, J. L. (2005). Alternatives to the combinatorial paradigm of linguistic theory based on domain general principles of human cognition. The Linguistic Review, 22(2-4). https://doi.org/10.1515/tlir.2005.22.2-4.381

Callan, D. E., Kawato, M., Parsons, L., & Turner, R. (2007). Speech and song: the role of the cerebellum. Cerebellum, 6(4), 321–327. https://doi.org/10.1080/14734220601187733

Chen, S. H. A., & Desmond, J. E. (2005a). Cerebrocerebellar networks during articulatory rehearsal and verbal working memory tasks. NeuroImage, 24(2), 332–338. https://doi.org/10.1016/j.neuroimage.2004.08.032

Chen, S. H. A., & Desmond, J. E. (2005b). Temporal dynamics of cerebro-cerebellar network recruitment during a cognitive task. Neuropsychologia, 43(9), 1227–1237. https://doi.org/10.1016/j.neuropsychologia.2004.12.015

Clark, A. (2013). Whatever next? Predictive brains, situated agents, and the future of cognitive science. The Behavioral and Brain Sciences, 36(3), 181–204. https://doi.org/10.1017/S0140525X12000477

de la Vega, A., Chang, L.J., Banich, M.T., Wager, T.D., Yarkoni, T. (2016) Large-Scale Meta-Analysis of Human Medial Frontal Cortex Reveals Tripartite Functional Organization. The Journal of Neuroscience: The Official Journal of the Society for Neuroscience, 36(24):6553–6562. https://doi.org/10.1523/JNEUROSCI.4402-15.2016

de la Vega, A., Yarkoni, T., Wager, T.D., Banich, M.T., (2018) Large-scale Meta-analysis Suggests Low Regional Modularity in Lateral Frontal Cortex. Cerebral Cortex. 28(10):3414–3428. https://doi.org/10.1093/cercor/bhx204

Diedrichsen, J., Balsters, J. H., Flavell, J., Cussans, E., & Ramnani, N. (2009). A probabilistic MR atlas of the human cerebellum. NeuroImage, 46(1), 39–46. https://doi.org/10.1016/j.neuroimage.2009.01.045

Diedrichsen, J., King, M., Hernandez-Castillo, C., Sereno, M., & Ivry, R. B. (2019). Universal Transform or Multiple Functionality? Understanding the Contribution of the Human Cerebellum across Task Domains. Neuron, 102(5), 918–928. https://doi.org/10.1016/j.neuron.2019.04.021

Diedrichsen, J., & Zotow, E. (2015). Surface-Based Display of Volume-Averaged Cerebellar Imaging Data. PloS One, 10(7), e0133402. https://doi.org/10.1371/journal.pone.0133402

D’Mello, A. M., Turkeltaub, P. E., & Stoodley, C. J. (2017). Cerebellar tDCS Modulates Neural Circuits during Semantic Prediction: A Combined tDCS-fMRI Study. The Journal of Neuroscience: The Official Journal of the Society for Neuroscience, 37(6), 1604–1613. https://doi.org/10.1523/JNEUROSCI.2818-16.2017

Eccles, J. C., Ito, M., & Szentágothai, J. (1967). Architectural Design of the Cerebellar Cortex. In J. C. Eccles, M. Ito, & J. Szentágothai (Eds.), The Cerebellum as a Neuronal Machine (pp. 195–204). Springer Berlin Heidelberg. https://doi.org/10.1007/978-3-662-13147-3_12

E, K.-H., Chen, S.-H. A., Ho, M.-H. R., & Desmond, J. E. (2014). A meta-analysis of cerebellar contributions to higher cognition from PET and fMRI studies. Human Brain Mapping, 35(2), 593–615. https://doi.org/10.1002/hbm.22194

Friston, K. J., Price, C. J., Fletcher, P., Moore, C., Frackowiak, R. S. J., & Dolan, R. J. (1996). The trouble with cognitive subtraction. NeuroImage, 4(2), 97–104. https://doi.org/10.1006/nimg.1996.0033

Galea, J. M., Vazquez, A., Pasricha, N., de Xivry, J.-J. O., & Celnik, P. (2011). Dissociating the roles of the cerebellum and motor cortex during adaptive learning: the motor cortex retains what the cerebellum learns. Cerebral Cortex, 21(8), 1761–1770. https://doi.org/10.1093/cercor/bhq246

Ganong, W. F., 3rd. (1980). Phonetic categorization in auditory word perception. Journal of Experimental Psychology. Human Perception and Performance, 6(1), 110–125. http://www.ncbi.nlm.nih.gov/pubmed/6444985

Gellersen, H. M., Guo, C. C., & O’Callaghan, C. (2017). Cerebellar atrophy in neurodegeneration—a meta-analysis. Journal of Neurology, Neurosurgery, and Psychiatry. https://jnnp.bmj.com/content/88/9/780.abstract?casa_token=WMQikIqm8nYAAAAA:AoWT85FpDf9xrP9eTJWw7yy8favpdm9Y2zpQLdGty-u0K9qYFq6MypgOKj4kT-k7PJ18AWi1-LTK

Gibo, T. L., Criscimagna-Hemminger, S. E., Okamura, A. M., & Bastian, A. J. (2013). Cerebellar motor learning: are environment dynamics more important than error size? Journal of Neurophysiology, 110(2), 322–333. https://doi.org/10.1152/jn.00745.2012

Glickstein, M., Strata, P., & Voogd, J. (2009). Cerebellum: history. Neuroscience, 162(3), 549–559. https://doi.org/10.1016/j.neuroscience.2009.02.054

Goldinger, S. D., & Azuma, T. (2003). Puzzle-solving science: the quixotic quest for units in speech perception. Journal of Phonetics, 31(3-4), 305–320. https://doi.org/10.1016/S0095-4470(03)00030-5

Grodd, W., Hülsmann, E., Lotze, M., Wildgruber, D., & Erb, M. (2001). Sensorimotor mapping of the human cerebellum: fMRI evidence of somatotopic organization. Human Brain Mapping, 13(2), 55–73. http://onlinelibrary.wiley.com/doi/10.1002/hbm.1025/full

Guediche, S., Holt, L. L., Laurent, P., Lim, S.-J., & Fiez, J. A. (2015). Evidence for Cerebellar Contributions to Adaptive Plasticity in Speech Perception. Cerebral Cortex, 25(7), 1867–1877. https://doi.org/10.1093/cercor/bht428

Guell, X., Gabrieli, J. D. E., & Schmahmann, J. D. (2018). Triple representation of language, working memory, social and emotion processing in the cerebellum: convergent evidence from task and seed-based resting-state fMRI analyses in a single large cohort. NeuroImage, 172, 437–449. https://doi.org/10.1016/j.neuroimage.2018.01.082

Guell, X., Schmahmann, J. D., Gabrieli, J. D. E., & Ghosh, S. S. (2018). Functional gradients of the cerebellum. eLife, 7. https://doi.org/10.7554/eLife.36652

Herculano-Houzel, S., Catania, K., Manger, P. R., & Kaas, J. H. (2015). Mammalian Brains Are Made of These: A Dataset of the Numbers and Densities of Neuronal and Nonneuronal Cells in the Brain of Glires, Primates, Scandentia, Eulipotyphlans, Afrotherians and Artiodactyls, and Their Relationship with Body Mass. Brain, Behavior and Evolution, 86(3-4), 145–163. https://doi.org/10.1159/000437413

Hertrich, I., Mathiak, K., & Ackermann, H. (2016). The role of the cerebellum in speech perception and language comprehension. In The linguistic cerebellum (pp. 33–50). Elsevier. https://www.sciencedirect.com/science/article/pii/B9780128016084000025

Holt, L. L., & Lotto, A. J. (2002). Behavioral examinations of the level of auditory processing of speech context effects. Hearing Research, 167(1-2), 156–169. https://doi.org/10.1016/s0378-5955(02)00383-0

Hurley, S. (2001). Perception And Action: Alternative Views. Synthese, 129(1), 3–40. https://doi.org/10.1023/A:1012643006930

Imamizu, H., Kuroda, T., Miyauchi, S., Yoshioka, T., & Kawato, M. (2003). Modular organization of internal models of tools in the human cerebellum. Proceedings of the National Academy of Sciences of the United States of America, 100(9), 5461–5466. https://doi.org/10.1073/pnas.0835746100

Ito, M. (2008). Control of mental activities by internal models in the cerebellum. Nature Reviews. Neuroscience, 9(4), 304–313. https://doi.org/10.1038/nrn2332

Ivry, R. B., & Gopal, H. S. (1993). Speech Production and Perception in Patients with Cerebellar Lesions. In Attention and Performance XIV. https://doi.org/10.7551/mitpress/1477.003.0045

Jalali, R., Miall, R. C., & Galea, J. M. (2017). No consistent effect of cerebellar transcranial direct current stimulation (tDCS) on visuomotor adaptation. Journal of Neurophysiology, jn.00896.2016. https://doi.org/10.1152/jn.00896.2016

Jayaram, G., Galea, J. M., Bastian, A. J., & Celnik, P. (2011). Human locomotor adaptive learning is proportional to depression of cerebellar excitability. Cerebral Cortex, 21(8), 1901–1909. https://doi.org/10.1093/cercor/bhq263

Johnson, J. F., Belyk, M., Schwartze, M., Pinheiro, A. P., & Kotz, S. A. (2019). The role of the cerebellum in adaptation: ALE meta-analyses on sensory feedback error. Human Brain Mapping, 40(13), 3966–3981. https://doi.org/10.1002/hbm.24681

Keller, G. B., & Mrsic-Flogel, T. D. (2018). Predictive Processing: A Canonical Cortical Computation. Neuron, 100(2), 424–435. https://doi.org/10.1016/j.neuron.2018.10.003

King, M., Hernandez-Castillo, C. R., Poldrack, R. A., Ivry, R. B., & Diedrichsen, J. (2019). Functional boundaries in the human cerebellum revealed by a multi-domain task battery. Nature Neuroscience, 22(8), 1371–1378. https://doi.org/10.1038/s41593-019-0436-x

Knolle, F., Schröger, E., Baess, P., & Kotz, S. A. (2012). The cerebellum generates motor-to-auditory predictions: ERP lesion evidence. Journal of Cognitive Neuroscience, 24(3), 698–706. http://www.mitpressjournals.org/doi/abs/10.1162/jocn_a_00167

Kotz, S. A., & Schwartze, M. (2010). Cortical speech processing unplugged: a timely subcortico-cortical framework. Trends in Cognitive Sciences, 14(9), 392–399. https://doi.org/10.1016/j.tics.2010.06.005

Kotz, S. A., Stockert, A., & Schwartze, M. (2014). Cerebellum, temporal predictability and the updating of a mental model. Philosophical Transactions of the Royal Society of London. Series B, Biological Sciences, 369(1658), 20130403. https://doi.org/10.1098/rstb.2013.0403

Kunert, R., & Slevc, L. R. (2015). A Commentary on: “Neural overlap in processing music and speech.” Frontiers in Human Neuroscience, 9, 330. https://doi.org/10.3389/fnhum.2015.00330

Ladefoged, P., & Broadbent, D. E. (1957). Information Conveyed by Vowels. The Journal of the Acoustical Society of America, 29(1), 98–104. https://doi.org/10.1121/1.1908694

Lametti, D. R., Oostwoud Wijdenes, L., Bonaiuto, J., Bestmann, S., & Rothwell, J. C. (2016). Cerebellar tDCS Dissociates the Timing of Perceptual Decisions from Perceptual Change in Speech. Journal of Neurophysiology, jn.00433.2016. https://doi.org/10.1152/jn.00433.2016

Lametti, D. R., Smith, H. J., Freidin, P., & Watkins, K. E. (2017). Cortico-cerebellar Networks Drive Sensorimotor Learning in Speech. Journal of Cognitive Neuroscience, 1–12. https://doi.org/10.1162/jocn_a_01216

Lametti, D. R., Smith, H. J., Watkins, K. E., & Shiller, D. M. (2018). Robust Sensorimotor Learning During Variable Sentence Level Speech. Current Biology: CB.

Lesage, E., Hansen, P. C., & Miall, R. C. (2017). Right Lateral Cerebellum Represents Linguistic Predictability. The Journal of Neuroscience: The Official Journal of the Society for Neuroscience, 37(26), 6231–6241. https://doi.org/10.1523/JNEUROSCI.3203-16.2017

Lesage, E., Morgan, B. E., Olson, A. C., Meyer, A. S., & Miall, R. C. (2012). Cerebellar rTMS disrupts predictive language processing. Current Biology: CB, 22(18), R794–R795. https://doi.org/10.1016/j.cub.2012.07.006

Lotto, A. J., & Holt, L. (2000). The illusion of the phoneme. https://kilthub.cmu.edu/articles/The_Illusion_of_the_Phoneme/6618578/1

Macklis, R. M., & Macklis, J. D. (1992). Historical and phrenologic reflections on the nonmotor functions of the cerebellum: love under the tent? Neurology, 42(4), 928–932. https://doi.org/10.1212/wnl.42.4.928

MacLeod, C. E., Zilles, K., Schleicher, A., Rilling, J. K., & Gibson, K. R. (2003). Expansion of the neocerebellum in Hominoidea. Journal of Human Evolution, 44(4), 401–429. https://doi.org/10.1016/s0047-2484(03)00028-9

Marek, S., Siegel, J. S., Gordon, E. M., Raut, R. V., Gratton, C., Newbold, D. J., Ortega, M., Laumann, T. O., Adeyemo, B., Miller, D. B., Zheng, A., Lopez, K. C., Berg, J. J., Coalson, R. S., Nguyen, A. L., Dierker, D., Van, A. N., Hoyt, C. R., McDermott, K. B., … Dosenbach, N. U. F. (2018). Spatial and Temporal Organization of the Individual Human Cerebellum. Neuron, 100(4), 977–993.e7. https://doi.org/10.1016/j.neuron.2018.10.010

Mariën, P., Ackermann, H., Adamaszek, M., Barwood, C. H. S., Beaton, A., Desmond, J., De Witte, E., Fawcett, A. J., Hertrich, I., Küper, M., Leggio, M., Marvel, C., Molinari, M., Murdoch, B. E., Nicolson, R. I., Schmahmann, J. D., Stoodley, C. J., Thürling, M., Timmann, D., … Ziegler, W. (2014). Consensus paper: Language and the cerebellum: an ongoing enigma. Cerebellum, 13(3), 386–410. https://doi.org/10.1007/s12311-013-0540-5

Mariën, P., & Borgatti, R. (2018). Chapter 11 - Language and the cerebellum. In M. Manto & T. A. G. M. Huisman (Eds.), Handbook of Clinical Neurology (Vol. 154, pp. 181–202). Elsevier. https://doi.org/10.1016/B978-0-444-63956-1.00011-4

Mariën, P., & Manto, M. (2015). The Linguistic Cerebellum. Academic Press. https://play.google.com/store/books/details?id=aQGdBAAAQBAJ

Martin, T. A., Keating, J. G., Goodkin, H. P., Bastian, A. J., & Thach, W. T. (1996). Throwing while looking through prisms. I. Focal olivocerebellar lesions impair adaptation. Brain: A Journal of Neurology, 119 (*Pt 4*), 1183–1198. https://www.ncbi.nlm.nih.gov/pubmed/8813282

Mathiak, K., Hertrich, I., Grodd, W., & Ackermann, H. (2002). Cerebellum and speech perception: a functional magnetic resonance imaging study. Journal of Cognitive Neuroscience, 14(6), 902–912. https://doi.org/10.1162/089892902760191126

McGurk, H., & MacDonald, J. (1976). Hearing lips and seeing voices. Nature, 264(5588), 746–748. http://www.ncbi.nlm.nih.gov/pubmed/1012311

Moberget, T., Gullesen, E. H., Andersson, S., Ivry, R. B., & Endestad, T. (2014). Generalized role for the cerebellum in encoding internal models: evidence from semantic processing. The Journal of Neuroscience: The Official Journal of the Society for Neuroscience, 34(8), 2871–2878. https://doi.org/10.1523/JNEUROSCI.2264-13.2014

Moberget, T., & Ivry, R. B. (2016). Cerebellar contributions to motor control and language comprehension: searching for common computational principles. Annals of the New York Academy of Sciences, 1369(1), 154–171. https://doi.org/10.1111/nyas.13094

Moberget, T., & Ivry, R. B. (2019). Prediction, Psychosis, and the Cerebellum. Biological Psychiatry. Cognitive Neuroscience and Neuroimaging, 4(9), 820–831. https://doi.org/10.1016/j.bpsc.2019.06.001

Morton, S. M., & Bastian, A. J. (2004). Prism adaptation during walking generalizes to reaching and requires the cerebellum. Journal of Neurophysiology, 92(4), 2497–2509. https://doi.org/10.1152/jn.00129.2004

Parrell, B., Agnew, Z., Nagarajan, S., Houde, J., & Ivry, R. B. (2017). Impaired Feedforward Control and Enhanced Feedback Control of Speech in Patients with Cerebellar Degeneration. The Journal of Neuroscience: The Official Journal of the Society for Neuroscience, 37(38), 9249–9258. https://doi.org/10.1523/JNEUROSCI.3363-16.2017

Peretz, I., Vuvan, D., Lagrois, M.-É., & Armony, J. L. (2015). Neural overlap in processing music and speech. Philosophical Transactions of the Royal Society of London. Series B, Biological Sciences, 370(1664), 20140090. https://doi.org/10.1098/rstb.2014.0090

Petacchi, A., Laird, A. R., Fox, P. T., & Bower, J. M. (2005). Cerebellum and auditory function: an ALE meta-analysis of functional neuroimaging studies. Human Brain Mapping, 25(1), 118–128. https://doi.org/10.1002/hbm.20137

Pleger, B., & Timmann, D. (2018). The role of the human cerebellum in linguistic prediction, word generation and verbal working memory: evidence from brain imaging, non-invasive cerebellar stimulation and lesion studies. Neuropsychologia, 115, 204–210. https://doi.org/10.1016/j.neuropsychologia.2018.03.012

Poeppel, D. (1996). A critical review of PET studies of phonological processing. Brain and Language, 55(3), 317–351; discussion 352–385. https://doi.org/10.1006/brln.1996.0108

Popa, L. S., & Ebner, T. J. (2018). Cerebellum, Predictions and Errors. Frontiers in Cellular Neuroscience, 12, 524. https://doi.org/10.3389/fncel.2018.00524

Rabe, K., Livne, O., Gizewski, E. R., Aurich, V., Beck, A., Timmann, D., & Donchin, O. (2009). Adaptation to visuomotor rotation and force field perturbation is correlated to different brain areas in patients with cerebellar degeneration. Journal of Neurophysiology, 101(4), 1961–1971. https://doi.org/10.1152/jn.91069.2008

Repp, B. H. (1979). Relative amplitude of aspiration noise as a voicing cue for syllable-initial stop consonants. Language and Speech, 22(2), 173–189. https://doi.org/10.1177/002383097902200207

Riedel, M. C., Ray, K. L., Dick, A. S., Sutherland, M. T., Hernandez, Z., Fox, P. M., Eickhoff, S. B., Fox, P. T., & Laird, A. R. (2015). Meta-analytic connectivity and behavioral parcellation of the human cerebellum. NeuroImage, 117, 327–342. https://doi.org/10.1016/j.neuroimage.2015.05.008

Schmahmann, J. D. (2019). The cerebellum and cognition. Neuroscience Letters, 688, 62–75. https://doi.org/10.1016/j.neulet.2018.07.005

Schmahmann, J. D., Guell, X., Stoodley, C. J., & Halko, M. A. (2019). The Theory and Neuroscience of Cerebellar Cognition. Annual Review of Neuroscience, 42, 337–364. https://doi.org/10.1146/annurev-neuro-070918-050258

Schwartze, M., Keller, P. E., & Kotz, S. A. (2016). Spontaneous, synchronized, and corrective timing behavior in cerebellar lesion patients. Behavioural Brain Research, 312, 285–293. https://doi.org/10.1016/j.bbr.2016.06.040

Schwartze, M., & Kotz, S. A. (2013). A dual-pathway neural architecture for specific temporal prediction. Neuroscience and Biobehavioral Reviews, *37*(10 Pt 2), 2587–2596. https://doi.org/10.1016/j.neubiorev.2013.08.005

Schwartze, M., & Kotz, S. A. (2016). Contributions of cerebellar event-based temporal processing and preparatory function to speech perception. Brain and Language, 161, 28–32. https://doi.org/10.1016/j.bandl.2015.08.005

Schwartze, M., Tavano, A., Schröger, E., & Kotz, S. A. (2012). Temporal aspects of prediction in audition: Cortical and subcortical neural mechanisms. International Journal of Psychophysiology: Official Journal of the International Organization of Psychophysiology, 83(2), 200–207. https://doi.org/10.1016/j.ijpsycho.2011.11.003

Shadmehr, R., Smith, M. A., & Krakauer, J. W. (2010). Error correction, sensory prediction, and adaptation in motor control. Annual Review of Neuroscience, 33, 89–108. https://doi.org/10.1146/annurev-neuro-060909-153135

Sheu, Y.-S., Liang, Y., & Desmond, J. E. (2019). Disruption of Cerebellar Prediction in Verbal Working Memory. Frontiers in Human Neuroscience, 13, 61. https://doi.org/10.3389/fnhum.2019.00061

Siman-Tov, T., Granot, R. Y., Shany, O., Singer, N., Hendler, T., & Gordon, C. R. (2019). Is there a prediction network? Meta-analytic evidence for a cortical-subcortical network likely subserving prediction. Neuroscience and Biobehavioral Reviews, 105, 262–275. https://doi.org/10.1016/j.neubiorev.2019.08.012

Sjerps, M. J., Mitterer, H., & McQueen, J. M. (2011). Listening to different speakers: on the time-course of perceptual compensation for vocal-tract characteristics. Neuropsychologia, 49(14), 3831–3846. https://doi.org/10.1016/j.neuropsychologia.2011.09.044

Skipper, J. I. (2014). Echoes of the spoken past: how auditory cortex hears context during speech perception. Philosophical Transactions of the Royal Society of London. Series B, Biological Sciences, 369(1651), 20130297. https://doi.org/10.1098/rstb.2013.0297

Skipper, J. I. (2015). The NOLB model: a model of the natural organization of language and the brain. In R. M. Willems & R. M. Willems (Eds.), Cognitive Neuroscience of Natural Language Use (pp. 101–134). Cambridge University Press. https://doi.org/10.1017/CBO9781107323667.006

Skipper, J. I., Devlin, J. T., & Lametti, D. R. (2017). The hearing ear is always found close to the speaking tongue: Review of the role of the motor system in speech perception. Brain and Language, 164, 77–105. https://doi.org/10.1016/j.bandl.2016.10.004

Skipper, J. I., & Hasson, U. (2017). A Core Speech Circuit Between Primary Motor, Somatosensory, And Auditory Cortex: Evidence From Connectivity And Genetic Descriptions. In bioRxiv (p. 139550). https://doi.org/10.1101/139550

Skipper, J. I., Nusbaum, H. C., & Small, S. L. (2005). Listening to talking faces: motor cortical activation during speech perception. NeuroImage, 25(1), 76–89. https://doi.org/10.1016/j.neuroimage.2004.11.006

Skipper, J. I., van Wassenhove, V., Nusbaum, H. C., & Small, S. L. (2007). Hearing lips and seeing voices: how cortical areas supporting speech production mediate audiovisual speech perception. Cerebral Cortex, 17(10), 2387–2399. https://doi.org/10.1093/cercor/bhl147

Smaers, J. B., Turner, A. H., Gómez-Robles, A., & Sherwood, C. C. (2018). A cerebellar substrate for cognition evolved multiple times independently in mammals. eLife, 7. https://doi.org/10.7554/eLife.35696

Smith, M. A., & Shadmehr, R. (2005). Intact ability to learn internal models of arm dynamics in Huntington’s disease but not cerebellar degeneration. Journal of Neurophysiology, 93(5), 2809–2821. https://doi.org/10.1152/jn.00943.2004

Stoodley, C. J., & Schmahmann, J. D. (2009). Functional topography in the human cerebellum: a meta-analysis of neuroimaging studies. NeuroImage, 44(2), 489–501. https://doi.org/10.1016/j.neuroimage.2008.08.039

Stoodley, C. J., & Schmahmann, J. D. (2018). Chapter 4 - Functional topography of the human cerebellum. In M. Manto & T. A. G. M. Huisman (Eds.), Handbook of Clinical Neurology (Vol. 154, pp. 59–70). Elsevier. https://doi.org/10.1016/B978-0-444-63956-1.00004-7

Stoodley, C. J., Valera, E. M., & Schmahmann, J. D. (2012). Functional topography of the cerebellum for motor and cognitive tasks: an fMRI study. NeuroImage, 59(2), 1560–1570. https://doi.org/10.1016/j.neuroimage.2011.08.065

Taylor, J. A., & Ivry, R. B. (2014). Cerebellar and prefrontal cortex contributions to adaptation, strategies, and reinforcement learning. Progress in Brain Research, 210, 217–253. https://doi.org/10.1016/B978-0-444-63356-9.00009-1

Tourville, J. A., Reilly, K. J., & Guenther, F. H. (2008). Neural mechanisms underlying auditory feedback control of speech. NeuroImage, 39(3), 1429–1443. https://doi.org/10.1016/j.neuroimage.2007.09.054

Van Overwalle, F., Baetens, K., Mariën, P., & Vandekerckhove, M. (2014). Social cognition and the cerebellum: a meta-analysis of over 350 fMRI studies. NeuroImage, 86, 554–572. https://doi.org/10.1016/j.neuroimage.2013.09.033

Wilson, S. M. (2009). Speech perception when the motor system is compromised [Review of *Speech perception when the motor system is compromised*]. Trends in Cognitive Sciences, 13(8), 329–330; author reply 330–331. https://doi.org/10.1016/j.tics.2009.06.001

Wilson, S. M., & Iacoboni, M. (2006). Neural responses to non-native phonemes varying in producibility: evidence for the sensorimotor nature of speech perception. NeuroImage, 33(1), 316–325. https://doi.org/10.1016/j.neuroimage.2006.05.032

Wolpert, D. M., Diedrichsen, J., & Flanagan, J. R. (2011). Principles of sensorimotor learning. Nature Reviews. Neuroscience, 12(12), 739–751. https://doi.org/10.1038/nrn3112

Wolpert, D. M., Miall, R. C., & Kawato, M. (1998). Internal models in the cerebellum. Trends in Cognitive Sciences, 2(9), 338–347. http://www.ncbi.nlm.nih.gov/pubmed/21227230

Yarkoni, T., Poldrack, R. A., Nichols, T. E., Van Essen, D. C., & Wager, T. D. (2011). Large-scale automated synthesis of human functional neuroimaging data. Nature Methods, 8(8), 665–670. https://doi.org/10.1038/nmeth.1635

Yip, M. (2002). Tone. Cambridge University Press. https://play.google.com/store/books/details?id=KFv2lojXjpwC

